# HOPS-dependent endosomal fusion required for efficient cytosolic delivery of therapeutic peptides and small proteins

**DOI:** 10.1101/374926

**Authors:** Angela Steinauer, Jonathan R. LaRochelle, Rebecca Wissner, Samuel Berry, Alanna Schepartz

## Abstract

Protein therapeutics represent a significant and growing component of the modern pharmacopeia, but their potential to treat human disease is limited because most proteins fail to traffic across biological membranes. Recently, we discovered that cell-permeant miniature proteins (CPMPs) containing a precisely defined, penta-arginine motif traffic readily to the cytosol and nucleus with efficiencies that rival those of hydrocarbon-stapled peptides active in animals and man. Like many cell-penetrating peptides (CPPs), CPMPs enter the endocytic pathway; the difference is that CPMPs are released efficiently from endosomes while other CPPs are not. Here, we seek to understand how CPMPs traffic from endosomes into the cytosol and what factors contribute to the efficiency of endosomal release. First, using two complementary cell-based assays, we exclude endosomal rupture as the primary means of endosomal escape. Next, using a broad spectrum of techniques, including an RNA interference (RNAi) screen, fluorescence correlation spectroscopy (FCS), and confocal imaging, we identify VPS39—a gene encoding a subunit of the homotypic fusion and protein sorting (HOPS) complex—as a critical determinant in the trafficking of CPMPs and hydrocarbon-stapled peptides to the cytosol. Although CPMPs neither inhibit nor activate HOPS function, HOPS activity is essential to efficiently deliver CPMPs to the cytosol. Subsequent multi-color confocal imaging studies identify CPMPs within the endosomal lumen, particularly within the intraluminal vesicles (ILVs) of Rab7^+^ and Lamp1^+^ endosomes that are the products of HOPS-mediated fusion. These results suggest that CPMPs require HOPS to reach ILVs—an environment that serves as a prerequisite for efficient endosomal escape.

## INTRODUCTION

Protein and peptide therapeutics—biologics—comprise a rapidly growing sector of the modern pharmacopeia (1). Seven of the top ten highest grossing therapeutic agents in 2017 are biologics used to treat cancer (2, 3), diabetes (4), and autoimmune inflammatory disorders such as rheumatoid arthritis and Crohn’s disease (5-7). In each case, the biologic acts by stimulating, inhibiting, or replacing a protein located within plasma or on an external membrane surface (1). Not one acts within the cell cytosol or nucleus, in large part because most proteins cannot effectively breach the barrier defined by the plasma membrane (8, 9). The well-known early exceptions to this rule discovered by Löwenstein (10), Pabo (11), and Derossi (12)—the HIV trans-activator of transcription (Tat) protein and the Antennapedia homeodomain—have inspired the synthesis, study, and (in some cases) clinical evaluation (13) of literally hundreds of arginine-rich “cell-penetrating peptides” (CPPs) (14). The problem is that when added to cells, most CPPs remain trapped in endosomes and fail to achieve significant concentrations in the cytosol or nucleus (15). The inefficient delivery of proteins, peptides, and their mimetics into the mammalian cell cytosol limits their potential as therapeutics and research tools.

Recently we discovered that, when added to cells, certain small, folded miniature proteins (16, 17) derived from avian pancreatic polypeptide (aPP) or an isolated zinc-finger (ZF) domain are taken up by the endocytic pathway and released into the cytosol with unprecedented efficiencies (18, 19). The most effective molecules are defined by a discrete array of five arginine residues on a folded α-helix (20); we refer to these molecules as cell-permeant miniature proteins (CPMPs). Treatment of HeLa cells in culture with the CPMP ZF5.3 leads to a ZF5.3 concentration in the cytosol that is roughly 67% of the extracellular incubation concentration; this value is at least ten-fold higher than that achieved by HIV-Tat_48-60_ peptide (21) or octaarginine (Arg8) and equal to that of hydrocarbon-stapled peptides under development as protein-protein interaction inhibitors (22). Comparable improvements in cytosolic access are observed when the CPMP ZF5.3 is fused to protein cargos with significant molecular mass (23).

Here, we describe experiments that seek to understand how CPMPs like ZF5.3 traffic from endosomes into the cytosol and what factors contribute to the efficiency of endosomal release. First, using two complementary cell-based assays, we exclude endosomal rupture as the primary means of endosomal escape. Next, using a broad spectrum of techniques, including an RNA interference (RNAi) screen, fluorescence correlation spectroscopy (FCS), genetic knockdowns, and confocal imaging, we identify VPS39—a gene encoding a subunit of the homotypic fusion and protein sorting (HOPS) complex—as a critical determinant in the trafficking of CPMPs and hydrocarbon-stapled peptides to the cytosol. HOPS activity is essential for cytosolic access; the closely related class C core vacuole/endosome tethering (CORVET) complex is not required. CPMPs neither inhibit nor activate HOPS activity, and we find no evidence for a direct HOPS-CPMP interaction. Subsequent multi-color confocal imaging studies identify CPMPs within the endosomal lumen, particularly within the intraluminal vesicles (ILVs) of Rab7^+^ and Lamp1^+^ endosomes that are the products of HOPS-mediated fusion (24). We conclude that HOPS allows CPMPs to traffic into intraluminal vesicles (ILVs), a favorable environment for endosomal escape. The identification of ILVs as a portal for passing proteins into the cytosol will aid the development of next-generation biologics that overcome the limitations imposed by cellular membranes.

## RESULTS

### Evaluating endosomal damage

The simplest way for a CPMP to escape from an endosome is if the endosome ruptures, in part or in full (25). Although there has been limited work on the effects of certain CPPs on the integrity of large unilamellar vesicles (LUVs) *in vitro* (26), the concentration-dependent effects of CPMPs or more traditional cell-penetrating peptides (CPPs) on endosomal integrity in cultured cells have not been thoroughly evaluated. Thus, we began our analysis with two complementary assays that together detect both subtle and severe endosomal damage in cells treated with a CPMP or CPP. One assay exploits a set of eGFP-labeled galectins to fluorescently tag damaged endosomes to enable their visualization using confocal microscopy, while the other employs a fluorescently tagged version of the nonalysine (Lys9) peptide to quantify the extent of endosome rupture in cells treated with a CPMP or CPP. In both cases, the effects of the two most efficient CPMPs—aPP5.3 (**1**) and ZF5.3 (**2**)—were compared with those of prototypic members of three CPP families: the hydrocarbon-stapled peptide SAH-p53-8 (**3**) (22); D-octaarginine (D-Arg8, **4**) (27), and a cyclic peptide containing both natural and unnatural amino acids, CPP12 (**5**) (26) (Figure 1 and Figure S1). SAH-p53-8 (**3**) is a hydrocarbon-stapled peptide that reaches the cell interior despite the absence of excess positive charge, D-octaarginine is the proteolytically stable enantiomer of the widely studied octapeptide L-Arg8 (27, 28), and CPP12 (**5**) is a cyclic peptide that reportedly reaches the cytosol with an efficiency that rivals aPP5.3 (26).

**Figure 1:**
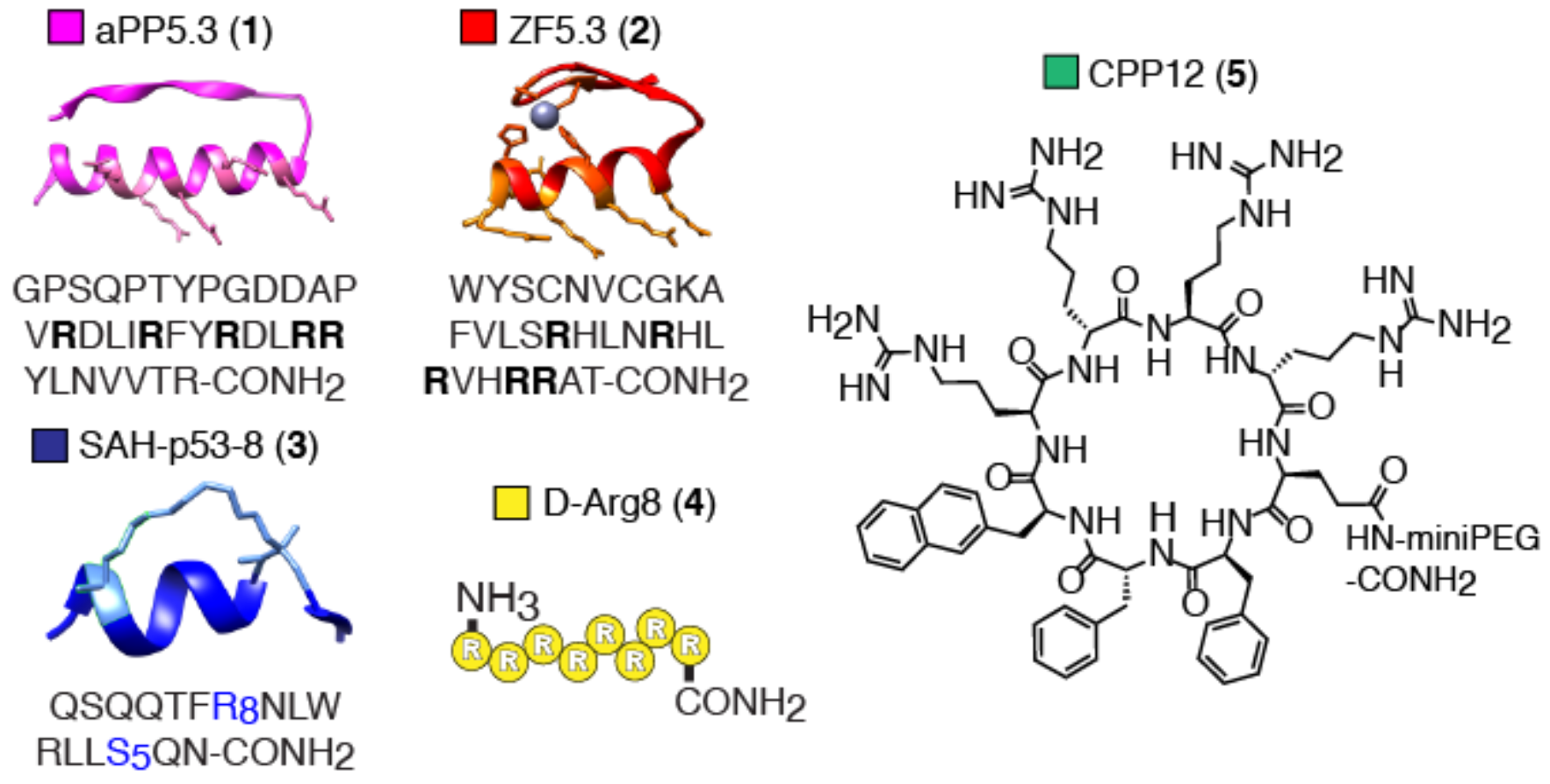
CPMPs and CPPs evaluated herein include aPP5.3. (**1**) (model structure from PDB: 1PPT); ZF5.3 (**2**) (model structure from PDB: 2EOZ); SAH-p53-8 (**3**) (model structure from PDB: 3V3B); D-Arg8 (**4**); and the cyclic peptide CPP12 (**5**). Unnatural amino acids: R_8_ = *R*-octenylalanine, S_5_ = *S*-pentenylalanine. See also Figure S1 and Table S1.

### CPMPs and CPPs do not induce galectin recruitment at sub-micromolar concentrations

A characteristic feature of endosomes that have been damaged by invading viruses (29, 30), bacteria-triggered autophagy (31-34), or endosomolytic nanoparticles (35) is the cytosolic display of β-galactosides linked to proteoglycans that are ordinarily found on the luminal side of endolysosomal compartments (36). Once displayed to the cytosol, β-galactosides recruit cytosolic galectin (Gal) proteins (37, 38), such as Gal1, 3, 4, 8, and 9, that share a conserved β-galactoside binding site (Figure S2A) (39, 40). In particular, Gal3 and Gal8 are recruited to damaged Rab7^+^ and Lamp1^+^ endosomes that form along the degradative branch of the endocytic pathway (33, 40). Previous work has shown that endosomal damage can be detected by monitoring the translocation of eGFP fusions of Gal3 or Gal8 from the cytosol to endosome surfaces (29-34). Fusions of Gal3 or Gal8 and eGFP have also been used to visualize late endosome damage upon siRNA release from endosomes (40), screen for small molecules that induce lysosomal rupture (41), visualize endosomal leakage induced by osmocytosis and propanebetaine (iTOP) (42), and evaluate cell-permeant peptide-polymer complexes (35). However, the effects of CPPs or CPMPs on the extent of galectin-recruitment to damaged endosomes have not previously been studied.

To evaluate whether CPMPs such as aPP5.3 (**1**) and ZF5.3 (**2**) lead to the recruitment of galectins to endosomal compartments, we made use of eGFP fusions of human galectin-3 (eGFP-hGal3) and human galectin-8 (eGFP-hGal8), both of which have been used previously to detect endosomal damage (31, 39, 40). Human osteosarcoma (Saos-2) cells transiently expressing eGFP-hGal3 or eGFP-hGal8 were first treated for 1 hour with two known endosomolytic agents at concentrations reported to induce endosomal rupture and imaged using confocal microscopy to assess galectin recruitment from the cytosol to damaged endosomes (Figure S2B). These positive control reagents included Lipofectamine RNAiMAX (referred to as RNAiMAX henceforth) (16 μL/mL) and L-leucyl-L-leucine methyl ester (LLOMe) (1 mM). RNAiMAX is a cationic lipid formulation known to induce endosomal rupture to deliver siRNAs for targeted gene knockdown (40, 43). To induce visually detectable levels of endosomal rupture, we chose a concentration that is 3-fold higher than the average recommended concentration for siRNA transfection. For LLOMe, we chose a concentration, which had previously been reported to induce the desired galectin recruitment phenotype (31). Even though we routinely evaluate CPMP delivery after a 30-minute incubation period, we chose to evaluate endosomal rupture after a 60-minute incubation period to ensure detection of even low levels of rupture and galectin recruitment. The amount of endosomal recruitment was quantified using ImageJ (44) by calculating an endosomal recruitment coefficient (ERC), which was defined as the %-area of punctate fluorescence observed in a single cell divided by the total cell area, multiplied by 100 (Figure S2C). As expected (36, 40, 45), expression of eGFP-hGal3 or eGFP-hGal8 in untreated Saos-2 cells led to uniform eGFP fluorescence throughout the cytosol and nucleus with negligible punctate staining (Figure S2B). The average ERC values calculated in untreated Saos-2 cells expressing eGFP-hGal3 or eGFP-hGal8 were 2.0 ± 0.6 and 3.4 ± 0.7, respectively (Figure S2D). By contrast, treatment of Saos-2 cells with either RNAiMAX or LLOMe, both of which stimulate Gal3 and Gal8 recruitment to endolysosomal membranes in multiple cell lines (31, 33, 39, 40), led to significant punctate staining (Figure S2B). As previously reported, galectin recruitment occured within minutes and persisted for several hours (39). The average ERC values calculated when Saos-2 cells were treated with RNAiMAX were 62 ± 8 (Gal-3) and 42 ± 5 (Gal-8), values that represent increases of 30- and 14-fold over untreated cells (Figure S2B and S2D). The effects of LLOMe were even more dramatic: the average ERC values in Saos-2 cells expressing eGFP-hGal3 or eGFP-hGal8 treated with LLOMe increased 147- and 68-fold compared to untreated cells, respectively. These data confirm the utility of eGFP-hGal3 and eGFP-hGal8 for monitoring CPP or CPMP-induced endolysosomal damage in Saos-2 cells.

We next examined the effects of CPMPs aPP5.3 (**1**) and ZF5.3 (**2**) on the endosomal recruitment of eGFP-hGal3 and eGFP-hGal8. Side-by-side experiments were performed with SAH-p53-8 (**3**), D-Arg8 (**4**), and CPP12 (**5**). Each CPMP or CPP was tagged with a lissamine rhodamine B fluorophore (R) to enable its selective visualization alongside the eGFP-fused galectins. Galectin-expressing Saos-2 cells were treated with 600 nM **1**^R^–**5**^R^ (60 min), washed, and incubated for 30 min in CPMP/CPP-free media prior to imaging with confocal microscopy. This analysis revealed punctate red fluorescence throughout the cytosol, indicating endocytic uptake of CPPs **1**^R^–**5**^R^, and a uniform distribution of eGFP-hGal3 and −8 throughout the cytosol and nucleus (Figure S2B). The average ERC values calculated when Saos-2 cells expressing eGFP-hGal3 or eGFP-hGal8 were treated with 600 nM CPPs **1**^R^–**5**^R^ were all less than 15 (Figure S2D), which is not significantly above the ERC of untreated Saos-2 cells. These data suggest that **1**^R^–**5**^R^ cause little or no galectin-positive endosomal damage at a treatment concentration of 600 nM.

### CPMPs and CPPs do not induce endosomal leakage at concentrations below 2 μM

Although monitoring galectin recruitment reports on endolysosomal damage characterized by the cytosolic display of β-galactosides, it does not necessarily capture transient damage that can result in partial or complete endosomal leakage. To explore whether treatment of cells with CPMPs or CPPs causes endosomal leakage, we developed an assay to detect the release of lissamine rhodamine B-tagged Lys9 (Lys9^R^) from endosomes into the cytosol in the presence of CPPs or CPMPs. The nonapeptide Lys9 is internalized via endocytosis but not released into the cytosol under normal physiological conditions (46, 47). We verified this finding, and then used fluorescence correlation spectroscopy (FCS) (19) to quantify the concentration of Lys9^R^ that reached the cytosol in the presence of CPMPs or CPPs (Figure S3A). Control experiments confirmed that Saos-2 cells treated with increasing concentrations of RNAiMAX (0 to 16 μL/mL) exhibited a dose-dependent increase in Lys9^R^, which was detected in the cytosol and nucleus using FCS. The highest concentration of RNAiMAX (16 μL/mL) yielded a 39-fold increase in the cytosolic (806 ± 87 nM) and a 46-fold increase in the nuclear (1212 ± 129 nM) concentrations of Lys9^R^ (Figure S3B). Similar titration experiments with LLOMe yielded no detectable endosomal leakage of Lys9^R^, even at the highest tested LLOMe concentration (1 mM) (Figure S3C). LLOMe selectively induces lysosomal permeabilization (48, 49); it is possible that Lys9^R^ leakage can only occur efficiently from endosomes that precede the lysosome—an observation that has also been made for the endosomal escape of siRNA lipoplexes (40) and a disulfide-bonded dimer of Tat (dfTat) (50).

Additional control experiments using FCS confirmed that the cytosolic access of each CPMP or CPP was similar in the presence and absence of unlabeled Lys9 (Lys9^UL^) (Figure S4A). As reported previously for experiments performed in HeLa cells, CPMPs **1**^R^ and **2**^R^ and the hydrocarbon-stapled peptide **3**^R^ reached the cytosol and nucleus with significantly greater efficiency than CPPs **4**^R^ and **5**^R^, although cytosolic delivery was generally about 2-fold lower in Saos-2 cells than in HeLa cells (19). Miniature proteins **1**^R^ and **2**^R^, and the hydrocarbon-stapled peptide **3**^R^ were present in the cytosol at 3- to 9-fold higher concentrations than D-Arg8 **4**^R^. We were unable to perform conclusive FCS experiments with CPPs **4**^R^ and **5**^R^. Specifically, in cells treated with **4**^R^ and **5**^R^, we observed diffusion times that were between 50- to 150-fold longer compared to their respective *in vitro* diffusion times (Figure S4B and Table S2). Diffusion times in cultured cells reportedly increase between 2- and 10-fold compared to their corresponding *in vitro* diffusion times (51). Increases greater than 10-fold may indicate intracellular aggregation or binding to a cellular component, preventing us from extracting meaningful parameters from the FCS data. To circumvent this problem, we performed biochemical fractionation experiments (19) to estimate the cytosolic concentrations attained by **4**^R^ and **5**^R^.

After confirming that the presence of Lys9 had no significant effect on the ability of CPMP/CPPs **1**^R^–**5**^R^ to reach the cytosol, we evaluated the extent to which Lys9^R^ leaked from endosomes into the cytosol or nucleus after treatment with an unlabeled CPMP or CPP (**1**^UL^–**5**^UL^) at concentrations between 0.3 and 2.4 μM. No significant leakage of Lys9^R^ into the cytosol or nucleus was observed at CPMP or CPP concentrations below 2 *μ*M (Figure S3D–H). At 2.4 μM, CPMPs **1**^UL^ and **2**^UL^ induced low levels of Lys9^R^ endosomal leakage (Figure S3D and S3E), corresponding to a 6- and 10-fold increase above background for **1**^UL^ and **2**^UL^, respectively. A similar increase in intracellular Lys9^R^ concentration was observed at 8 μL/mL treatment with RNAiMAX (Figure S3B), a concentration that is commonly used for efficient siRNA transfection (43). The remaining polypeptides **3**^UL^–**5**^UL^ did not induce significant Lys9^R^ leakage at any concentration tested in this assay (Figure S3F–H). Taken together, the galectin recruitment and Lys9 leakage assays demonstrate that CPMPs and CPPs **1**^R^–**5**^R^/**1**^UL^–**5**^UL^ do not induce endosomal rupture at concentrations below 2 μM. Above 2 μM, **1**^UL^ and **2**^UL^ induce low levels of galectin recruitment and Lys9^R^ leakage, indicating that they are causing endosomal damage. Toxicity studies that monitored metabolic activity over an extended 4 h timeframe by detecting cellular ATP (CellTiter-Glo) revealed EC_50_ values for **1**^UL^, **2**^UL^**, 4**^UL^, **5**^UL^ that were >20 *μ*M; the value for SAH-p53-8, **3**^UL^, was 4.0 *μ*M (Figure S4C).

### Design of a genome-wide RNAi screen to probe endosomal release

The observation that sub-micromolar concentrations of CPMPs aPP5.3 (**1**) and ZF5.3 (**2**) can efficiently access the cytosol without significantly perturbing endosomal membranes raised the possibility that CPMP release from endosomes exploits a distinct cellular mechanism. To characterize this mechanism, we designed a genome-wide RNAi screen to identify candidate genes whose knockdown increase or decrease the ability of CPMP **1** to reach cytosol (Figure 2A). To quantify cytosolic delivery of **1**, we made use of a previously reported glucocorticoid-induced eGFP translocation (GIGT) assay (18, 20, 52) that couples the cytosolic delivery of a molecule tagged with dexamethasone (Dex) to the nuclear translocation of a reporter protein consisting of a glucocorticoid receptor variant with exceptional affinity for dexamethasone (GR*) (53, 54) fused to eGFP (GR*-eGFP) (Figure S5A). The effect of the Dex-tagged molecule on the ratio of GFP fluorescence in the nucleus and cytosol, termed the translocation ratio (TR), can be measured with precision using an Opera High-Content Screening System and thereby provides a quantifiable readout of cytosolic access (18, 20). Previously, we demonstrated that the GIGT/Opera combination assay is associated with robust Z-factors of ≥0.5 and is therefore suitable for quantifying the cytosolic delivery of Dex-labeled CPMPs aPP5.3^Dex^ (**1**^Dex^) and ZF5.3^Dex^ (**2**^Dex^) in a high-throughput setting (20). Saos-2 cells stably expressing GR*-GFP (Saos-2(GIGT) cells) were prepared as described previously (20).

**Figure 2:**
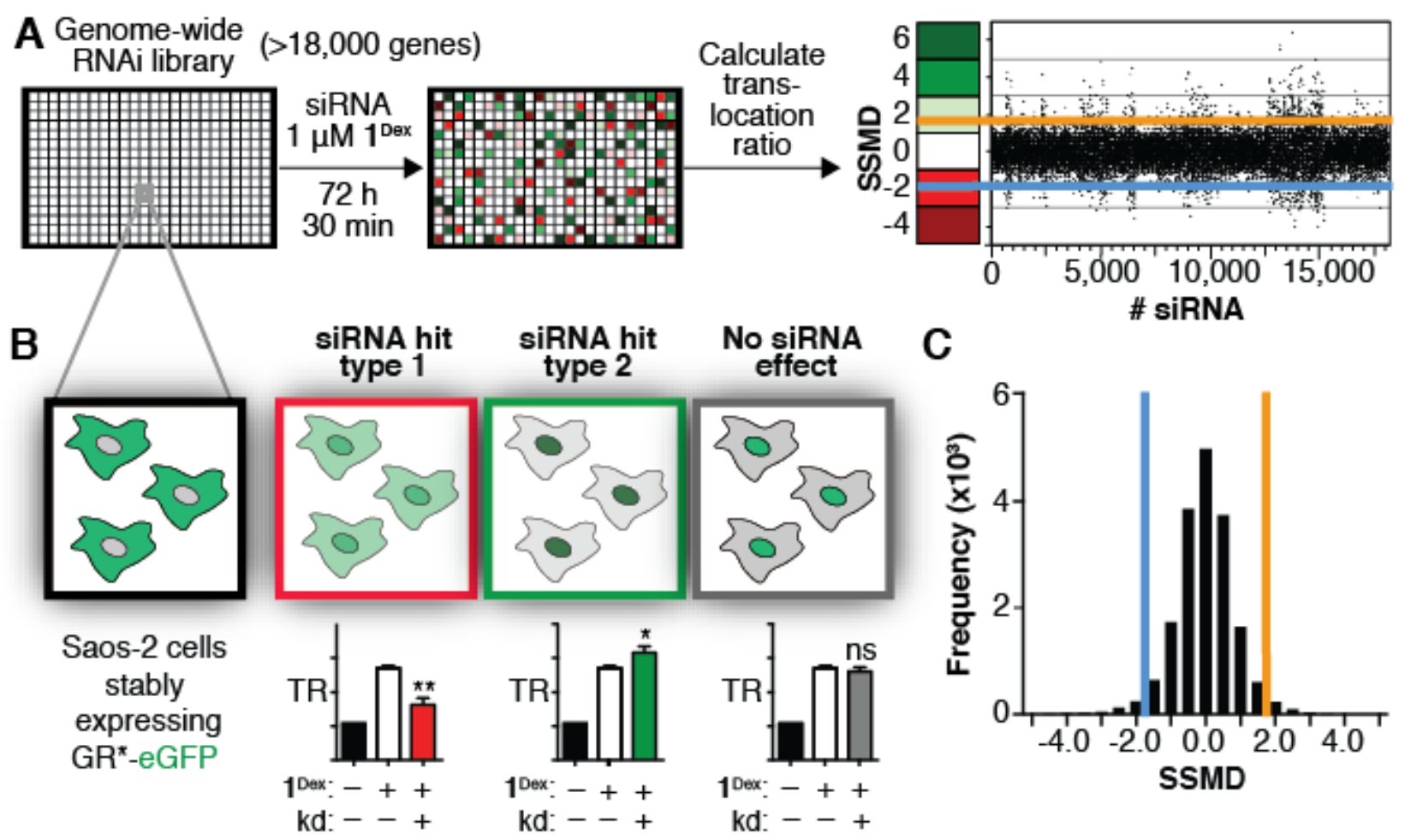
Genome-wide high content RNAi screen identifies genes involved in CPMP trafficking to cytosol. (A) We screened a Dharmacon SMARTpool human siRNA libraries (>18,000 genes) in Saos-2 cells stably expressing GR(C638G)-eGFP (GR*-eGFP) to identify genes whose knockdown led to significant changes in the cytosolic trafficking of CPMP **1**^Dex^. Cytosolic localization was monitored using the GIGT assay (Holub et al., 2013) where the reporter translocation ratio (TR) refers to the ratio of the median GFP signal in the nucleus to the median signal within a 2-μm annulus of cytosol that surrounds the nucleus (Appelbaum et al., 2012). Saos-2(GIGT) cells were transfected with pools of four siRNAs (in triplicate) for 72 hours, serum-starved, and then treated with **1**^Dex^ (1 μM) for 30 minutes. The cells were fixed, stained with Hoechst 33342, and imaged on an Opera High Content Screening System. Raw TR values associated with each well were normalized to percent effect values, relative to control cells transfected with non-targeting siRNA and treated with **1**^Dex^. We identified 428 primary hits based on the strictly standardized mean difference (β = SSMD) (Zhang et al., 2007) threshold of ±2.0. (B) Candidate genes whose knockdown inhibited the cytosolic access of **1**^Dex^ were defined by low TRs relative to control cells treated with non-targeting siRNA and **1**^Dex^ (siRNA hit type 1); candidate genes whose knockdown enhanced the cytosolic access of **1**^Dex^ were defined by high TRs relative to control cells (siRNA hit type 2); candidate genes whose knockdown led to no significant TR changes in the ability of **1**^Dex^ to reach the cytosol were excluded from further analyses (no siRNA effect). (C) An SSMD cutoff of ±2.0 resulted in an overall hit rate of 2.4%.

Before conducting the RNAi screen in a genome-wide format, we optimized cell density and siRNA transfection conditions to integrate the GIGT assay with high-content RNAi screening in Saos-2(GIGT) cells (Figure S5B). We then performed a duplicate pilot screen with 320 randomly chosen siRNAs from the Dharmacon human genome library. With this subset of siRNAs, we assessed assay performance in Saos-2(GIGT) cells treated with the CPMP **1**^Dex^ at 1 *μ*M concentration, which yielded the highest S/B (Figure S5C). This concentration is also well below 2.4 *μ*M, the lowest concentration at which some endosomal leakage was detected. Performance was robust across this siRNA test panel, with a mean signal-to-background (S/B) ratio of 2.3 when comparing the translocation ratios (TR) of Saos-2(GIGT) cells treated with **1**^Dex^ to untreated cells; the S/B ratio of an average siRNA screen is 2.9 (55). The observed coefficient of variation (CV), a measure of data variability, was also excellent; the 11.2% value observed was below the average observed in both small-molecule and siRNA screens (55). The calculated Z-factor (Z′) was 0.3, within the acceptable range for siRNA screens (55), and the reproducibility between replicate wells was high (Pearson correlation coefficient: *r* = 0.8) (Figure S5C). Collectively, these statistics are modestly better than the average of representative RNAi screens (55) and highlight the suitability of the GIGT/Opera combination assay for high-content RNAi screening.

We then used the GIGT/Opera combination assay to evaluate the effects of siRNAs targeting the majority (18,118) of human genes on the cytosolic localization of aPP5.3^Dex^ (**1**^Dex^). Saos-2(GIGT) cells were transfected in triplicate for 72 hours with pools of four siRNAs targeting different regions of the same human mRNA, serum-starved overnight, and treated with 1 *μ*M **1**^Dex^ for 30 minutes. The cells were fixed, stained with the nuclear dye Hoechst 33342, and the relative amounts of GR*-eGFP in the cytosol and nucleus quantified to generate a translocation ratio (TR) (Figure 2A). The raw TR of each experimental well was converted to a normalized percent effect value (Equation 10, see Supplementary Information). The average TR of Saos-2(GIGT) cells transfected with a non-targeting siRNA (RISC-Free) and treated with 1 *μ*M **1**^Dex^ was defined as 0% effect, while that of Saos-2(GIGT) cells without **1**^Dex^ transfected with non-targeting siRNA was defined as 100% effect.

### Hit identification and initial prioritization

Hits were identified from the data set by applying a strictly standardized mean difference (SSMD, β) threshold of 2.0 (56). This SSMD threshold identified 428 candidate genes that altered the GR*-eGFP translocation in the presence of **1**^Dex^ across three replicates (Figure 2A and 2C). The set of 428 candidate genes consisted of 165 genes whose knockdown inhibited and 263 genes whose knockdown enhanced GR*-eGFP translocation with **1**^Dex^ (Table S4). Genes whose knockdown inhibited GR*-eGFP translocation in the presence of **1**^Dex^ displayed average percent effect values between –26 and –100% (average = –44 ± 10%) compared to positive control cells treated with a non-targeting siRNA and **1**^Dex^ (Figure 2B: siRNA hit type 1). Genes whose knockdown increased GR*-eGFP translocation displayed average percent effect values between +50 and +250% (average = +96 ± 29%) higher than controls (Figure 2B: siRNA hit type 2). Genes whose knockdown failed to alter the GR*-eGFP translocation induced by **1**^Dex^ displayed mean percent effect values that remained unchanged when compared to control cells (Figure 2B: no siRNA hit). The primary hit rate of 2.4% (Figure 2C) falls within the range typically observed during cell-based RNAi screens (median primary hit rate of 2.3%) (57) and reflects the conservative nature of the SSMD value used for thresholding hits from the primary screen. While genes implicated in endocytosis (58) were enriched among the set of 428 initial hits (102 genes, 24% of the total), a protein-protein interaction analysis using the String database (59) revealed many interactions but no singular enriched mechanism or pathway.

To focus our attention on genes involved in the endosomal release of CPMP **1**^Dex^, we prioritized the set of 428 initial hits to focus on genes of unknown function (29 genes) and those implicated previously in endocytic trafficking (102 genes) (Table S5). Unknown genes were identified using the publicly available GeneCards and Rat Genome Database (RGD) databases (60, 61). Genes implicated in endocytic trafficking were identified using a previously reported systems-level survey of endocytosis (58). The 297 genes lost in this filtering step encompass diverse functions. To confirm that we had successfully prioritized genes of unknown function and those implicated previously in endocytic trafficking, we cross-referenced the set of 131 prioritized genes with the Database for Annotation, Visualization and Integrated Discovery (DAVID) (62, 63). We specified the gene ontology classification “cellular compartment” to evaluate the “enrichment” of this set of 131 genes with respect to categories of subcellular organelles. The most enriched organelle categories, according to the database analysis, were the Golgi stack, the lysosomal membrane, and the late endosome. This analysis provides confidence that the set of 131 prioritized genes were worthy of subsequent study.

We recognized that some of the remaining siRNAs could be associated with the GR signaling pathway and affect the cytosolic to nuclear distribution of GR*-eGFP even in the absence of Dex or **1**^Dex^. To identify these genes, we evaluated the individual effect of each of the four gene-specific siRNAs targeting the remaining 131 candidate genes on the distribution of GR*-GFP in non-treated Saos-2(GIGT) cells. We discarded a gene if three of the four gene-specific siRNAs significantly increased the TR compared to that measured in Saos-2(GIGT) cells transfected with a non-targeting siRNA, indicating that the gene knockdown primarily affects GR*-GFP distribution independent of the cytosolic presence of Dex or a Dex-labeled peptide. This process eliminated 61 genes from consideration (Table S6), leaving 70 genes for subsequent validation.

Next, focusing on these remaining 70 genes, we evaluated the individual effect of each of the four gene-specific siRNAs on the TR measured in Saos-2(GIGT) cells treated with either CPMP **1**^Dex^ or **2**^Dex^, which both carry the discrete arginine array that exemplifies a penta-arg motif. We reasoned that this siRNA deconvolution procedure would identify those genes involved in cellular trafficking of both **1**^Dex^ and **2**^Dex^, and simultaneously minimize false positives that result from siRNA off-target effects (64). Genes were retained if at least two of the four siRNAs in the pool led to significant TR changes (by one-way ANOVA test with Dunnett post hoc test) in the presence of either **1**^Dex^ or **2**^Dex^ when compared to that observed in Saos-2(GIGT) cells transfected with a non-targeting siRNA; 28 of the 70 candidate genes passed this filter (Table S7). This final set of 28 genes displayed significant enrichment for membrane compartments (19 of 28 genes) when specifying the gene ontology term “cellular compartments,” but did not exhibit enriched protein-protein interaction networks (59).

### Prioritizing hits based on mechanism: genetic knockdowns that selectively mediate endosomal escape

Although the GIGT/Opera combination assay is useful for analyzing the effect on siRNA knockdowns on cytosolic access in a high-throughput mode, it is inherently qualitative and does not differentiate between knockdowns that alter CPMP/CPP uptake from those that only affect endosomal escape. To further validate and differentiate between the remaining 28 candidate genes on the basis of these criteria, we turned to two quantitative methods: flow cytometry (FC) and FCS (19). When used together, FC and FCS effectively discriminate knockdowns that alter overall CPMP/CPP uptake (quantified by FC) from those that affect endosomal escape (quantified by FCS). To evaluate the effects of each siRNA knockdown on the trafficking of CPMPs using FC and FCS, we chose to focus on CPMP **2**^R^—while CPMPs **1**^R^ and **2**^R^ follow a similar endocytic trafficking pattern into the cell interior (18), **2**^R^ is more stable and reaches the cytosol more efficiently (19).

To evaluate the effect of the 28 candidate genes on CPMP uptake and endosomal release, Saos-2 cells were treated with pooled gene-specific siRNAs for 72 h. Transfected Saos-2 cells were then treated with 600 nM **2**^R^ for 30 minutes, lifted with trypsin to remove plasma membrane-bound peptide (18), and analyzed using FC and FCS to quantify both overall cellular uptake and the concentration of **2**^R^ that reached the cell interior (Figure 3A). Real-time quantitative polymerase chain reaction (RT-qPCR) experiments confirmed that the knockdown efficiency of each siRNA pool was >70% in Saos-2 cells (Figure S6A). Additional control experiments confirmed that neither RNAiMAX nor knockdown of the housekeeping gene *GAPD* affected overall CPMP uptake by Saos-2 cells (as determined by FC) or intracellular delivery of **2**^R^ (as determined by FCS) (Figure S6B).

**Figure 3:**
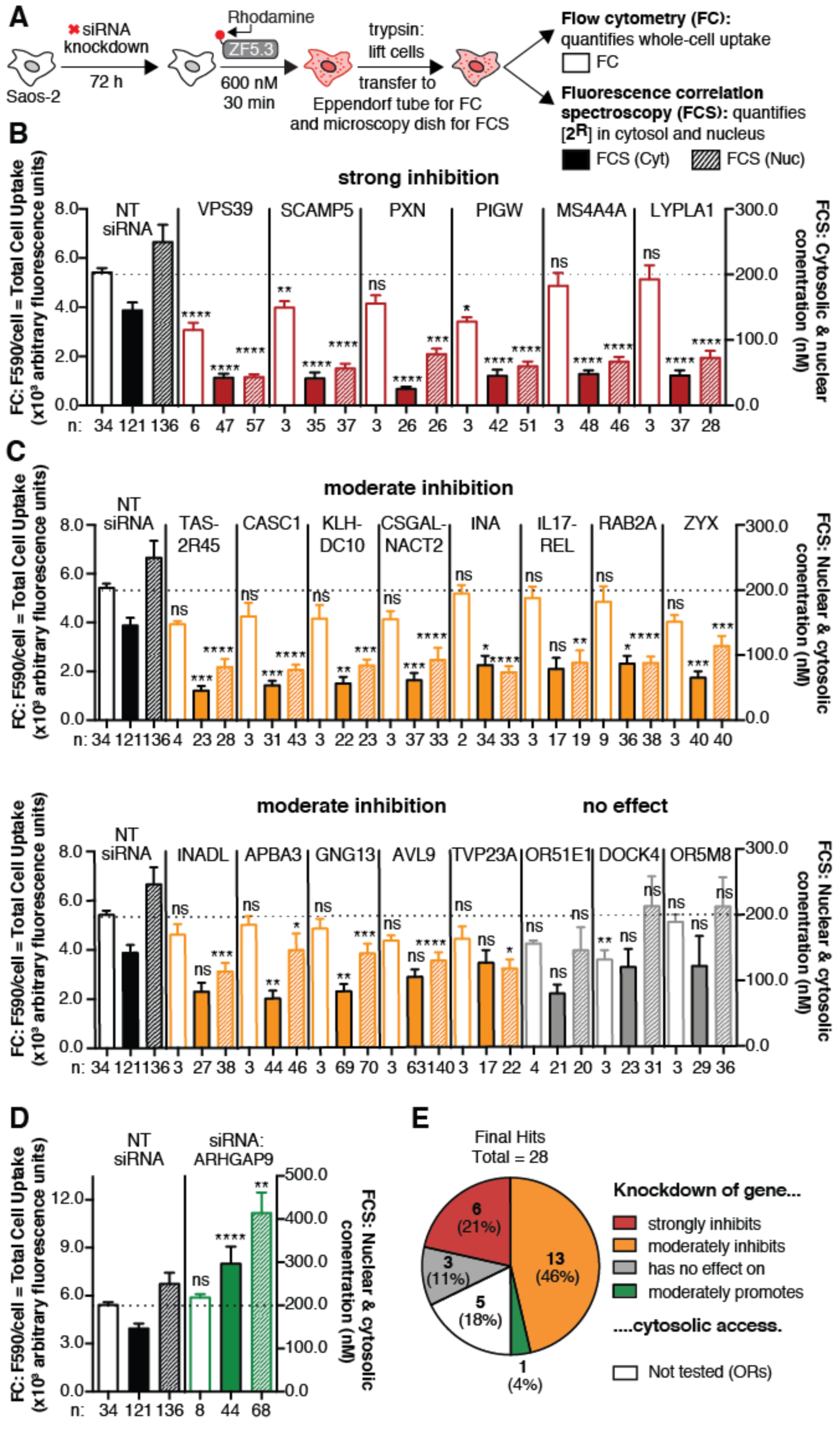
Categorizing candidate genes using flow cytometry (FC) and fluorescence correlation spectroscopy (FCS). (A) Knockdowns were achieved by transfecting Saos-2 cells with siGENOME SMARTpool siRNA and Lipofectamine RNAiMAX for 72 h according to the manufacturer’s protocol. Cells were treated with **2**^R^ (600 nM) for 30 minutes and exogenously bound peptide was removed with TrypLE Express. Cells were then evaluated using FC to assess the levels of whole-cell fluorescence intensity and using FCS to calculate the concentration of **2**^R^ in the cytosol and nucleus. (B–D) FC and FCS data illustrating the effects of gene knockdowns on both total cellular fluorescence and the cytosolic localization of **2**^R^ relative to the effects of non-targeting (RISC-Free) siRNA (NT siRNA). Knockdown of genes shown in (B) strongly inhibit (>70%) intracellular (cytosolic and nuclear) access of **2**^R^. Knockdown of genes shown in (C) moderately inhibit (30–70%) intracellular access of **2**^R^. Knockdown of the *ARHGAP9* gene shown in (D) promotes (149%) cytosolic access of **2**^R^. Left axis: FC = total cell uptake, fluorescence intensity per cell at 590 nm. Each data point (n) represents one biological replicate. For each FC replicate, the median fluorescence intensity at 590 nm was measured for at least 10,000 Saos-2 cells (gated for live cells). Right axis: FCS: cytosolic and nuclear concentration (nM). Each data point (n) denotes a 50-second FCS measurement recorded in a single cell. Error bars represent the standard error of the mean. ****p < 0.0001, ***p < 0.001, **p < 0.01, *p < 0.05, and not significant (ns) for p > 0.05 from one-way ANOVA with Dunnett post hoc test. (E) Of the 28 final hits identified in the GIGT high-throughput screen, knockdown of six genes strongly inhibited cytosolic access, knockdown of 13 genes moderately inhibited cytosolic access, knockdown of one gene promoted cytosolic access, and knockdown of three genes had no effect on cytosolic access (false positives). The remaining five genes, which were classified as false positives, belong to a group of seven olfactory receptors (ORs), which exhibit high sequence homology (>40%) among each other. Two representative ORs were tested and displayed no significant effects compared to NT siRNA.

Knockdown of the 28 candidate genes led to significant differences in the overall uptake and intracellular delivery of **2**^R^. Knockdown of only four of the 28 candidate genes (*VPS39*, *SCAMP5*, *PIGW*, and *DOCK4*) significantly decreased the overall uptake of **2**^R^, as determined using FC (left axis in Figure 3B–D); none of the siRNAs significantly increased overall uptake when measured in this way (Figure 3B–D). Different trends emerged when siRNA knockdown effects were evaluated using FCS (right axis in Figure 3B–D); in this case 19 of 28 knockdowns led to a significant decrease in the concentration of **2**^R^ in the cytosol or nucleus. Knockdown of six of the 28 candidate genes strongly reduced (>70%) the delivery of **2**^R^ to the nucleus and cytosol: these genes include *VPS39*, *SCAMP5*, *PXN*, *PIGW*, *MS4A4A*, and *LYPLA1* (Figure 3B). Knockdown of an additional 13 candidate genes moderately reduced (40–70%) the delivery of **2**^R^ to the nucleus and cytosol: these genes include *TAS2R45*, *CASC1*, *KLHDC10*, *CSGALNACT2*, *INA*, *IL17REL*, *RAB2A*, *ZYX*, *INADL*, *ABPA3*, *GNG13*, *AVL9*, and *TVP23A* (Figure 3C). Notably, knockdown of one candidate gene, *ARHGAP9*, led to a significant (+49%) increase in the delivery of **2**^R^ to the nucleus and cytosol (Figure 3D); this finding suggests that inhibitors of ARHGAP9 could improve endosomal release. It is also notable that genes whose knockdown strongly reduced the delivery of **2**^R^ encompass multiple cellular activities related to membrane homeostasis, including membrane tethering (*VPS39*, *RAB2A*), glycosylphosphatidylinositol (GPI)-anchor biosynthesis (*PIGW*), thioesterases (*LYPLA1*), and cytoskeletal genes (*PXN*, *ZYX*). The remaining eight genes consisted of *DOCK4* and seven olfactory receptors (ORs: *OR4C6*, *OR4F15*, *OR51E1*, *OR51Q1*, *OR52N1*, *OR5M8*, *OR8D1*), which were eventually classified as false positives (Figure 3C). In summary, of the 28 genes identified in the GIGT high-throughput screen, knockdown of six strongly inhibited cytosolic access, knockdown of 13 moderately inhibited cytosolic access, and knockdown of one gene promoted cytosolic access (Figure 3E). These data provide evidence that we successfully screened and prioritized genes that primarily regulate intracellular trafficking—with 20 out 28 genes affected by FCS—and not overall uptake—with only four genes affected by FC.

Hydrocarbon-stapled α-helical peptides are a promising class of therapeutic candidates because they can exhibit improved pharmacological properties—such as increased binding affinity, proteolytic resistance, and cell permeability—compared to staple-free analogs (65-67). Although hydrocarbon-stapled peptides lack the defined array of arginine side chains that characterizes CPMPs **1** and **2**, they reach the cytosol with high efficiency. In particular, the hydrocarbon-stapled peptide SAH-p53-8^R^ (**3**^R^) achieves average intracellular concentrations (as determined by FCS in HeLa or Saos-2 cells) that are only 10–50% lower than those attained by ZF5.3^R^ (**2**^R^), depending on the cell line used (19). To investigate whether **2** and **3** share a common mode of endosomal release, Saos-2 cells were treated with pooled siRNAs targeting the set of 20 candidate genes, treated with **3**^R^ (600 nM), and both whole-cell uptake and cytosolic delivery were quantified with FC and FCS as described above (Figure S6C).

A plot showing the percent effect of each statistically significant knockdown (that is, significant for both **2**^R^ and **3**^R^ by one-way ANOVA with Dunnett post hoc test) on the cytosolic localization of **2**^R^ or **3**^R^ showed moderate correlation (Pearson *r* = 0.47, P = 0.17) (Figure S7A). Knockdown of five genes—*VPS39*, *INA*, *LYPLA1*, *KLHDC10*, and *CASC1* strongly (>50%) inhibited the appearance of both **2**^R^ and **3**^R^ in the cytosol (blue open circles). Knockdown of another four genes—*MS4A4A*, *TAS2R45*, *RAB2A*, and *APBA3*—resulted in a moderate (30–50%) decrease in the appearance of both **2**^R^ and **3**^R^ in the cytosol (green dots). Knockdown of AVL9, although statistically significant, did not have a strong %-effect on the ability of either **2**^R^ or **3**^R^ to reach the cytosol (Figure S7A). When the effects of these gene knockdowns on overall uptake of **2**^R^ and **3**^R^ are plotted, the effects are small, as expected (Figure S7B). Overall, the observed correlation between the effects of gene knockdowns on the cytosolic localization of CPMP **2**^R^ and hydrocarbon-stapled peptide **3**^R^ suggests that these genes play common roles in the intracellular trafficking of these two molecular families. The five genes that most strongly regulate cytosolic delivery of both **2**^R^ or **3**^R^ can be divided into four categories: (1) genes with known roles in endocytic trafficking (*VPS39*); (2) genes involved in the processing of post hoc translational protein modifications (*LYPLA1)*, (3) genes involved in RNA processing (*CASC1*), and (4) genes with poorly characterized or unknown functions (*INA* and *KLHDC10*).

### siRNA depletion of HOPS but not CORVET inhibits the cytosolic delivery of CPPs/CPMPs 1^R^–5^R^

The goal of the RNAi screen was to identify candidate genes that significantly enhance or inhibit the intracellular localization of multiple CPMPs and CPPs. One gene transcript, when depleted, strongly inhibited the cellular uptake and cytosolic delivery of both CPMP **2**^R^ and stapled peptide **3**^R^: the *VPS39* gene, also known as hVamp6. *VPS39* encodes an 875-residue protein that is conserved in eukaryotes (68) and is one of six protein subunits comprising the homotypic fusion and vacuole protein sorting (HOPS) membrane tethering complex (Figure 4A). Working with VPS41 and various effector proteins (69, 70), VPS39 recruits the HOPS complex to endosomes positive for the small GTPase Rab7, such as maturing (Rab5^+^/Rab7^+^) and late (Rab7^+^) endosomes. The remaining four core HOPS subunits, VPS11, VPS16, VPS18, and VPS33A, serve primarily a structural role (70). A fully assembled HOPS complex is required to initiate fusion of Rab7^+^ maturing and late endosomes as they are converted into lysosomes (Figure 6B). Depletion of any subunit delays lysosome maturation and inhibits cargo degradation (Figure 4B) (70-72). HOPS is closely related to a structurally similar complex, the CORVET complex (Figure 4A), that precedes HOPS along the endocytic pathway by promoting the fusion of Rab5^+^ early and maturing endosomes (73). CORVET shares four core subunits with HOPS but contains TGFBRAP1 and VPS8 (73) in place of VPS39 and VPS41.

**Figure 4:**
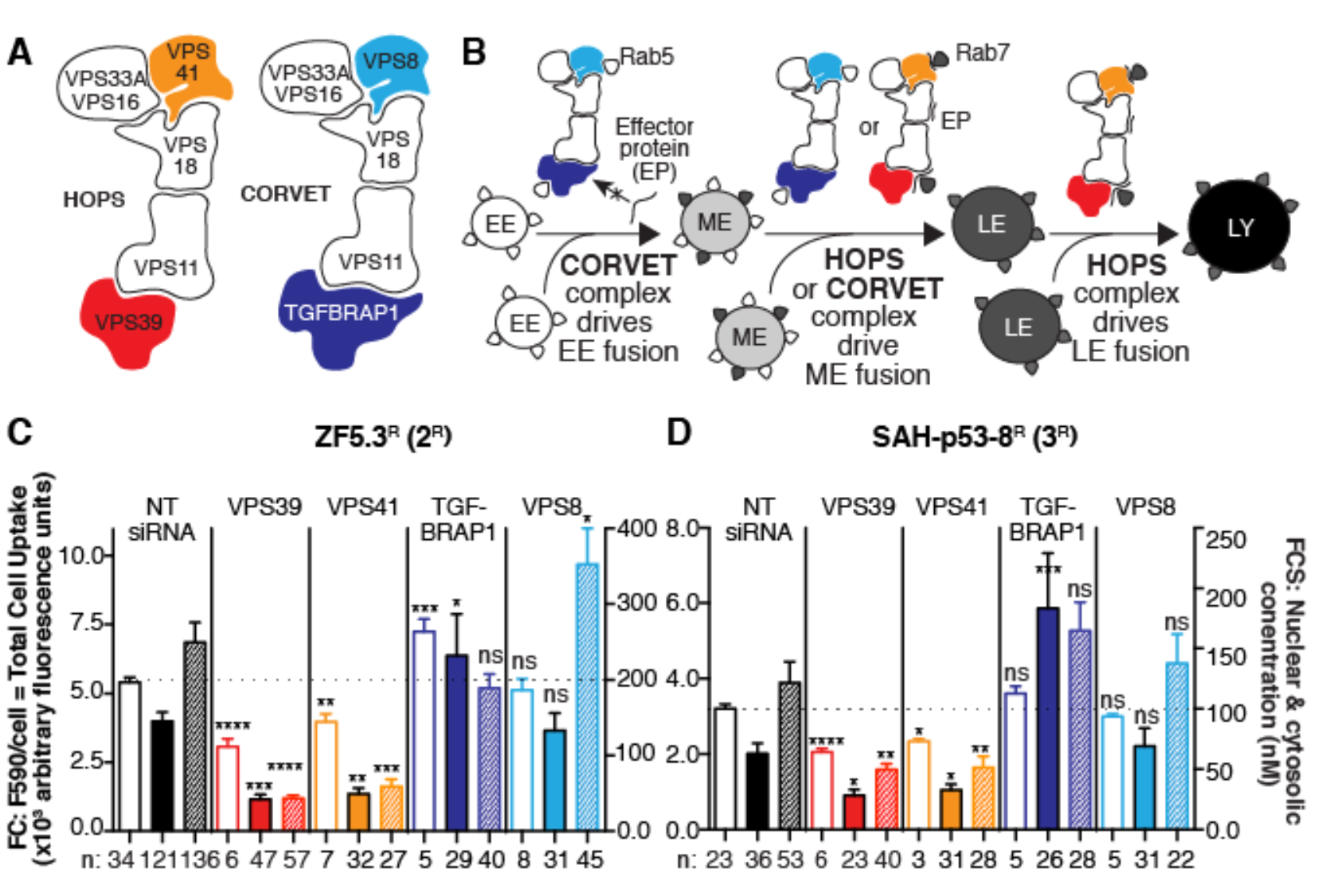
Efficient trafficking of both CPMPs and hydrocarbon-stapled peptides requires the human HOPS complex. (A) Cartoons illustrating the protein components of the human HOPS and CORVET complexes. (B) CORVET and HOPS initiate endosomal contact and fusion along the degradative endocytic pathway. CORVET drives the fusion of early endosomes (EEs) and maturing endosomes (MEs), while HOPS drives fusion of MEs and late endosomes (LEs) to form lysosomes (LYs). (C and D) The effect of VPS39 and VPS41 knockdown (HOPS-specific subunits) compared to TGFBRAP1 and VPS8 knockdown (CORVET-specific subunits) on both overall uptake (FC) and cytosolic access (FCS) of **2**^R^ (C) and of **3**^R^ (D) relative to the effect of non-targeting siRNA (NT siRNA). Left axis: FC = total cell uptake, fluorescence intensity per cell at 590 nm. n refers to the number of biological replicates. For each FC replicate, the median fluorescence intensity at 590 nm was measured for at least 10,000 Saos-2 cells (gated for live cells). Right axis: FCS: cytosolic and nuclear concentration (nM). Each data point (n) denotes one 50-second FCS measurement recorded in the nucleus or cytosol of a single Saos-2 cell. Error bars represent the standard error of the mean. ****p < 0.0001, ***p < 0.001, **p < 0.01, *p < 0.05, not significant (ns) for p > 0.05 from one-way ANOVA with Dunnett post hoc test.

Having identified VPS39 as a key regulator of the cytosolic delivery of **2**^R^ and **3**^R^, we next asked whether the observed effect was dependent on VPS39 alone or on the other unique protein subunit of the HOPS complex: VPS41. Saos-2 cells transfected with a non-targeting (RISC-Free) or a pool of four VPS41-targeting siRNAs were treated with **2**^R^ or **3**^R^ (600 nM) for 30 minutes and analyzed using both FC and FCS as described above (Figure 4C and 4D). In our primary screen, we found that siRNAs against VPS41 displayed a mild inhibitory effect on GR*-eGFP translocation in the presence of **1**^Dex^ (27% mean percent effect, 1.58 Z-score, 1.65 SSMD), which was below our hit threshold. Here, upon VPS41 depletion, we observed a moderate decrease (–27%) in overall uptake and a strong (–71%) decrease in the average intracellular concentration for **2**^R^ (Figure 4C); these changes are comparable to those observed after VPS39 knockdown. In VPS41-depleted cells treated with hydrocarbon-stapled peptide **3**^R^, we also observed a significant decrease in overall uptake (–27%) and in the average intracellular delivery (–55%) by FCS (Figure 4D); these changes are also similar to those observed after VPS39 knockdown. By contrast, no significant decreases in overall uptake or cytosolic localization of **2**^R^ or **3**^R^ were observed upon depletion of the CORVET-specific subunits TGFBRAP1 and VPS8 (Figure 4C and 4D); in fact, depletion of the TGFBRAP subunit led to significant increases in both overall uptake and cytosolic concentration of **2**^R^ and **3**^R^ (Figure 4C and 4D). Taken together, these data support a hypothesis in which both unique components of the HOPS complex, VPS39 and VPS41, are required to allow cytosolic access of both CPMP **2**^R^ and hydrocarbon-stapled peptide **3**^R^. Similar trends were observed in comparable experiments performed using CPMP **1**^R^ (Figure S8A) implying that HOPS-dependent membrane tethering is also necessary for the efficient delivery of CPMP **1**^R^ to the cytosol.

Because of the unusual diffusion dynamics observed for CPPs **4**^R^ and **5**^R^ in cultured cells (Figure S4B), we were unable to perform FCS experiments to evaluate whether their delivery to the cytosol required HOPS subunits or activity. We were able to estimate their relative delivery into the cytosol on the basis of fluorescence intensity values obtained during FCS measurements (without autocorrelation); these values are proportional to the intracellular concentration in the confocal volume (Figure S8E). Knockdown of either VPS39 or VPS41 led to a significant (23–35%) decrease in whole-cell uptake of **4**^R^ or **5**^R^ (as determined by FC) and a 46–62% decrease in their average cytosolic fluorescence intensity (as determined by FCS without autocorrelation) (Figure S8B and S8C). In contrast, knockdown of the CORVET subunits TGFBRAP1 or VPS8 led to either no change (**5**^R^) or a 30% increase (**4**^R^) in overall uptake (as determined by FC) (Figure S8B and S8C). Knockdown of TGFBRAP1 led to no change in the average cytosolic fluorescence intensity of **4**^R^ or **5**^R^ (as determined by FCS without autocorrelation), while knockdown of VPS8 either stimulated (**4**^R^) or had no effect (**5**^R^) on the average cytosolic fluorescence intensity as determined by FCS without autocorrelation.

Taken together, these data show that HOPS-specific subunits VPS39 and VPS41, but not CORVET-specific subunits TGFBRAP1 and VPS8, are required for the cytosolic delivery of all CPMP/CPPs tested herein. These CPMPs and CPPs differ significantly in their intrinsic ability to reach the cytosol (Figure S4A), but in all cases this ability demands an active HOPS complex. Importantly, the more active CPMPs **1** and **2** are significantly more sensitive to knockdown of the HOPS subunit VPS39 than hydrocarbon-stapled peptide **3** or CPPs **4** and **5**; knockdown of VPS39 led to 79% and 74% decrease in the cytosolic concentrations of CPMPs **1** and **2,** whereas the decreases observed for hydrocarbon-stapled peptide **3** and CPPs **4** and **5** ranged from 49–56%. These data suggest that even though all peptides require HOPS-dependent fusion of late endosomes to access the cytosol, CPMPs **1** and **2** may be more strongly dependent on this pathway—or make more efficient use of it—than hydrocarbon-stapled peptides and CPPs **3**–**5**.

### HOPS remains active upon treatment with CPMPs and CPPs 1–5

Although the experiments described above indicate that efficient cytosolic delivery of CPMP/CPPs **1**–**5** demands the *presence* of a fully assembled HOPS complex, they do not discriminate between two limiting explanations for this dependence: one in which cytosolic access demands the *activity* of the HOPS complex and another in which cytosolic access demands the *inhibition* this activity. To discriminate between these two possibilities, we monitored HOPS activity in the presence of CPMPs **1**^UL^–**5**^UL^ using a previously reported fluorescence colocalization protocol that quantifies HOPS-dependent delivery of dextran to lysosomal compartments (71, 72). In this assay, the endocytic system is flooded with Alexa Fluor 488-tagged dextran for 2 h, after which the cells are washed and stained with Hoechst 33342, re-plated in dextran-free media, incubated for 1 h, stained with Magic Red, and imaged immediately by confocal microscopy (Figure S9A). Magic Red is a dye that becomes fluorescent in the presence of active cathepsin B—an enzyme exclusively localized to mature lysosomes. Dextran is only delivered to lysosomal compartments if endosomal fusion of Rab7^+^ endosomes occurs. This process requires a fully assembled and functional HOPS complex (71, 72).

To validate this assay, we first measured the effect of HOPS-specific subunit depletion on the colocalization of Magic Red with dextran. Saos-2 cells were transfected with siRNA pools targeting either VPS39 or VPS41, or with a non-targeting siRNA as a negative control. After 72 hours, cells were treated as described above and the colocalization of dextran and Magic Red was calculated using ImageJ (44). In untreated and non-targeting siRNA-treated Saos-2 cells, the colocalization of Alexa Fluor 488-dextran and Magic Red was characterized by Manders coefficients of 0.30 ± 0.030 and 0.23 ± 0.020, respectively; these values represent the fraction of dextran-containing vesicles that also contain cathepsin B and are comparable to those measured previously (74) (Figure S9B and S9C). As expected, depletion of HOPS-specific subunits VPS39 and VPS41 significantly reduced the colocalization of Alexa Fluor 488-dextran and Magic Red, leading to significantly lower Manders coefficients of 0.11 ± 0.024 and 0.12 ± 0.016, respectively (Figure S9B and S9C).

With a validated assay in hand, we next asked whether addition of CPMPs or CPPs **1**^UL^–**5**^UL^ had an observable effect on the intrinsic activity of the HOPS complex. Saos-2 cells were pre-treated with unlabeled CPMPs **1**^UL^–**5**^UL^ at 600 nM for 30 min and labeled with Alexa Fluor 488-dextran and Magic Red as described above (Figure S9D). In cells treated with CPMPs/CPPs **1**^UL^–**5**^UL^, the colocalization of Alexa Fluor 488-dextran and Magic Red was characterized by Manders coefficients that ranged from 0.27 ± 0.03 (when treated with **5**^UL^) to 0.33 ± 0.03 (when treated with **1**^UL^), well within the range of values measured in cells that had not been treated with CPMPs or CPPs (Figure S9E and S9F). We conclude that the treatment of Saos-2 cells with CPMPs **1**–**5** neither inhibits nor enhances the intrinsic function of HOPS-mediated endosomal fusion. As a whole, our studies suggest that the endosomal escape of all CPMPs and CPPs tested is dependent on the presence of a functional, active HOPS complex.

### How does HOPS mediate endosomal escape?

HOPS is required for the fusion of Rab7^+^ endosomes, a process initiated by HOPS-mediated membrane tethering and completed by the mechanical action of SNAP (soluble NSF(*N*-ethylmaleimide-sensitive factor) attachment protein) receptor (SNARE) proteins and the HOPS complex itself (75). Depletion of HOPS-specific subunits significantly decreases the cytosolic and nuclear concentrations of all CPMP/CPPs tested. Three limiting models are consistent with these experimental findings. The first invokes a direct interaction between a CPMP/CPP located within or on the surface of a Rab7^+^ late endosome and a component of the HOPS complex, perhaps VPS39 or VPS41, that effectively shuttles the CPMP or CPP from an endosome into the cytosol (model 1). The second model also positions the CPMP/CPP within or on the surface of a Rab7^+^ late endosome but lacks a direct HOPS-CPMP/CPP interaction; in this case HOPS facilitates escape of the CPMP/CPP during the fusion process (76, 77) (model 2). In the third model, the CPMP or CPP escapes after HOPS-dependent fusion is complete, when maturing and late endosomes have fused into compartments containing intraluminal vesicles (ILVs) (model 3). In this case, interactions between CPMPs and ILVs in the late endosome may facilitate endosomal escape; indeed, some evidence for CPP-ILV interactions have been reported (50).

Implicit in models 1 and 2 is the colocalization of a CPMP/CPP with or upon the membrane of a Rab7^+^ late endosome. Confocal microscopy of Saos-2 cells expressing either the early endosomal (EE) marker Rab5-GFP, the late endosomal marker (LE) Rab7-GFP, or the lysosomal (LY) marker Lamp1-GFP and treated with **2**^R^ (300 nM, 30 min) revealed low colocalization between **2**^R^ and EEs (Pearson *r* = 0.20 ± 0.019) but high colocalization of **2**^R^ with LEs (*r* = 0.55 ± 0.034) and LYs (*r* = 0.58 ± 0.027) (Figure 5A, 5B, 5D). However, these experiments could not resolve whether the fluorescence of **2**^R^ was localized to the endosomal lumen or the membrane. To better resolve the location of **2**^R^, we treated Saos-2 cells with the small molecule YM201636, an inhibitor of the phosphoinositide kinase PIK-fyve that enlarges Rab7^+^ endolysosomes (78, 79). Again, in cells expressing GFP-labeled endosomal markers, we observed virtually no colocalization between **2**^R^ and EEs (*r* = 0.034 ± 0.025), good colocalization with LEs (*r* = 0.57 ± 0.043), and excellent colocalization with LYs (*r* = 0.73 ± 0.023) (Figure 5C and 5D). Line profiles across these enlarged vesicles provide no evidence for accumulation of **2**^R^ on the LE or LY membrane; instead, plots of fluorescence intensity as a function of position show clearly separable distributions of fluorescence due to GFP (green) and lissamine rhodamine B (red) (Figure 5E and 5F). These distributions suggest that **2**^R^ resides primarily within Rab7^+^ and Lamp1^+^ vesicles and not on their surface and is inconsistent with models 1 or 2 as the primary means of endosomal escape.

**Figure 5:**
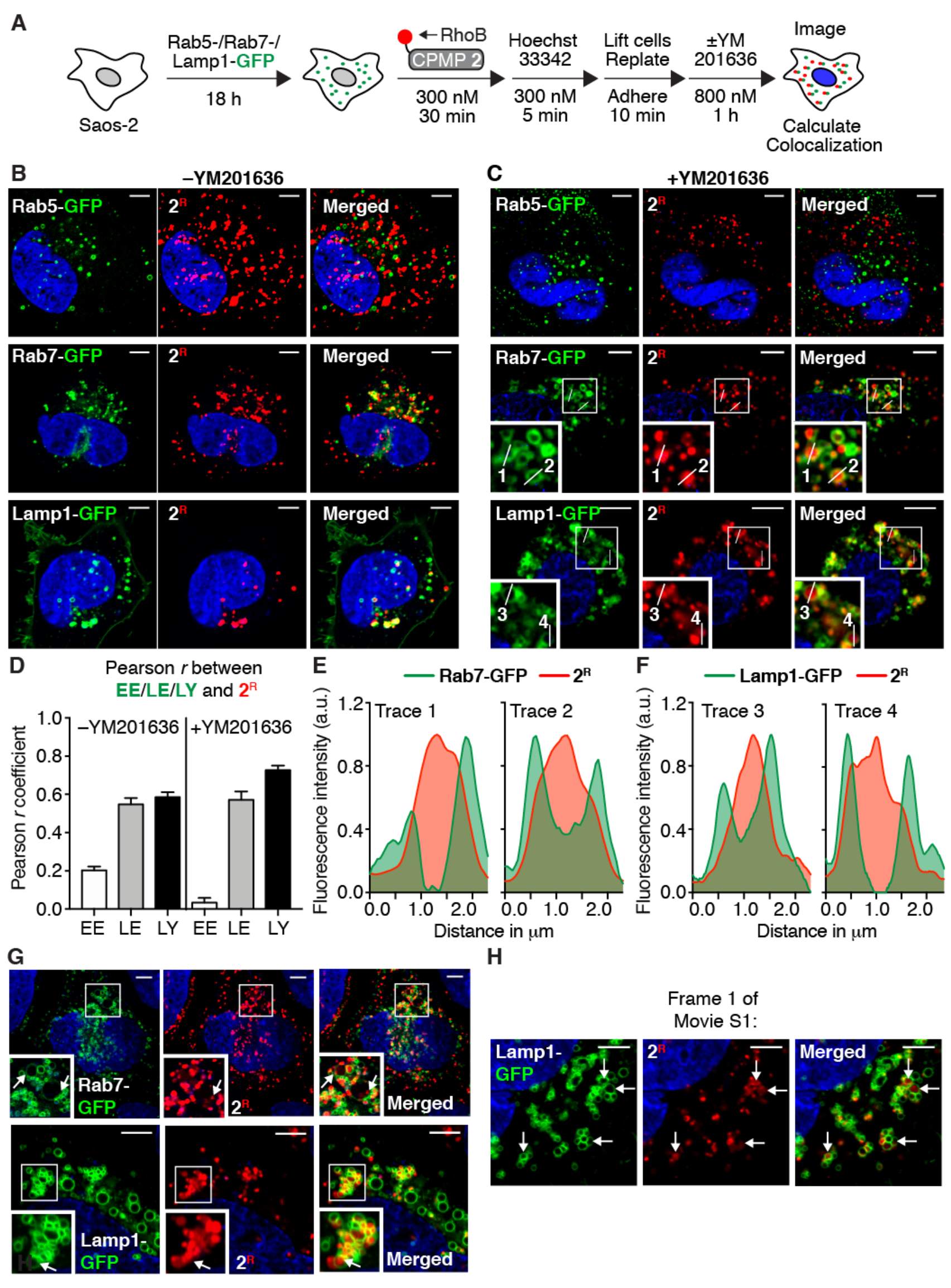
CPMP 2 localizes to the lumen of LEs and LYs. (A) Saos-2 cells were transduced with Rab5-, Rab7-, or Lamp1-GFP for 18 h using CellLight Reagents (BacMam 2.0). Cells were washed, incubated with CPMP 2^R^, stained with Hoechst 33342, lifted with TrypLE, and re-plated into microscopy slides. Cells were then incubated in media ±YM201636 for 1 hour. (B and C) Representative live-cell confocal fluorescence microscopy images of Saos-2 cells prepared as described in (A) in media without (B), or with (C) YM201636. (D) Pearson correlation coefficients characterizing colocalization of GFP markers and 2^R^ in the presence and absence of YM201636. (E and F) Fluorescence intensity line profiles of endosomes 1–4 (see panel C). showing the relative location of emission due to 2^R^ (red) and either (E) Rab7-GFP or (F) Lamp1-GFP. (G) Representative live-cell confocal fluorescence microscopy images of Saos-2 cells prepared as described in (A) in the presence of YM201636. White arrows identify smaller vesicles present within the boundaries of Rab7-GFP^+^ or Lamp1-GFP^+^ cells. (H) Representative live-cell confocal fluorescence image of the first frame of Movie S1. Arrows identify Lamp1-GFP^+^ endosomes that contain moving vesicles labeled with CPMP 2^R^ (+YM201636), as shown in Movies S1 and S2.

Closer examination of enlarged endosomes in Rab7- or Lamp1-GFP-expressing cells revealed the presence of smaller vesicles within the boundaries of late endosomal membranes, as shown in Figure 5G and 5H and in two movies of cells expressing Lamp1-GFP and treated with **2**^R^ (white arrows in Figure 5G and 5H and Movies S1 and S2). Late endosomes are characterized, among other properties, by the presence of ILVs. To determine whether the smaller **2**^R^-containing vesicles observed in late endosomes represent ILVs, we used a known ILV marker, the lipid *N*-Rh-PE, a lissamine rhodamine B-tagged version of 1,2-dipalmitoyl-*sn*-glycero-3-phosphoethanolamine (80) (Figure S10A). *N*-Rh-PE is internalized via the endocytic pathway into ILVs (81-83) before it is eventually secreted in exosomes (80, 82, 84). As expected (80), *N*-Rh-PE localizes to the luminal side of Rab7-GFP^+^ late endosomes (*r* = 0.39 ± 0.050) and Lamp1-GFP^+^ lysosomes (*r* = 0.45 ± 0.031), but not to Rab5-GFP^+^ early endosomes (*r* = 0.104 ± 0.004) (Figure S10B and S10D). To image *N*-Rh-PE and CPMP **2** simultaneously, we tagged **2** with a silicon-rhodamine (SiR) dye that emits at 660–670 nm (85) (lissamine rhodamine B emits at 585–595 nm). Saos-2 cells treated with **2**^SiR^ alone showed a colocalization profile that mirrored that of **2**^R^, as expected (compare Figure 5B/5D to S10C/S10E). Saos-2 cells treated with both *N*-Rh-PE and **2**^SiR^ showed significant colocalization (*r* = 0.53 ± 0.028) (Figure S10F and S10G) providing evidence that CPMP **2**^SiR^ indeed localizes to ILVs or to LEs and LYs that contain ILVs, providing support for model 3.

We also performed a four-color confocal microscopy experiment to visualize Saos-2 cells expressing either an EE, LE, or LY marker (as a GFP fusion) and treated with *N*-Rh-PE, **2**^SiR^, as well as Hoechst 33342. As expected, we observed virtually no colocalization of **2**^SiR^ with EEs (*r* = 0.071 ± 0.014) but good colocalization with LEs (*r* = 0.35 ± 0.036) and LYs (*r* = 0.41 ± 0.037). Also as expected, we observed minimal colocalization of *N*-Rh-PE with EEs (*r* = 0.078 ± 0.023) and good colocalization with LEs (*r* = 0.39 ± 0.039) and LYs (*r* = 0.43 ± 0.045). Good colocalization between **2**^SiR^ and *N*-Rh-PE was observed in Saos-2 cells regardless of which endosomal marker was present (Figure S11A and S11C). To enlarge Rab7^+^ endolysosomes for better resolution, we again treated Saos-2 cells with YM201636 (78, 79) (Figure S11B). Pearson *r* values were similar in cells with and without YM201636 present (Figure S11D). By generating line profiles of endosomes displayed in Figure S11B, we again observed minimal overlap between GFP markers for Rab7 or Lamp1 and either CPMP **2**^SiR^ or ILV marker *N*-Rh-PE (Figure S11E and S11F). These results further support the conclusion that CPMP **2** localizes within (as opposed to on the surface of) late endosomes and lysosomes along with a marker for ILVs and provides additional support for model 3.

### HOPS is required to deliver CPMP 2 to LEs and LYs

The observation that **2** colocalizes with *N*-Rh-PE, a lipid marker for ILVs, along with the absence of evidence that CPMP **2** localizes within the late endosomal membrane, provided circumstantial evidence for model 3, in which HOPS is required to deliver CPMPs into a cell compartment that then facilitates endosomal escape. If so, then knockdown of HOPS components (but not CORVET components) should reduce trafficking of **2**^SiR^ to LEs and LYs. To test this hypothesis, we transfected Saos-2 cells with siRNAs against HOPS subunit VPS39 and CORVET subunit TGFBRAP1. In control cells transfected with a non-targeting (RISC-Free) siRNA, we measured virtually no colocalization between **2**^SiR^ and EE marker Rab5-GFP (*r* = 0.05 ± 0.01), moderate colocalization with LE marker Rab7-GFP (*r* = 0.25 ± 0.016), and good colocalization with LY marker Lamp1-GFP (*r* = 0.57 ± 0.027) (Figure 6A and 6D). In HOPS-deficient cells, we also observed virtually no colocalization of **2**^SiR^ with EEs (*r* = 0.05 ± 0.01) and moderate colocalization with LEs (*r* = 0.21 ± 0.016). The colocalization with LYs (*r* = 0.35 ± 0.026), on the other hand, was significantly lower compared to control cells (Figure 6B and 6D). No such changes were observed in CORVET-deficient cells (virtually no colocalization of **2**^SiR^ with EEs (*r* = 0.083 ± 0.013); moderate colocalization with LEs (*r* = 0.21 ± 0.015), high colocalization with LYs (*r* = 0.47 ± 0.021) (Figure 6C and 6D). These data suggest that knockdown of HOPS but not CORVET inhibits trafficking of **2**^SiR^ to Lamp1^+^ LEs and LYs and is consistent with model 3, in which CPP/CPMPs require HOPS to reach a favorable environment for escape.

**Figure 6:**
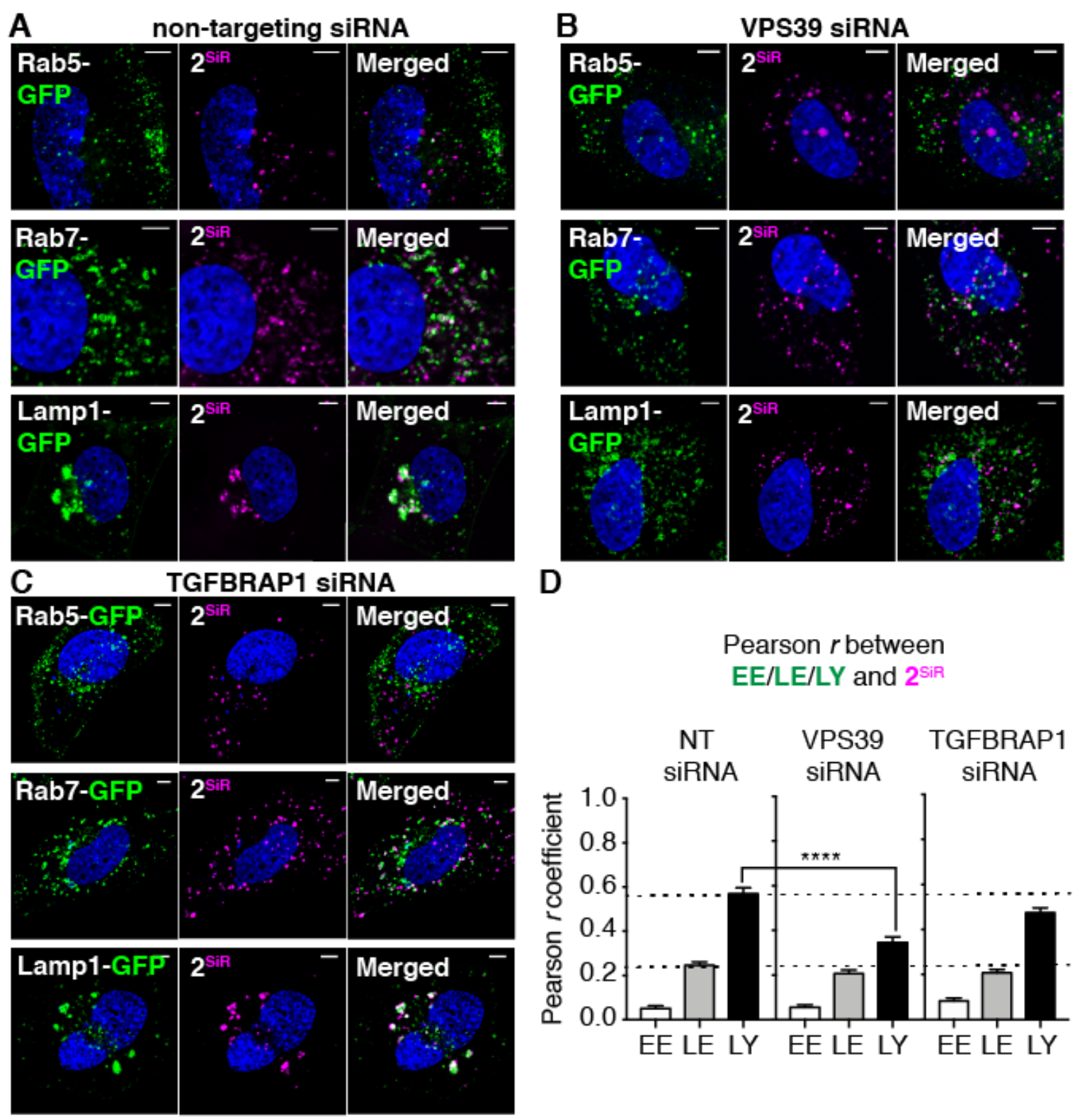
HOPS knockdown inhibits trafficking of CPMP 2 to Lamp1^+^ LEs and LYs. Saos-2 cells were transfected with pooled siRNAs against VPS39 or a non-targeting (NT) control siRNA (RISC-Free) for 72 h. Rab5-, Rab7-, or Lamp1-GFP were expressed by transduction with CellLight Reagents (BacMam 2.0) for 18 h. Cells were incubated with CPMP **2**^SiR^ (600 nM) for 30 min and nuclei were stained with Hoechst 33342 (300 nM) for 5 min. Cells were lifted using TrypLE Express to remove exogenously bound CPMP and re-plated for confocal microscopy. (A–C) Representative live-cell confocal microscopy images of Saos-2 cells transfected with a NT (RISC-Free) siRNA (A), VPS39-targeting siRNAs (B), or TGFBRAP1-targeting siRNAs (C). (D) Pearson correlation coefficients between GFP markers and 2^SiR^ in non-targeting siRNA control cells, VPS39, and TGFBRAP1 knockdown cells. Scale bars = 5 μm.

## DISCUSSION

Unlike many small molecules, most proteins and peptides with therapeutic potential must circumnavigate a complex journey to access the cytosol or nucleus of mammalian cells. There is compelling evidence that this journey often involves endocytosis (86, 87), the natural cellular process by which extracellular material is taken up into vesicles formed from the plasma membrane. However, the pathway into the cytosol requires not just uptake into endocytic vesicles, but also endosomal release, and this second step is widely recognized as the bottleneck hindering the efficient delivery of peptidic materials into the cytosol and nucleus (18, 88-96). We discovered several years ago that CPMPs such as ZF5.3 (18, 97, 98) overcome this bottleneck to reach the cytosol with exceptional efficiency, both with (23) and without (19) an appended protein cargo. Here, we sought to understand the cellular processes that support the trafficking of CPMPs into the cytosol. First, using two complementary cell-based assays, we ruled out the most obvious mechanism of endosomal release–a partial or complete disruption of endosomal integrity. Next, using a spectrum of tools, including a genome-wide RNA interference (RNAi) screen, quantitative fluorescence correlation spectroscopy (FCS), and live-cell, multi-color confocal imaging, we identified *VPS39*—a gene encoding a subunit of the homotypic fusion and protein sorting (HOPS) complex—as a critical determinant in the trafficking of CPMPs to the cytosol. CPMPs neither inhibit nor activate HOPS activity; indeed, HOPS activity is essential for the cytosolic access of CPMPs as well as other CPPs and hydrocarbon-stapled peptides. Subsequent multi-color confocal imaging studies identify CPMPs within the endosomal lumen, particularly within the intraluminal vesicles (ILVs) of Rab7^+^ and Lamp1^+^ endosomes that are the products of HOPS-mediated fusion. These results suggest that CPMPs require HOPS to reach ILVs—an environment that serves as a prerequisite for efficient endosomal escape.

The activity of the HOPS complex has also been implicated in the cytosolic trafficking of numerous bacterial, fungal, and viral pathogens, including coronavirus (CoV) (99, 100), the Ebola virus (101), *Fusarium graminearum* (102), *Candida albicans* (103, 104), *Cryptococcus neoformans* (105), and *Aspergillus nidulans* (106). In particular, although their genomes lack sequences encoding a canonical penta-arg motif, both CoV and Ebola require HOPS during the later steps of their postulated mechanism of cell entry: after the viral particle has been endocytosed but before it reaches the cytosol. Passage of CoV into the cytosol requires processing of viral proteins by lysosomal proteases to enable fusion with lysosomal membranes and release into the cytosol; HOPS depletion prevents CoV from reaching the lysosome and thus fusion and release cannot occur (100). Passage of Ebola virus is regulated by HOPS in a similar manner: Ebola viral particles are unable to fuse with the lysosomal membranes in HOPS-deficient cells (101). It is important to note that, in all cases where HOPS is a required cytosolic entry factor, no direct interactions between HOPS and the pathogen have been identified. HOPS operates in a supporting role by allowing the pathogen to escape during HOPS-mediated endosomal fusion, or by providing the pathogen with access to a favorable environment for escape, fully consistent with the model proposed for the cytosolic trafficking of CPMPs and hydrocarbon-stapled peptides (model 3).

Significant experiments performed *in vitro* and in cultured cells provide additional support for an endosomal escape pathway that relies on the fusogenic activity of ILVs (26, 50, 78, 107). Experiments performed *in vitro* indicate that Tat_48–60_ translocates efficiently across liposomal membranes whose lipid components resemble those of late endosomal ILVs (lysobisphosphatidic acid (LBPA)/1,2-dioleoyl-*sn*-glycero-3-phosphocholine (PC)/1,2-dioleoyl-*sn*-glycero-3-phosphoethanolamine (PE), 77:19:4), but not across those that resemble the late endosomal membrane (PC/PE/phosphatidylinositol (PI)/LBPA, 5:2:1:2), and not at all across those whose lipids resemble the plasma membrane (PC/PE/sphingomyelin (SM)/cholesterol, 1:1:1:1.5) (107). Similarly, a disulfide-bonded dimer of Tat (dfTat) induces leakage of the green fluorophore calcein from liposomes composed of ILV-like lipids (LBPA/PC/PE, 77:19:4), but not from liposomes that mimic EEs or the plasma membrane (PC:PE:cholesterol, 65:15:20) (50). The nuclear delivery of dfTat (as estimated by counting cells with nucleolar fluorescence staining from dfTat) is also significantly reduced in cells pre-incubated with an antibody against LBPA. LBPA is highly abundant in ILVs (108, 109) and its fusogenic activity is thought to be responsible for ILV back-fusion with endosomal membranes (110). Finally, certain cyclic CPPs, such as CPP12 (**5**), induce inward and outward membrane budding when added to giant unilamellar vesicles (GUVs) composed of LE lipids (PC/PE/PI/LBPA, 5:2:1:2) (26). The observed budding behavior is reminiscent of the dynamic nature of multivesicular LEs in live cells, for which ESCRT-dependent ILV inward budding and exosome outward budding are tightly connected to the unique LBPA-enriched lipid mixture observed in LEs (111). Finally, cell-permeant phosphorothioate-modified antisense oligonucleotides (PS-ASO) also colocalize with LBPA-containing ILVs inside LEs, and knockdown of ALIX protein—known to cause a reduction in cellular LBPA levels (110, 112, 113)—reduces cytosolic delivery of PS-ASO (78). Together, these data provide additional support for a model for cytosolic trafficking that requires HOPS to reach a favorable environment for endosomal escape: LEs and LYs containing ILVs enriched in the fusogenic lipid LBPA. The identification of ILVs as a portal for passing proteins into the cytosol will aid the development of next-generation biologics that overcome the limitations imposed by cellular membranes.

## MATERIALS AND METHODS

See Supplementary Information.

## Acknowledgments

This work was supported by US National Institutes of Health (NIH) grants R01 GM74756 and CA 170741 (to A. Sc.). A. St. is grateful to the Howard Hughes Medical Institute for an International Student Research Fellowship. We are all grateful to Elizabeth Rhoades and Xiaohan Li for initial help with FCS, to the Yale Center for Molecular Discovery for assistance with the RNAi screen, and to Dr. Andreas Ernst for helpful comments on the manuscript.

## Author Contributions

A. St, J.R.L., R.F.W., and A. Sc. designed research; A. St., J.R.L., R.F.W., and S.B. performed research; A. St., J.R.L., R.F.W., and A. Sc. analyzed data and wrote the manuscript.

## Supplementary Information

### Reagents, Chemicals, & Experimental Model Systems

#### Cell culture

Opti-MEM I reduced serum medium (Gibco, #31985). McCoy’s 5A medium with phenol red, without L-glutamine (Sigma, #M8403). McCoy’s 5A medium without phenol red (HyClone, #SH30270). Dulbecco’s modified eagle medium (DMEM), high glucose (Gibco, #11965). DMEM without phenol red, high glucose, with HEPES (Gibco, #21063). Dulbecco’s phosphate buffered saline (DPBS) (Gibco, #14190). Fetal bovine serum, heat inactivated (Sigma, #F4135). Penicillin streptomycin (pen strep) (Gibco, #15140). GlutaMAX supplement (100X) (Gibco, #35050). Sodium pyruvate (100 mM, 100X) (Gibco, #11360). TrypLE express (without phenol red) (Gibco, #12604). Fibronectin from bovine plasma (Sigma, #F1141). Lipofectamine 3000 transfection kit (Invitrogen, #L3000). Lipofectamine RNAiMAX reagent (Invitrogen, #13778).

#### Chemicals

Sucrose, BioUltra, for molecular biology (Sigma, #84097). Dextran, Alexa Fluor 488, 10,000 MW, anionic, fixable (Invitrogen, #D22910). Lissamine rhodamine B ethylenediamine (Thermo Fisher Scientific, #L2424). Lissamine rhodamine B sulfonyl chloride (Acros Organics, #413230010). Alexa Fluor 594 hydrazide (Life Technologies, #A10438). Hoechst 33342, trihydrochloride, trihydrate (Molecular Probes, #H3580). (*R*)-*N*-Fmoc-2-(7’-octenyl)alanine (Fmoc-R8-OH, #OK-UA-09222) and (*S*)-*N*-Fmoc-2-(4’-pentenyl)alanine (Fmoc-S5-OH, #OK-UA-09216) (Okeanos Tech Jiangsu Co.). Silicon rhodamine (SiR)-carboxyl was prepared as described previously (1).

#### Oligonucleotides

siRNAs and RT-qPCR primers. See Table S3.

#### Expression Plasmids

The DNA sequences for human galectins (hGal) 3 and 8 were cloned into the 3rd generation lentiviral eGFP expression vector pLenti CMV GFP Puro. The sequence for hGal3 was excised from plasmid pEGFP-hGal3 using restriction enzymes BsrGI-HF (New England Biolabs, #R3575S) and SalI (New England Biolabs, #R0138S) and ligated into the pLenti vector backbone digested with the same restriction enzymes using T4 DNA ligase (New England Biolabs, #B0202S) resulting in the plasmid pLenti CMV eGFP-hGal3 (this paper). The sequence of hGal8 (isoform A, 359 amino acids) was ordered as a codon-optimized gBlock gene fragment from Integrated DNA Technologies (IDT) with overhangs containing restriction enzyme sites for BsrgI-HF and SalI. After double digest and PCR cleanup, this fragment was ligated into the linearized pLenti CMV GFP puro vector using T4 ligase resulting in the plasmid pLenti CMV eGFP-hGal8 (this paper). For lentiviral particles, VSV-G envelope expressing plasmid pMD2.G and 2nd generation lentiviral packaging plasmid psPAX2 were used. pLenti CMV GFP Puro was a gift from Eric Campeau and Paul Kaufman (Addgene plasmid #17448) (2). pEGFP-hGal3 was a gift from Tamotsu Yoshimori (Addgene plasmid #73080) (3). pMD2.G and psPAX2 plasmids were a gift from Didier Trono (Addgene plasmids #12259 and #12260).

#### Bacterial Strains

For DNA plasmid amplification *Escherichia coli* (*E. coli*) strain XL10-Gold (Agilent Technologies, #200315) was used and cultured in LB medium.

#### Cell Lines

All mammalian cell lines were purchased from the American Type Culture Collection. Human osteosarcoma (Saos-2) (ATCC, HTB-85) cells were cultured in McCoy’s 5A without L-glutamine supplemented with 15% fetal bovine serum (FBS), 1x GlutaMAX, sodium pyruvate (1 mM), penicillin (100 units/mL), and streptomycin (100 μg/mL). Human embryonic kidney (HEK) 293T cells were cultured in DMEM supplemented with 10% FBS, L-glutamine (2 mM), penicillin (100 units/mL), and streptomycin (100 μg/mL). Cultures were maintained at 37 °C in a humidified atmosphere at 5%, CO_2_.

### Supplementary Methods

#### M1. Synthesis of Peptides and Miniature Proteins

##### Automated Peptide Synthesis

All peptides and miniature proteins were synthesized on H-PAL ChemMatrix or H-Rink ChemMatrix resin on a 50 μmol scale to generate products carrying C-terminal amides. All peptides and miniature proteins were synthesized on an Initiator+ Alstra synthesizer (Biotage) using microwave acceleration. Prior to the first amino acid coupling, the resin was swelled in *N*,*N*-dimethylformamide (DMF) (4.5 mL) at 70 °C for 20 minutes (min). For standard amino acid couplings, DMF (4.5 mL), *N*,*N*,*N*′,*N′*-tetramethyl-*O*-(1*H*-benzotriazol-1-yl)uronium hexafluorophosphate (HBTU) (5 equiv.), 1-hydroxybenzotriazole hydrate (HOBt) (5.0 equiv.), *N*,*N*-diisopropylethylamine (DIEA) (10.0 equiv.), and Fmoc-protected amino acid (5.0 equiv.) were subjected to microwave-assisted standard amino acid couplings (75 °C for 5 min). All arginine, cysteine, and histidine residues were coupled at 50 °C. All arginine residues were coupled twice. To couple olefinic residues, DMF (4.5 mL), PyClocK (3.75 equiv.), Fmoc-protected olefinic amino acids (Fmoc-S_5_-OH or Fmoc-R_8_-OH, 3.75 equiv.) were combined and allowed to stir at room temperature (RT) for 2 hours (h). Residues following olefinic amino acids were coupled twice. Fmoc deprotections were performed using 20% piperidine in DMF (4.5 mL) with microwave assistance (70 °C for 3 min, twice). For peptides containing aspartate or glutamate, 0.1 M HOBt was added to the deprotection solution to minimize aspartimide formation. The resin was washed thoroughly with DMF (four times, 4.5 mL/wash) between each coupling and deprotection step. Following synthesis, the resin was transferred to a custom glass reaction vessel containing a stir bar. Both vessel and stir bar were coated with SigmaCote before the resin was transferred.

##### Peptide Modifications: N-terminally SDex-Labeled Peptides

**1**^Dex^ and **2**^Dex^ CPMPs were synthesized and purified as described before (4). Briefly, Boc-Lys(Fmoc)-OH was coupled onto the N-terminus of a resin-bound peptide in a custom glass vessel using the peptide coupling conditions described above. The Fmoc-protected Lys sidechain was deprotected with 20% piperidine in DMF containing 0.1 M HOBt. Following Fmoc deprotection, dexamethasone-21-thiopropionic acid (SDex) (2.5 equiv.), 1-hydroxy-7-azabenzotriazole (HOAt) (2.5 equiv.), 1-[Bis(dimethylamino)methylene]-1*H*-1,2,3-triazolo[4,5-*b*]pyridinium 3-oxid hexafluorophosphate (HATU) (2.5 equiv.), DIEA (10.0 equiv.) and 2,6-lutidine (7.0 equiv.) in DMF (3.0 mL) were added to the resin. The resin-containing glass vessel was shaken for 18 h at RT on an orbital shaker. The resin was then thoroughly washed with DMF, dichloromethane (DCM), and methanol (MeOH), after which it was dried under a stream of nitrogen overnight.

##### Peptide Modifications: Hydrocarbon-Stapled Peptides

Hydrocarbon-stapled peptide variants of SAH-p53-8 (**3**) bearing an N-terminal acetyl group or an N-terminal lissamine rhodamine B linked at a Lys side chain were synthesized and purified as described before (5, 6). Briefly, to couple olefinic residues, DMF (4.5 mL), PyClocK (3.75 equiv.), and Fmoc-protected olefinic amino acids (Fmoc-S_5_-OH or Fmoc-R_8_-OH, 3.75 equiv.) were combined and allowed to stir at RT for 2 h. Residues following olefinic amino acids were coupled twice. Following automated peptide synthesis, resin-bound stapled peptides were washed three times with dichloroethane (DCE) (10 mL) and cyclized on resin in a custom glass vessel, immediately prior to acetylation or rhodamine labeling, using Grubbs Catalyst I (20 equiv.) in DCE (4 mL) for 2 h under a constant stream of nitrogen. This cyclization step was repeated an additional time.

##### Peptide Modifications: N-Terminal Acetylation

N-terminally acetylated peptides were generated by treating the terminally Fmoc-deprotected resin-bound peptide with a mixture containing acetic anhydride, DIEA, and DMF (2 mL, in a 85:315:1600 volumetric ratio) and stirring at RT for 45 min in a custom glass vessel.

##### Peptide Modifications: N-Terminal Lissamine Rhodamine B Labeling

To label polypeptides with lissamine rhodamine B, Boc-Lys(Fmoc)-OH or—for stapled peptides—Fmoc-β-Ala-OH was appended to the N-terminus of the respective sequence by combining the N-terminally deprotected resin with 7-azabenzotriazol-1-yloxy)trispyrrolidinophosphonium hexafluorophosphate (PyAOP) (5 equiv.), HOAt (5 equiv.), and DIEA (10 equiv.) in DMF (4 mL) under microwave irradiation in a MARS-5 microwave-accelerated system (CEM Corporation) for 20 min or at RT without microwave irradiation for 45 min. Then, the N-terminal residue was deprotected twice with 25% piperidine in DMF (5 mL) for 10 min each at RT. Following deprotection, the resin was washed thoroughly with alternating DMF (5 mL) and DCM (5 mL), three times, and then with DMF (5 mL), six times. Finally, the peptide synthesis vessel was purged with nitrogen and the resin was washed an additional five times with anhydrous DMF (4 mL) under nitrogen atmosphere. Lissamine rhodamine B sulfonyl chloride (10 equiv.) was solubilized in anhydrous DMF (3 mL) and added directly to the resin after the final anhydrous DMF wash, followed by the addition of anhydrous DIEA (10 equiv.). The reaction vessel was purged with nitrogen, sealed, and covered with aluminum foil to minimize light exposure. The labeling reaction was occur overnight at RT while shaking on an orbital shaker. After labeling, the resin was washed thoroughly with alternating DMF (10 mL) and DCM (10 mL), three times, then extensively with DMF until no excess lissamine rhodamine B was observed in the wash solution. To shrink the resin, it was washed with MeOH (5 mL) twice. Finally, the resin was dried under a stream of nitrogen overnight.

##### Peptide Modifications: N-to-C Cyclization

N-to-C cyclized peptide CPP12 (5) was synthesized and purified as described before (7). Briefly, the cyclization of the peptide was achieved by incorporating a main chain alloc-protected glutamic acid residue at the C-terminus of the peptide. The allyl group was removed selectively by treatment with Pd(PPh_3_)_4_ (0.1 equiv.), phenylsilane (10 equiv.) in anhydrous DCM (three times, 15 min each) releasing the C-terminal carboxylate. After allyl deprotection, the N-terminal Fmoc group was removed with piperidine (25%) in DMF and the peptide was cyclized by treatment with (benzotriazol-1-yloxy)tripyrrolidinophosphonium hexafluorophosphate (PyBOP) (5 equiv.), HOBt (5 equiv.) and DIEA (10 equiv.) in DMF overnight (16 h).

##### Peptide Cleavage from Resin

Once peptide synthesis was complete, the resin was washed thoroughly with alternating DMF (10 mL) and DCM (10 mL), three times, and with MeOH (5 mL), after which it was dried overnight under nitrogen. The peptide was cleaved from the resin using 4 mL (per 50 μmol) of a cocktail containing trifluoroacetic acid (TFA) (88%), phenol (5%), water (5%), and triisopropylsilane (TIPS) (2%) for 3 h at RT. For peptides containing cysteines, the cocktail consisted of TFA (81.5%), thioanisole (5%), phenol (5%), water (5%), ethanedithiol (2.5%), and TIPS (1%). SDex-labeled peptides were cleaved with a cocktail consisting of TFA (92.5%), water (2.5%), TIPS (2.5%), and 3,6-dioxa-1,8-octanedithiol (2.5%). Cleaved peptides were precipitated in diethyl ether (40 mL, chilled to –80 °C), pelleted by centrifugation, redissolved in a mixture of water and acetonitrile (ACN) (max. 20% ACN), frozen, lyophilized to dryness, and reconstituted in 1–2 mL dimethyl sulfoxide (DMSO) prior to purification by high-performance liquid chromatography (HPLC).

##### Peptide Purification by HPLC

Peptide solutions in DMSO were filtered through nylon syringe filters (0.45 μm, 4 mm from Thermo Fisher Scientific) prior to HPLC purification. All peptides were purified using an Agilent 1260 Infinity HPLC system, a reverse phase Triaryl-C18 (YMC-Triaryl-C18, 150 mm x 10 mm, 5 μm, 12 nm) column (YMC America, Inc.) and eluent gradients of water in ACN containing 0.1% TFA. Peptides were detected at 214 and 280 nm. For lissamine rhodamine B (R)- and silicon rhodamine (SiR)-labeled peptides were also detected at 560 nm and at 645 nm, respectively. Peptide purity was verified using a Shimadzu Analytical ultra-performance liquid chromatography (UPLC) system (ES Industries, Shimadzu Corporation) and a C8 reverse phase (Sonoma C8(2), 3 μm, 100 Å, 2.1 x 100 mm) or a C18 reverse phase (Agilent Poroshell 120, 2.7 μm, 4.5 x 50 mm) analytical column. Analytical samples were eluted using solvent gradients of water in ACN containing 0.1% TFA over 20–25 min and were detected at 214, 280, and, for R- and SiR-labeled peptides, at 560 nm and 645 nm, respectively.

##### Mass Spectrometry

The molecular mass of each peptide was determined by liquid chromatography-mass spectrometry (LC-MS), using a Waters XEVO Q-TOF mass spectrometer equipped with an Acquity UPLC BEH C18 1.7 μm column (for Chemdraw structures and mass spectrometry data of all peptides see Figure S1 and Table S1, respectively).

##### Peptide Reconstitution, Concentration Determination, and Storage

Purified, unlabeled peptides containing tyrosine or tryptophan were dissolved in water and the concentration of the solution was determined using the extinction coefficient of the peptide at 280 nm (ε_280_), which was estimated using the ProtParam peptide properties calculator on the ExPASy Proteomics Server. For SDex-labeled peptides, concentrations were measured using an SDex extinction coefficient of 12,000 cm^−1^ M^−1^ at 242 nm in water, as described previously (4, 8). For R-labeled peptides, the extinction coefficient for lissamine rhodamine B diethylamine was determined in DPBS (pH 7.4) at 25 °C to be 112,000 ± 2,000 cm^−1^ M^−1^ at λ_max_(569 nm). This extinction coefficient was used for all rhodamine-labeled peptides. For unlabeled cyclic peptide CPP12, the extinction coefficient of naphthylalanine (3,936 cm^−1^ M^−1^ at λ_max_(280 nm)) was used (9). For SiR-labeled peptides, the reported extinction coefficient was used (1). Purified zinc finger CPMPs (**2**^UL^, **2**^Dex^, and **2**^R^) were dissolved in argon-purged 10 mM Tris buffer, pH 7.4. To reduce cysteines, DTT (2 equiv.) was added and the solution was left to react for 15 min. Then, ZnCl_2_ (2 equiv.) was added to the solution to induce the typical zinc finger fold. Zinc finger peptides were stored in solution at 4 °C. Repeated freeze-thawing cycles of zinc finger and aPP-based miniature proteins were avoided due to peptide aggregation and precipitation. All other peptides (**3**–**5**) were stored as a lyophilized powder or in solution at –20 °C and tolerated repeated freeze-thaw cycles. All peptides were routinely assessed for identity and purity by LC-MS and UPLC analyses.

#### M2. Fluorescence correlation spectroscopy (FCS)

##### Determination of the Confocal Volume by FCS

FCS measurements were perfomred on an LSM 880 Airyscan system NLO/FCS Confocal microscope (Zeiss) with a C-apochromat 40x N1.2 UV-VIS-IR Korr. water immersion objective (Zeiss), with a gallium arsenide phosphide (GaAsP) detector. Alexa 594 standards and lissamine rhodamine B-labeled peptides were excited at 561 nm, and the fluorescence filters were set to 570–660 nm. Measurements were made in 8-well chambered Lab-Tek 8-well coverglass slides (Thermo Fisher Scientific, #155411). *In vitro* standards with Alexa 594 were measured at 25 °C or 37 °C in Milli-Q water. *In cellulo* experiments were performed at 37 °C in serum-free DMEM without phenol red containing 25 mM HEPES (pH 7.2). Autocorrelation data were collected over 5-s intervals with 10 repeats. Prior to FCS measurements in cells, the microscope was calibrated using Alexa 594 dye. The coverglass thickness of each 8-well microscopy dish was measured with a digital micrometer (Mitutoyo) to adjust the correction collar of the C-apochromat 40x N1.2 water immersion objective to the correct thickness. The pinhole of the 561 nm laser was aligned in both the x and the y direction. A standard solution of Alexa 594 dye (100 nM) was measured at 37 °C in the same 8-well coverglass slides used for *in cellulo* FCS experiments with one well reserved for Alexa 594 standard (no fibronectin coating). Autocorrelations were fit to diffusion equations described below, using custom scripts Matlab (Version R2017a, MathWorks), as described before (5, 10).

Autocorrelation curves from *in vitro* measurements were fit to a 3D diffusion equation (Equation 1):

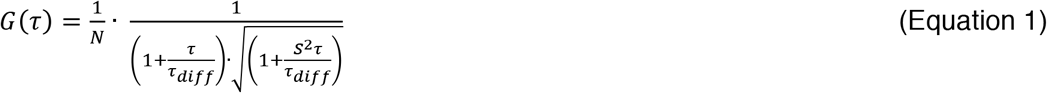

*N* is the average number of diffusing molecules in the effective confocal volume (*V_eff_*) and *τ_diff_* is the diffusion time, the average time a molecule takes to transit the laser focus. The shape factor *S* of the effective confocal volume *V_eff_* was determined from the fit of the autocorrelation function of Alexa 594 (12.5 nM) in water at 25°C (*S* = 0.2 ± 0.007) and was fixed for all subsequent analyses at *S* = 0.2. *V_eff_* was determined to be 0.66 ± 0.090 fl and was calculated according to Equations 2 and 3:

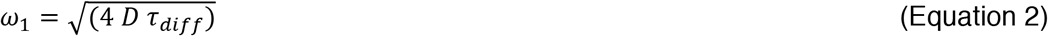

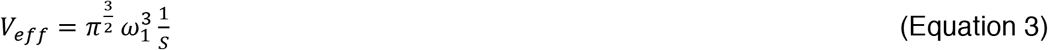

*D* is the diffusion coefficient of Alexa 594 at 37 °C (5.20 x 10^−6^ cm^2^ s^−1^) and *τ_diff_* is the measured diffusion time. The diffusion coefficient *D* of Alexa 594 at 37 °C was calculated using Equation 4:

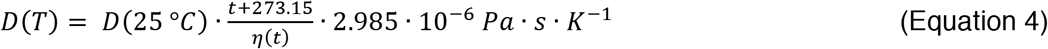

Where *t* = 37 °C, *D* of Alexa 594 at is 25 °C (3.88 x 10^−6^ cm^2^ s^−1^) (11), and the viscocity *η* of water at 37 °C is 0.6913 mPa s.

The final concentration *C* in the effective confocal volume *V_eff_* was calculated as follows (Equation 5):

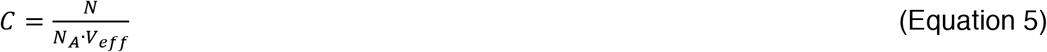

Where *N_A_* is Avogardro’s number (6.0221413 x 10^23^ mol^−1^).

Autocorrelation curves from *in cellulo* measurements were fit to an anomalous diffusion model:

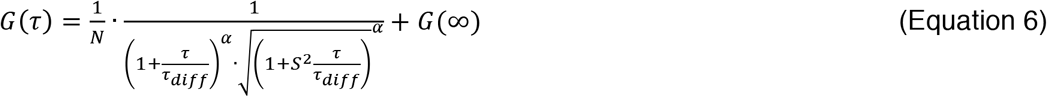

*G (∞)* represents the level of background autocorrelation at long time scales and *α* is the anomalous diffusion coefficient, which represents the degree to which diffusion is hindered over long distances (12).

The fit autocorrelation traces from live-cell measurements were then evaluated and filtered as described before (5). Briefly, for follow-up analysis, we selected traces displaying diffusion times (*τ_diff_*) of up to 10-fold of the observed value for the identical CPMP/CPP in buffer. We discarded traces that displayed poor signal with counts per molecule (cpm) below 1 kHz and/or low anomalous diffusion coefficients (α < 0.3) (13). With these parameters, we typically retained at least 75% of the collected data points.

##### Calculation and Measurement of Diffusion Coefficients of Peptides 1–5

The theoretical diffusion coefficient of a spherical, globular macromolecule can be determined from its respective molecular weight. First, its molecular mass (*m*) is converted to the hydrodynamic radius (*r*) with the following relationship (Equation 7):

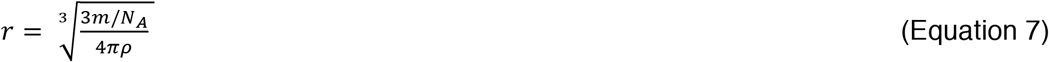

*Where ρ* is mean density, which is 1.2 g/cm^3^ for proteins.

Second, the theoretical diffusion coefficient (*D*) is calculated from *r* with the Stokes-Einstein equation (Equation 8):

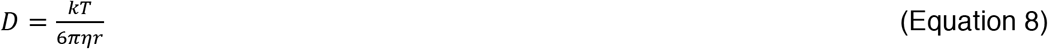

*k* is the Boltzmann constant (1.381 × 10^−16^ erg × K^−1^), *η* the viscosity of water at 25 °C (0.891 g m^−1^ s^−1^), and *T* is the absolute temperature.

The diffusion coefficients of peptides **1**^R^–**5**^R^ was measured *in vitro* at 25 °C in Milli-Q water (Table S2).

#### M3. Cell-based assays

##### Preparation of eGFP-hGal3 and eGFP-hGal8 Lentiviral Particles for Transduction

HEK 293T cells (ATCC, CRL-3216) were cultured in complete growth medium (DMEM containing 4.5 g/L glucose, phenol red, 10% FBS, 5 mM sodium pyruvate, without antibiotics). Lentiviral particles were prepared in HEK 293T cells using Lipofectamine 3000 (Invitrogen, #L3000) transfection reagent according to the manufacturer’s protocol. Briefly, 293T cells were seeded (1.2 Mio/well) into tissue-culture treated 6-well dishes in 2 mL packaging medium (Opti-MEM I containing 5% FBS and 1 mM sodium pyruvate) and left to adhere overnight. For transfection, a 1:1 mixture of solution A containing Lipofectamine 3000 in Opti-MEM I (250 μL) and solution B containing P3000 reagent, psPAX2 (3 equiv. 1688 ng/well), pDM2.G packaging plasmids (536 ng/well), and either eGFP-hGal3 or eGFP-hGal8 in pLenti CMV GFP puro plasmids (750 ng/well) in Opti-MEM I (250 μL), was prepared and incubated for 10 min at RT. The 1:1 mixture was added dropwise to the plated HEK 293T cells (total 500 μL/well) in 6-well plates. The medium was replaced with packaging medium 6 h after transfection. Lentiviral particles were harvested 24 h after transfection by carefully collecting the 2 mL cell supernatant from each well. Wells were replenished with fresh packaging medium, which was collected a second time 52 h post transfection. All lentivirus-containing fractions were stored at 4 °C for two days, combined, filtered through a 0.45 μm pore syringe filter, and concentrated using spin-filter concentrators (Amicon Ultra-15, 100 kD MWCO, EMD Millipore, #UFC910024). The concentrated lentiviral stock solutions were split into aliquots and frozen at –80 °C until used. To determine the viral titer, cells were transduced with increasing amounts of the lentiviral solution and the fluorescence intensity at 509 nm was measured and compared to untreated cells using an Attune NxT flow cytometer (Life Technologies).

##### Galectin-Based Assay to Measure Endosomal Damage

To begin, 8-well microscopy slides (Thermo Scientific, #155411) were coated with fibronectin from bovine plasma in DPBS (1:100 dilution, 200 μL/well) for 1 h at 37 °C (200 μL/well). The fibronectin solution was removed by aspiration and the dishes were washed with DPBS (3x) before use. Saos-2 cells were seeded (15,000 cells/well) into the fibronectin-coated dishes in complete McCoy’s 5A medium without phenol red (200 μL/well, 15% FBS, 1 mM sodium pyruvate, without P/S) and left to adhere overnight (16 h). Cells were transduced with eGFP-hGal3 or eGFP-hGal8 at a lentiviral particle concentration determined to yield >50% transduced cells as measured by by flow cytometry compared to a mock-transduced cell population. After a 16-h transduction period, the medium was removed, cells were washed with DPBS (2x), and incubated with pre-warmed clear McCoy’s 5A medium without phenol red (without FBS, without sodium pyruvate, without P/S) containing medium only (control), Lipofectamine RNAiMAX (16 μL/mL), LLOMe (1 mM), or CPP/CPMPs **1**–**5** (600 nM) for 1 h at 37 °C, 5% CO_2_. Cells were washed with pre-warmed DPBS (3x) and incubated with HEPES-containing clear DMEM without phenol red (25 mM HEPES, without P/S, without FBS, without sodium pyruvate) for 30 min at 37°C, 5% CO_2_, after which they were imaged by confocal microscopy at 37 °C on a Zeiss laser scanning microscope (LSM 880). For galectin recruitment assays, careful attention was paid not to overexpose cells to allow detection of even small amounts of galectin recruitment to endosomal membranes. The endosomal recruitment coefficient (ERC) was determined by measuring the area of punctate fluorescence in each cell, dividing it by the overall area of the same cell, and then multiplying it by 10,000 (Figure S2C) using ImageJ (14). The ERC represents the %-area of punctate fluorescence in each cell multiplied by 100.

##### CellTiter-Glo Luminescent Cell Viability Assay

Saos-2 cells were seeded into 96-well plates (Corning, #3603; 3000 cells/well, 100 μL/well) in complete McCoy’s 5A medium without phenol red (10% FBS, without P/S, 1 mM sodium pyruvate) and left to adhere overnight (16 h). The next day, the medium was replaced with clear McCoy’s 5A medium without phenol red (100 μL/well, without FBS, without P/S, without sodium pyruvate) and peptides **1**^UL^–**5**^UL^ were added in triplicate to each well at increasing concentrations (0.0, 0.3, 0.6, 1.2, 2.4, 4.8, 9.6, 19.2 μM). Cells were left to incubate at 37°C, 5% CO_2_ for 4 h. CellTiter-Glo (Promega, #G7571) luminescent substrate was added to each well (50 μL/well). The plates were orbitally shaken for 15 min and the resulting luminescence was recorded with a Biotek Synergy 2 plate reader.

##### FCS-Based Assay to Measure Endosomal Damage Using Lys9^R^

One day prior to the experiment, Saos-2 cells were seeded in tissue culture-treated 6-well plates (100,000 cells/well) in complete McCoy’s 5A medium without phenol red (2 mL/well, 15% FBS, 1 mM sodium pyruvate, without P/S). The day of the experiment, the medium was removed and cells were washed twice with DPBS. Cells were incubated with Lys9^R^ (600 nM) in clear McCoy’s 5A without phenol red (1 mL/well, without FBS, without sodium pyruvate, without P/S) for 30 min at 37 °C, 5% CO_2_. While cells were incubating, an 8-well microscopy slide (Thermo Scientific, #155411) was coated with fibronectin from bovine plasma in DPBS (1:100 dilution, 200 μL/well) for 1 h at 37 °C (200 μL/well). The fibronectin solution was removed by aspiration and the dish was washed with DPBS (3x) before use. After the 30-min incubation period, cells were washed thoroughly with DBPS (3x) and incubated with increasing concentrations of Lipofectamine RNAiMAX (0, 4, 8, 12, 16 μL/mL medium), LLOMe (0, 0.125, 0.25, 0.5, 1.0 mM), or unlabeled peptides **1**^UL^–**5** ^UL^ (300, 600, 1200, 2400 nM) in clear McCoy’s 5A without phenol red for 1 h at 37 °C, 5% CO_2_. Nuclei were labeled with Hoechst 33342 (300 nM) for 5 min at the end of the 1- h incubation period by adding the nuclear stain directly to the reagent-containing medium. To remove exogenously bound peptide, cells were washed with DBPS (3x) and treated with TrypLE (500 μL/well) for 10 min until all cells were lifted from the culture dish. The cells were transferred to 15-mL Falcon tubes, each well was rinsed twice with complete McCoy’s 5A without phenol red (twice, 1 mL each), and the cell suspensions, including media used for rinsing, were pooled and pelleted at 200 *g* for 3 min. The supernatant was removed and cells were resuspended in HEPES-containing clear DMEM without phenol red (1 mL, 25 mM HEPES, without FBS, without sodium pyruvate, without P/S) to wash the pellet. The cells were pelleted again at 200 *g* for 3 min. The supernatant was removed and the cells were resuspended in HEPES-containing clear DMEM without phenol red (600 μL). Of this suspension, one third (200 μL) was plated into the prepared fibronectin-coated microscopy dish. The cells were left to adhere at 37 °C, 5% CO_2_ for 20 min before FCS experiments were performed.

Before beginning experiments at the LSM 880, the live-cell chamber was equilibrated to 37 °C, and the 561-nm (0.5%) laser line was warmed up, both for approximately 30 min. After *in vitro* calibration with Alexa 594, confocal images of Saos-2 cells were obtained. FCS measurements were obtained by positioning the 561-nm laser’s crosshair in the nucleus or the cytosol of Saos-2 cells. A minimum of 20 cells was measured for each experimental condition. To minimize background fluorescence and artifacts from diffusion, the laser’s confocal volume was placed in a cytosolic area devoid of bright endosomes. Measurements in the nucleus were less prone to diffusional artifacts. For each FCS measurement, ten consecutive five-second time intervals were recorded. These ten traces were converted to their corresponding autocorrelation curves using a custom Matlab script (5). Low quality traces were discarded as described above, the remaining traces were averaged, and the averaged trace was fit to an anomalous diffusion model (Equation 6). From the fit of the averaged autocorrelation curve, we obtained the number of Lys9^R^ molecules in the focal volume of the laser. This number of Lys9^R^ molecules was converted to the effective cytosolic or nuclear Lys9^R^ concentration using the laser’s measured focal volume, as described in equations 2–5 above. Evaluation of the correlation curves revealed that cells generally displayed wide variation in the maximal autocorrelation signal, indicating an equally wide variation in the concentration of each molecule in the cytosol.

##### FCS-Based Control Experiment With Lys9^UL^

Except for two modifications, the same experimental protocol was followed as for the experiment with labeled Lys9^R^, described above. Instead of Lys9^R^, cells were incubated with or without Lys9^UL^ (600 nM), and instead of unlabeled peptides **1**–**5**, cells were incubated with RhoB-tagged peptides **1**^R^– **5**^R^ (300, 600, 1200, 2400 nM) (Figure S4A).

##### Cytosolic Fractionation to Quantify Cytosolic Access of D-Arg8^R^ (4^R^) and CPP12^R^ (5^R^)

When we incubated Saos-2 cells with D-Arg8^R^ (**4**^R^) and CPP12^R^ (**5**^R^), and performed FCS experiments similar to the ones described above, we observed *in cellulo* diffusion times that were more than 25 times longer compared to diffusion times commonly observed for aPP5.3^R^ (**1**^R^), ZF5.3^R^ (**2**^R^), or SAH-p53-8^R^ (**3**^R^) (Table S2 and Figure S4B). An increase in diffusion time may be a sign of supramolecular aggregation or binding to intracellular factors, which renders FCS data analysis impossible. Therefore, we turned to our previously described cytosolic fractionation assay to calculate the cytosolic concentration of **4**^R^ and **5**^R^ relative to **2**^R^. We performed the same incubation protocol as described for the Lys9 FCS-based assays above, but with a larger population of Saos-2 cells. One day prior to the experiment, Saos-2 cells were plated in tissue culture-treated 100-mm dishes (1.5 × 10^6^ cells/dish) in complete McCoy’s 5A without phenol red (5 mL/dish, 15% FBS, 1 mM sodium pyruvate, without P/S). The next day, the medium was removed and cells were washed twice with DPBS. The cells were incubated with or without Lys9^UL^ (600 nM) in clear McCoy’s 5A without phenol red (5 mL/well, without FBS, without sodium pyruvate, without P/S) for 30 min at 37 °C, 5% CO_2_. After the 30-min incubation, cells were washed thoroughly with DBPS (3x), and incubated with increasing concentrations of RhoB-tagged peptides **1**^R^–**5**^R^ (300, 600, 1200, 2400 nM) in clear McCoy’s 5A medium without phenol red for 1 h at 37 °C, 5% CO_2_. To remove exogenously bound peptide, cells were washed with DBPS (3x) and treated with TrypLE (1 mL/dish) for 10 min until all cells were lifted from the culture dish. Cells were transferred to 15-mL Falcon tubes, each well was rinsed twice with complete McCoy’s 5A without phenol red (twice, 1 mL each), and the cell suspensions, including the media used for rinsing, were pooled and pelleted at 200 *g* for 3 min. The cells were washed with DPBS (1 mL) twice and pelleted at 200 *g* for 3 min. The cells were resuspended in 1 mL pre-cooled buffered isotonic sucrose buffer (290 mM sucrose, 10 imidazole pH 7.0, added immediately prior to use: 1 mM DTT and 1 cOmplete protease inhibitor cocktail per 10 mL buffer). A 20-uL aliquot was removed for counting, then the suspension was pelleted again at 200 *g* for 3 min. Saos-2 cells were counted using an automated cell counter (Auto T4, Cellometer). The supernatant was removed and cells were suspended in pre-cooled buffered isotonic sucrose to reach a concentration of 10,000 cells/μL. For each experimental condition, 1.5 x 10^6^ cells were suspended in 150 μL isotonic sucrose, transferred to a 0.5-mL microtube containing 1.4mm ceramic beads (Omni International, #19-626), and stored on ice for transport. Cells were homogenized using a Bead Ruptor 4 (Omni International) at speed 1 for 10 seconds. The homogenized cells were transferred to Beckman polycarbonate ultracentrifuge tubes (0.2 mL, Beckman Coulter, #343775) and sedimented using a Beckman tabletop ultracentrifuge at 350 K*g* for in an ultracentrifuge (TL-100; Beckman Coulter) for 30 min at 4 °C using a TLA-100 rotor (20 x 0.2 mL). The supernatant (= cytosolic fraction) was analyzed in a 96-well plate (NBS-treated, low-protein binding, Corning, #3651) on a fluorescence plate reader (infinite F500, Tecan) with excitation at 535 ± 25 nm and an emission at 590 ± 20 nm. The concentration of the protein conjugate in the cytosol was calculated using a calibration curve. For the calibration curve, known concentrations of peptides **2**^R^, **4**^R^, and **5**^R^ between 1.0 and 1000.0 nM were added to cytosolic extracts of the untreated negative control cells. For background subtraction, several wells containing cytosolic extracts from untreated cells were averaged. The average background was subtracted from each well.

##### Screen Optimization: Seeding Density and Growth Medium

To evaluate the effect of seeding density and the choice of RNAi growth medium on the TR of Saos-2(GIGT) cells, Saos-2(GIGT) cells were plated at various seeding densities in either 50 μL of complete growth medium (McCoy’s 5A, with pen/strep, 15% FBS) or transfection medium (complete growth medium/Opti-MEM/RNAi duplex buffer, 3:1:1) in 384-well plates (#1052, Nexus Biosystems). Saos-2(GIGT) cells were allowed to adhere and remain in the respective media for 48 h, after which the cells were serum-starved overnight (16 h), by replacing the media with 40 μL of serum-free DMEM. The following morning, cells were treated with DMEM containing either 1 *μ*M Dex or Dex-peptide conjugate (**1**^Dex^ or **2**^Dex^) for 30 min, after which cells were fixed with 4% paraformaldehyde for 20 min at RT, stained with the nuclear stain Hoechst 33342 (300 nM) for 20 min at RT, and imaged on an Opera high-throughput imaging system. Control cells containing no peptide were treated with media only. Robust TRs between 3.5–4.0 were detected for Dex-treated cells under all plating densities and media tested. For **1**^Dex^ and **2**^Dex^, TRs were higher in transfection media (around 3.0 for both) compared to complete media (around 2.0 for both). For RNAi screening, 2,500 cells per well were plated in transfection medium, as these conditions resulted in a high number of cells per well (150–250 per image) without overcrowding and the highest TRs for Dex alone (TR = 3.4 ± 0.40), dex-peptide conjugates **1**^Dex^ (TR = 3.0 ± 0.40), and **2**^Dex^ (TR = 3.0 ± 0.090) (Figure S5B).

##### Integration of Saos-2(GIGT) cells with High-Content RNAi Screening

High-Content RNAi screening was performed in 384-well format. To each well, siRNA duplex buffer (10 μL) with or without siRNA (20 nM), Opti-MEM (10 μL) containing Lipofectamine RNAiMAX (0.1 μL), and Saos-2(GIGT) cells (2500 per well) in complete McCoy’s 5A medium without phenol red (30 μL per well, 10% FBS, without antibiotics) were added using a Multidrop Combi reagent dispenser (Thermo Scientific). The outer four columns of each 384-well plate contained experimental controls, while the inner 20 columns contained variable siRNAs, with 4 duplexes targeting a single gene within each well (SMARTpool siRNA). Background GR*eGFP translocation ratios (TRs) were determined from column 24, which contained untreated GR*eGFP cells transfected with RISC-Free non-targeting siRNA, while **1**^Dex^ TRs were represented in columns 2 and 23. Plate-wide transfection efficiency was determined from column 1, which contained alternating buffer-only and Kif11 siRNA control wells. Knockdown of Kif11 results in mitotic arrest, phenotypically observable enlarged nuclei, and dead cells (15). The significant loss in cell number upon Kif11 transfection, relative to RISC-Free transfection, is commonly used as a measure of siRNA transfection efficiency during high-throughput screens. In a pilot screen, we identified that siRNAs targeting the *SYNJ2BP* gene enhanced GR*eGFP translocation in the presence of **1**^Dex^ and yields a hit-like phenotype. We therefore included SYNJ2BP siRNA in columns 2 and 23 of every plate, to provide a representation of a hit candidate. Cells were reverse-transfected as follows: siRNA (20 nM) in sterile duplex buffer (10 μL) were first dispensed into each well, followed by Lipofectamine RNAiMAX (0.1 μL) in Opti-MEM (10 μL). After siRNAs and Lipofectamine RNAiMAX were allowed to complex for 20 min, Saos-2(GIGT) cells in McCoy’s 5A without phenol red (30 μL, 10% FBS, without antibiotics) were dispensed into each well. Saos-2(GIGT) cells were allowed to settle for 30 min at RT, prior to placement into a 37 °C humidified incubator with 5% CO_2_ for 56 h. After 56 h of reverse-transfection, cells were serum-starved for 16 h by replacing the media in each well with McCoy’s 5A medium, without phenol red (30 μL, without FBS, without antibiotics).

##### High-Content Imaging of Saos-2(GIGT) Cells

After serum starvation, 10 μL clear McCoy’s 5A medium supplemented with ligand at 5X concentration was overlaid onto wells, and Saos-2(GIGT) cells were incubated in the presence of Dex-peptide conjugates for 30 min. Following treatment, cells were fixed with 4% paraformaldehyde for 20 min at RT, washed twice with DPBS, stained with Hoechst 33342 (300 nM) for 45 min at RT, washed twice more, and imaged on an Opera high-throughput spinning disk confocal microscope.

Next, the reproducibility of GIGT assay was tested across two identical panels of 320 siRNAs targeting genes across the Dharmacon Human Genome Library. Non-treated Saos-2(GIGT) cells transfected with non-targeting siRNA displayed a mean TR of 1.2 ± 0.1 while identical cells treated with 1 *μ*M **1**^Dex^ for 30 min displayed a mean TR of 2.4 ± 0.3. Across a panel of 320 test siRNAs, this system yielded a mean signal-to-background of 2.3, coefficient of variation (CV) of 11.2%, Z-factor of 0.30, and Pearson R correlation coefficient of 0.80 between two replicate plates (Figure S5C). Collectively, these results underscore the adequacy of the GIGT system towards high-content RNAi screening.

Screening system: Opera (PerkinElmer Life and Analytical Sciences) using a 20x 0.45 NA lens. GR*-eGFP fluorescence was detected using a solid state 488 nm laser and a 540/75 band-pass filter, while Hoechst 33342 was detected using a 405 nm laser and a 450/50 band-pass filter. Three images were taken per well, and each image typically contained ~200–300 cells. GR*-eGFP TRs were determined using Acapella high content imaging and analysis software (Perkin Elmer).

##### Plate Performance (Z-Factor Analysis)

Individual plate performance was determined based on the Z-factor (Z′) between positive RISC-Free transfected Saos-2(GIGT) cells treated with 1 *μ*M **1**^Dex^ and negative cells transfected with RISC-Free and left untreated, which was calculated using Equation 9:

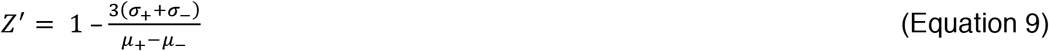

Where *μ_+_* and *σ_+_* refer to the mean and standard deviation of Saos-2(GIGT) cells transfected with RISC-Free siRNA and treated with 1 *μ*M **1**^Dex^, while *μ_–_* and *σ_–_* refer to the mean and standard deviation of nontreated cells transfected with RISC-Free non-targeting siRNA.

##### RNAi Translocation Data Normalization

Raw TRs were converted to normalized percent effect values using Equation 10. Data were normalized with respective controls per-plate (16).

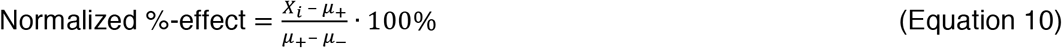

Where *X_i_* is the mean GR*-eGFP TR of the respective RNAi knockdown, *μ_+_* is the mean TR of cells transfected with RISC-Free siRNA and treated with 1 *μ*M **1**^Dex^ and *μ_–_* is the mean TR of nontreated cells transfected with RISC-Free siRNA. The normalized %-effect of Saos-2(GIGT) cells transfected with RISC-Free siRNA and treated 1 *μ*M **1**^Dex^ was defined as 0%, while that of nontreated Saos-2(GIGT) cells transfected with RISC-Free siRNA was defined as 100%.

##### Calculation of Z-Score

The Z-score, or the number of standard deviations between each knockdown and the mean TR of Saos-2(GIGT) cells transfected with RISC-Free siRNA and treated with 1 *μ*M **1**^Dex^ (*μ_+_*), were calculated using Equation 11:

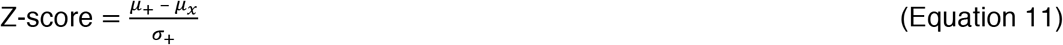

Where *μ_+_* is the mean TR of Saos-2(GIGT) cells transfected with RISC-Free siRNA and treated with 1 *μ*M **1**^Dex^, and *σ_+_* is the standard deviation of this population. *μ_x_* refers to the mean TR of the respective RNAi knockdown.

##### RNAi Screen Hit Assignment Using Strictly Standardized Mean Difference (SSMD)

To assign hits within the data set, normalized %-effect values from triplicate knockdowns were converted to strictly standardized mean difference (SSMD or β) values with Equation 12 (17):

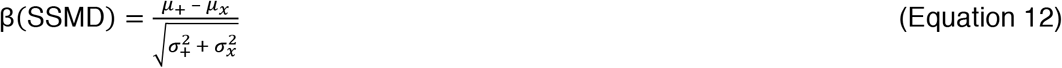

Where *μ_x_* is the mean %-effect of Saos-2(GIGT) cells transfected with gene-specific siRNAs and treated with 1 *μ*M **1**^Dex^ and *σ_x_* refers to the standard deviation of this population. *μ_+_* and *σ_+_* refer to the mean %-effect of Saos-2(GIGT) cells transfected with RISC-Free siRNA and treated 1 *μ*M **1**^Dex^ and to the standard deviation of this population.

##### siRNA Reverse Transfection for Hit Validation by FC and FCS

This protocol was adapted from the manufacturer’s protocol and optimized for 6-well plates. It was used for RT-qPCR, FC and FCS experiments. Saos-2 cells (passage 5–20) were lifted with TrypLE for 10 min at 37 °C, 5% CO_2_, and pelleted in complete McCoy’s 5A medium without phenol red (15% FBS, 1 mM sodium pyruvate, without P/S) at 200 *g* for 3 min. Cells were resuspended in the same media and counted. For each well to be transfected, the appropriate amount of siRNA (100 nM final concentration, Table S3) was diluted in Opti-MEM (200 μL) and vortexed briefly to mix (solution A). In parallel, for each well to be transfected, Lipofectamine RNAiMAX (5 μL) was diluted in Opti-MEM (200 μL), and vortexed briefly to mix (solution B). Solutions A and B were mixed in a 1:1 ratio, vortexed briefly to mix, and incubated for 5 min at RT. To each well, Saos-2 cells (80,000 cells/well) were added in complete McCoy’s 5A medium without phenol red (1.6 mL, 15% FBS, 1 mM sodium pyruvate, without P/S). The RNAi duplex-Lipofectamine RNAiMAX complexes (400 μL) were added dropwise to the cell solution in each well. Cells were mixed gently by rocking the 6-well plate back and forth (final volume of medium/well = 2 mL). Cells were moved to the incubator (37 °C, 5% CO_2_). After 3.5 h, Saos-2 cells were washed 3 times with pre-warmed DPBS (4 mL/well, 3 min/wash) to remove Lipofectamine RNAiMAX. Media was replaced with fresh complete growth medium without phenol red (2 mL/well, 15% FBS, 1 mM sodium pyruvate, without antibiotics). Saos-2 cells were incubated for 72 h at 37 °C, 5% CO_2_ for optimal gene knockdown. Control cells were transfected with Lipofectamine RNAiMAX only, or with a non-targeting siRNA (RISC-Free) and Lipofectamine RNAiMAX. To assess transfection efficiency, Kif11 siRNA was used. Knockdown of Kif11 leads to mitotic arrest, which causes the cells to round up and die. This was used as a visual control for transfection efficiency.

##### Confirming Knockdown by RT-qPCR

Saos-2 cells were reverse-transfected in 6-well plates as described above. Total RNA was extracted using the RNeasy Mini kit (Qiagen, #74104), 48 or 72 h after transfection. RNA was reverse-transribed with Superscript III reverse transcriptase (Thermo Fisher Scientific) and gene-specific primers (IDT, PrimeTime qPCR primers, Table S3) according to the manufacturer’s protocol. For RT-qPCR, cDNA was amplified with SsoFast EvaGreen Supermix (Bio-Rad, Hercules, CA, USA) on a Bio-Rad CFX96 real-time PCR detection system. Each biological replicate was run in triplicate, assayed for the siRNA-targeted gene and compared to GAPDH serving as an endogenous reference gene (Figure S6A).

##### FC and FCS to Quantify Cytosolic Access

Saos-2 cells were reverse-transfected in 6-well plates as described above. The next day, the medium was removed and cells were washed twice with DPBS. Cells were incubated with peptides **1**^R^–**5**^R^ (600 nM) in clear McCoy’s 5A medium without phenol red (1 mL/well, without FBS, without sodium pyruvate, without P/S) for 30 min at 37 °C, 5% CO_2_. While cells were incubating, an 8-well microscopy slide (Thermo Scientific, #155411) was coated with fibronectin from bovine plasma in DPBS (1:100 dilution, 200 μL/well) for 1 h at 37 °C (200 μL/well). The fibronectin solution was removed by aspiration and the dish was washed with DPBS (3x) before use. Nuclei were labeled with Hoechst 33342 (300 nM) for 5 min at the end of the 30-min incubation period by adding the nuclear stain to the peptide-containing medium. To remove exogenously bound peptide, cells were washed with DBPS (3x) and treated with TrypLE (500 μL/well) for 10 min until all cells were lifted from the culture dish. Cells were transferred to 15-mL Falcon tubes, each well was rinsed twice with complete McCoy’s 5A medium without phenol red (twice, 1 mL each), and the cell suspensions, including media used for rinsing, were pooled and pelleted at 200 *g* for 3 min. The supernatant was removed and cells were resuspended in HEPES-containing clear DMEM without phenol red (1 mL, 25 mM HEPES, without FBS, without sodium pyruvate, without P/S) to wash the pellet. Cells were pelleted again at 200 *g* for 3 min. The supernatant was removed and cells were resuspended in HEPES-containing clear DMEM without phenol red (600 μL). Of this suspension, one third (200 μL) was plated into the prepared fibronectin-coated microscopy dish and cells were left to adhere at 37°C, 5% CO_2_ for at least 20 min before FCS experiments were performed. The remaining two thirds of the cell suspension were pelleted again at 200 *g* for 3 min, resuspended in DPBS (200 μL) and transferred to a polystyrene round bottom 96-well plate (#353077, Falcon) for flow cytometry analysis. Flow cytometry was performed on a BD Accuri C6 Flow Cytometer (excitation laser: 488 nm, emission filter: 585 ± 40 nm) or on an Attune NxT flow cytometer (excitation laser: 561 nm, emission filter: 585 ± 16 nm) at RT. At least 10,000 cells were analyzed for each sample. FCS experiments were performed as described above. Flow cytometry data was analyzed using FlowJo software (version 7.6.1, FlowJo, LLC).

##### HOPS Activity Assay Using Alexa 488 Dextran and Magic Red

Magic Red is a fluorogenic substrate for the lysosomal hydrolase cathepsin B (18). It becomes fluorescent in the presence of active cathepsin B—an enzyme exclusively localized to mature lysosomes. For the first experiment in Figures S9A–C, Saos-2 cells were transfected with siRNAs targeting VPS39 or VPS41 72 h before the assay was performed. For the second experiment described in Figures S9D–F, Saos-2 cells were plated in tissue culture-treated 6-well plates (100,000 cells/well) in complete McCoy’s 5A medium without phenol red (2 mL/well, 15% FBS, 1 mM sodium pyruvate, without P/S) 18 h before the assay was performed. After 18 or 72 h, respectively, the medium was removed and cells were washed twice with DPBS. Cells were incubated with 0.5 mg/mL dextran Alexa Fluor 488 10,000 MW, anionic, fixable (Life Technologies) in clear McCoy’s 5A medium without phenol red (1 mL/well, without FBS, without sodium pyruvate, without P/S) for 2 h at 37 °C, 5% CO_2_ followed by a 1-h chase period in dextran-free McCoy’s 5A medium. While cells were incubating, an 8-well microscopy slide (Thermo Scientific, #155411) was coated with fibronectin from bovine plasma in DPBS (1:100 dilution, 200 μL/well) for 1 h at 37 °C (200 μL/well). The fibronectin solution was removed by aspiration and the dish was washed with DPBS (3x) before use. During the hour-long chase period, cells were rinsed with DPBS (3x) and nuclei were stained with Hoechst 33342 (300 nM) for 5 min. To remove exogenously bound dextran, cells were then washed with DBPS (3x) and treated with TrypLE (500 μL/well) for 10 min until all cells were lifted from the culture dish. Cells were transferred to 15-mL Falcon tubes, each well was rinsed twice with complete McCoy’s 5A without phenol red (twice, 1 mL each), and the cell suspensions, including media used for rinsing, were pooled and pelleted at 200 *g* for 3 min. The supernatant was removed and cells were re-suspended in HEPES-containing clear DMEM without phenol red (1 mL, 25 mM HEPES, without FBS, without sodium pyruvate, without P/S) to wash the pellet. Of this suspension, half (300 μL) was plated into the prepared fibronectin-coated microscopy dish. Cells were left to adhere at 37 °C, 5% CO_2_ for the remainder of the hour-long chase period at 37 °C, 5% CO_2_. Cells were then incubated in a 1:2600 dilution of Magic Red Cathepsin B substrate (ImmunoChemistry Technologies, #937) in clear DMEM without phenol red for 5 min at 37 °C, 5% CO_2_. Cell images were recorded on an LSM 880 microscope. The degree of colocalization of two channels reflecting the fraction of dextran Alexa Fluor 488 colocalizing with Magic Red was measured by Pearson correlation coefficient (19) using ImageJ software (version 1.48s, NIH).

##### Multi-Color Confocal Fluorescence Microscopy

*With siRNA knockdown:* Saos-2 cells were transfected with siRNAs targeting VPS39 or TGFBRAP, or a non-targeting siRNA (RISC-Free) 72 h (day –3) prior to the experiment. *Without siRNA knockdown:* Saos-2 cells were plated in tissue culture-treated 6-well plates (100,000 cells/well) in complete McCoy’s 5A medium without phenol red (2 mL/well, 15% FBS, 1 mM sodium pyruvate, without P/S) 48 h (day –2) prior to the experiment. On day –1, cells were transduced with CellLight early endosomes, late endosomes, or lysosomes-GFP (BacMam 2.0, Life Technologies, #C10586, #C10588, #C10596) according to the manufacturer’s protocol. On day 0 (= 18 h after transduction), the media were removed and cells were washed twice with DPBS. Cells were incubated with *N*-Rh-PE (16:0 Liss Rhod PE, Avanti Polar Lipids, #81058) in clear McCoy’s 5A without phenol red (1 mL/well, without FBS, without sodium pyruvate, without P/S) for 1 h at 4 °C. Cells were washed with DPBS three times and incubated with CPMP **2**^R^ (300 nM) or **2**^SiR^ (600 nM) in clear McCoy’s 5A media without phenol red for 30 min at 37 °C, 5% CO_2_. While cells were incubating, an 8-well microscopy slide (Thermo Scientific, #155411) was coated with fibronectin from bovine plasma in DPBS (1:100 dilution, 200 μL/well) for 1 h at 37 °C (200 μL/well). The fibronectin solution was removed by aspiration and the dish was washed with DPBS (3x) before use. After the 30-min incubation period with CPMP **2**, nuclei were stained with Hoechst 33342 (300 nM) for 5 min at 37 °C, 5% CO_2_. To remove exogenously bound CPMPs, cells were washed with DBPS (3x) and treated with TrypLE (500 μL/well) for 10 min until all cells were lifted from the culture dish. Cells were transferred to 15-mL Falcon tubes, each well from the tissue culture dish was rinsed twice with complete McCoy’s 5A media without phenol red (twice, 1 mL each), and the cell suspensions including media used for rinsing were combined and pelleted at 200 *g* for 3 min. The supernatant was removed and cells were resuspended in HEPES-containing clear DMEM without phenol red (1 mL, 25 mM HEPES, without FBS, without sodium pyruvate, without P/S) to wash the pellet. Of this suspension, half (300 μL) was plated into the prepared fibronectin-coated microscopy dish. To enlarge Rab7^+^-endosomes, Saos-2 cells were incubated with YM201636 (800 nM) in clear DMEM without phenol red for 1 h at 37 °C, 5% CO_2_. Cell images were recorded on an LSM 880 microscope (Zeiss) using the following imaging settings: Hoechst excitation: 405 nm (Diode 405-30 laser) and emission: 410–496 nm. eGFP excitation: 488 nm (Argon laser), emission: 490–526 nm. Lissamine RhoB excitation: 561 nm (DPSS 561-10 laser), emission: 570–607 nm. Silicon rhodamine excitation: 633 nm (HeNe633 laser), emission 650–747 nm. The degree of colocalisation of two channels was measured by Pearson *r* correlation coefficient (19) using ImageJ software (version 1.48s, NIH).

#### Quantification and Statistical Analysis

All the data were presented as mean ± standard error from at least three independent trials. All data fitting and statistical analysis was performed using GraphPad Prism software (version 7.0a).

**Fig. S1.**
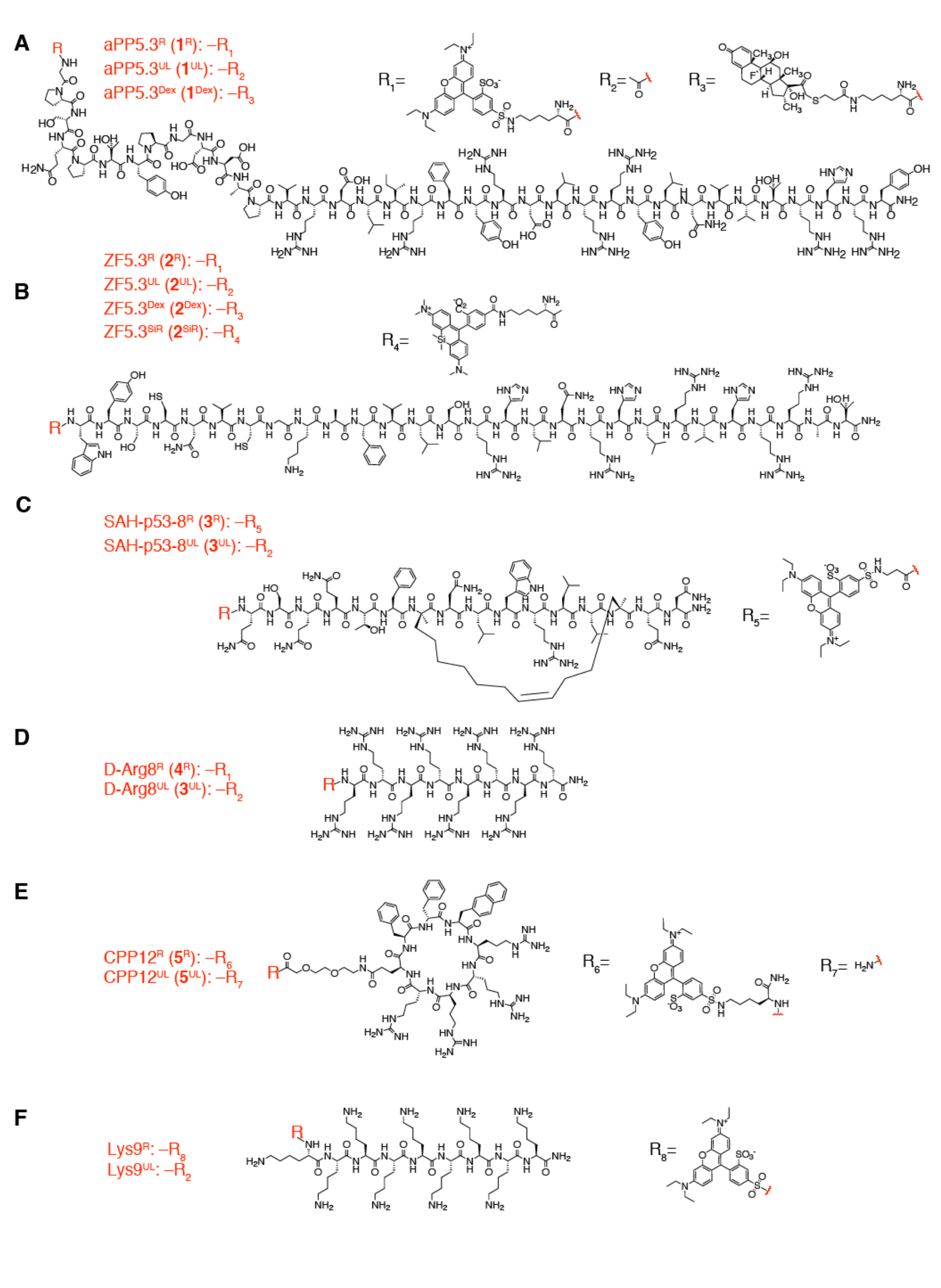
Structures of CPMP and CPP variants. Structures of unlabeled (UL), Lissamine Rhodamine B (R)-labeled, dexamethasone (Dex)-tagged, and silicon rhodamine (SiR)-labeled variants off CPMPs and CPPs studied in this work. (A) aPP5.3 (**1**) variants; (B) ZF5.3 (**2**) variants; (C) SAH-p53-8 (**3**) variants; (D) D-Arg8 (**4**) variants; (E) CPP12 (**5**) variants; (F) Lys9 variants. Note: for **1**^R^, **2**^R^, **4**^R^, and **5**^R^, the RhoB-tag was attached to a lysine side chain, while in **3**^R^, the RhoB-tag was appended to β-alanine at the N-terminus. Lys9^R^ was labeled with RhoB at the N-terminus.

**Fig. S2.**
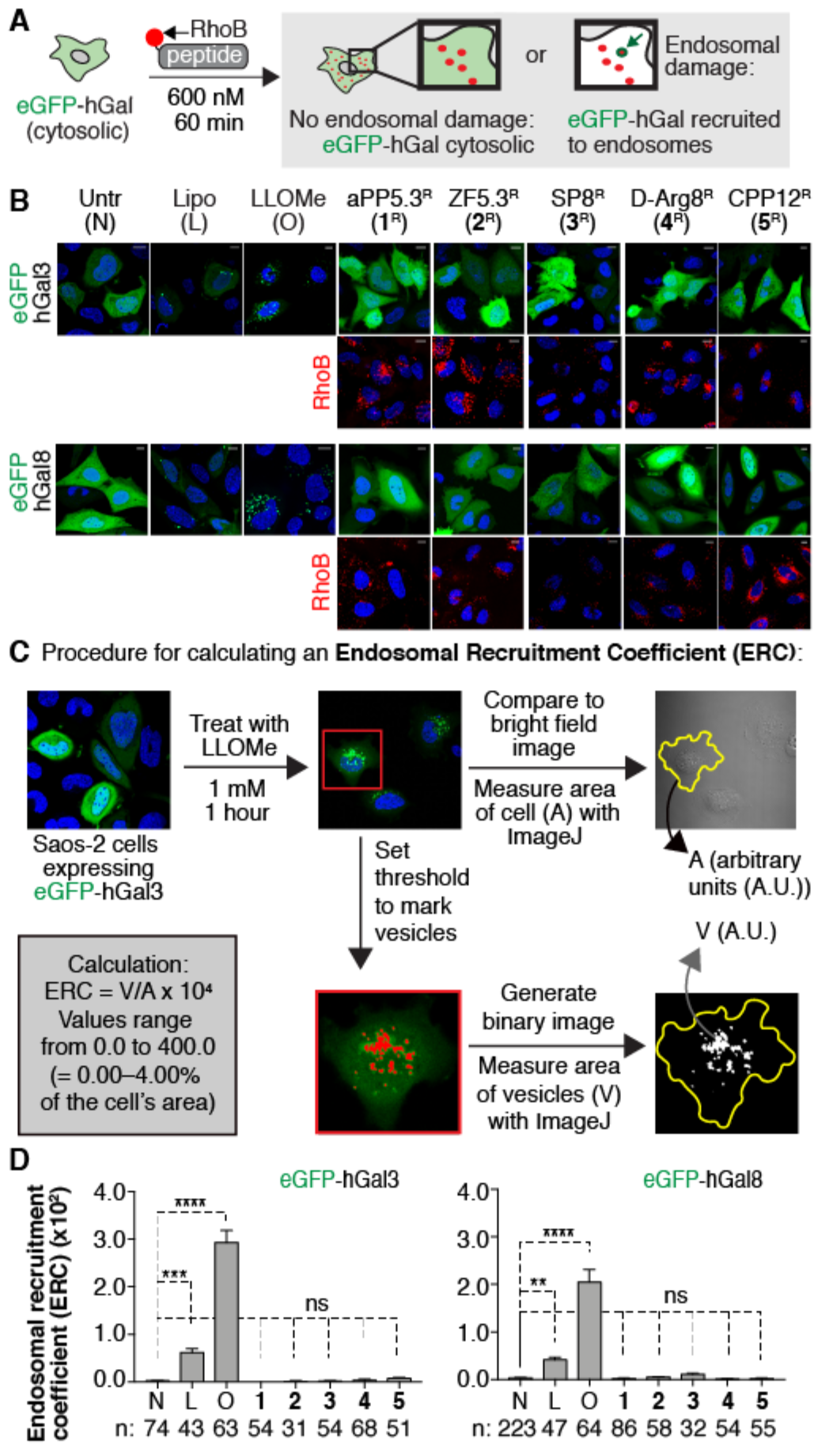
CPMPs/CPPs induce little or no galectin recruitment at 600 nM. (A) Experimental scheme. Saos-2 cells transiently expressing eGFP fusions of human galectins 3 and 8 (eGFP-hGal3 or eGFP-hGal8) were treated with 600 nM of the indicated Lissamine Rhodamine B (RhoB)-labeled CPMP or CPP (**1**^R^–**5**^R^) for 60 min. This incubation was followed by a 30 min chase in CPP/CPMP-free culture media and visualization of the cells using confocal microscopy. Cells lacking endosomal damage exhibit diffuse cytosolic fluorescence due to cytosolic eGFP-hGal3 or eGFP-hGal8. Cells with endosomal damage exhibit punctate fluorescence indicating hGal recruitment by cytosolically displayed β-galactosides on endosomal membranes. (B and D) Representative live-cell confocal fluorescence microscopy images of Saos-2 cells expressing eGFP-hGal3 or eGFP-hGal8 incubated with McCoy’s 5A media only (Untr, N), Lipofectamine RNAiMAX (16 μL/mL) (Lipo, L), LLOMe (1 mM) (O), or Rho-tagged CPMPs/CPPs **1**^R^–**5**^R^ at 600 nM. Nuclei were stained with Hoechst 33342. Top rows: green channel (detecting eGFP-hGal). Bottom rows: Rho channel (detecting RhoB). (C) Procedure for calculating an endosomal recruitment coefficient (ERC). In the example shown, Saos-2 cells expressing eGFP-hGal3 were treated with LLOMe (1 mM) for 1 hour to induce endolysosomal damage. Before LLOMe treatment, cells displayed diffuse cytosolic eGFP-hGal3 fluorescence. After LLOMe treatment, the eGFP-hGal3 fluorescence became predominantly punctate. To calculate the ERC, we measured the area (A) of each cell in the bright field channel using ImageJ. Second, we manually set a threshold to mark vesicular structures and generated a binary image thereof. Third, we measured the area of the punctate vesicles (V). Lastly, the ERC was calculated as the ratio of V over A multiplied by a factor of 10^4^. We routinely obtained ERC values around 0.0 in untreated cells and 400.0 in LLOMe-treated cells, which corresponds to 0% and 4% vesicular staining compared to the total cell area. (D) ERCs were calculated for n cells and error bars represent the standard error of the mean. ****p < 0.0001, ***p < 0.001, **p < 0.01, *p < 0.05 and not significant (ns) for p > 0.05 from one-way ANOVA with Dunnett post-test. Scale bars = 10 μm.

**Fig. S3.**
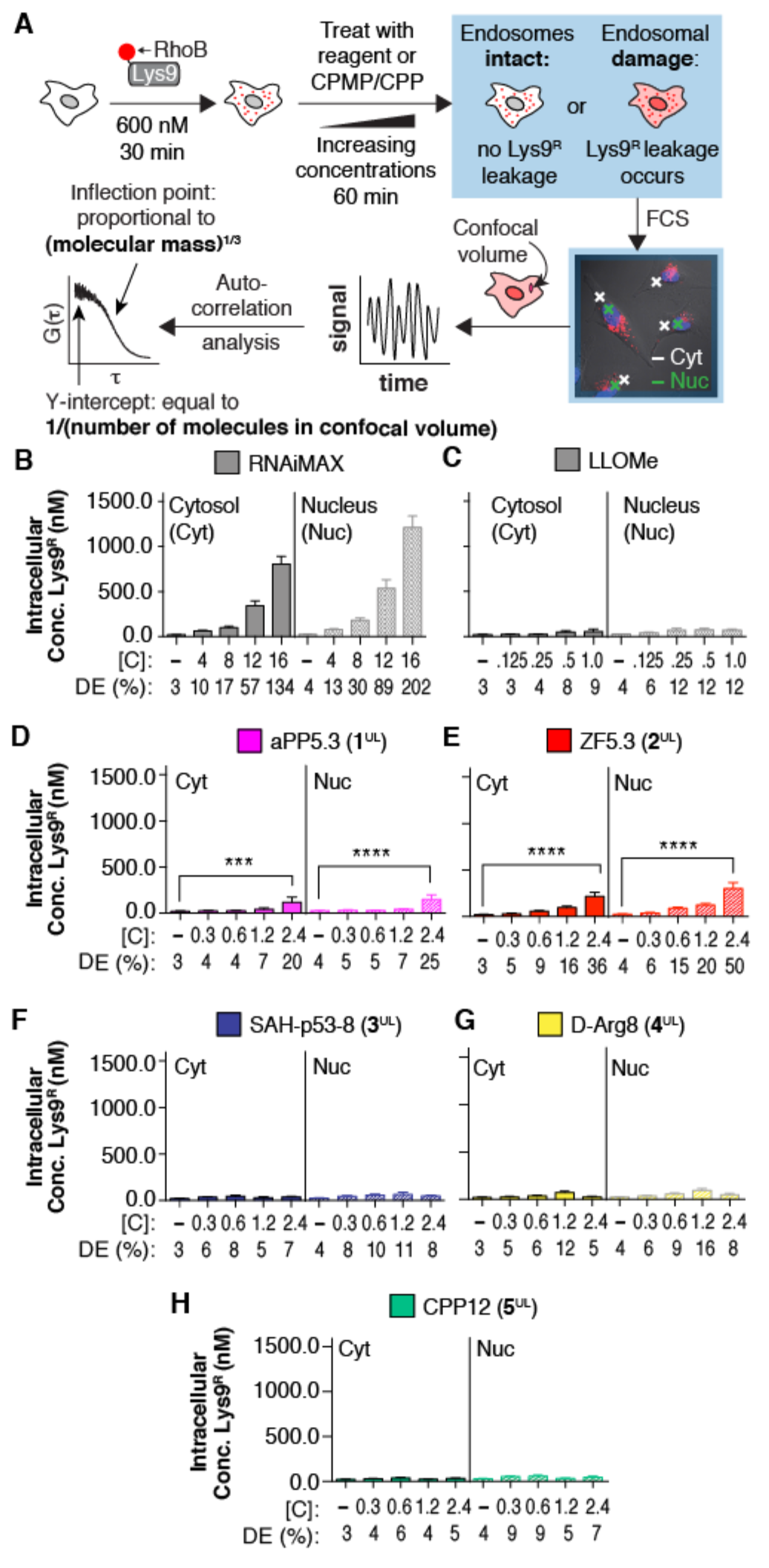
CPMPs/CPPs do not induce endosomal leakage at concentrations below 2 μM. (A) Experimental scheme. Saos-2 cells were treated with Lys9^R^ (600 nM) for 30 minutes, washed extensively, and treated with either Lipofectamine RNAiMAX (0, 4, 8, 12, 16 μL/mL), LLOMe (0, 0.125, 0.25, 0.5, 1.0 mM) or with CPMP or CPP **1**^UL^–**5**^UL^ (0.3, 0.6, 1.2, 2.4 μM) for 60 minutes. The concentration of Lys9^R^ in the cytosol and nucleus was determined using FCS. Cells whose endosomes remain intact are characterized by low concentrations of Lys9^R^ in the cytosol and nucleus, whereas cells whose endosomes have released some or all of their contents are characterized by higher levels of Lys9^R^ in the cytosol and nucleus. FCS data was obtained using a commercial Zeiss LSM 880 microscope. Images of cells were acquired in the x–y plane, and the laser’s crosshair (= confocal volume) was placed in the nucleus or cytosol of cells, avoiding areas with high, punctate endosomal signal. Fluorescent fluctuations over time were recorded on a GaAsP detector and traces were converted to autocorrelation curves using a custom MATLAB script. Individual autocorrelation traces were averaged prior to fitting to a 3D diffusion model containing parameters for anomalous diffusion and background autocorrelation as described in LaRochelle et al., 2015. The fitted data points were assessed individually and filtered for data quality by evaluating diffusion time (τ), signal intensity, and the anomalous diffusion coefficient (α). The inflection point of the autocorrelation fit is proportional to the cubic root of the molecular mass. The y-intercept equals the inverse of the number of molecules in the confocal volume. For more details, see methods section. (B–G) Intracellular Lys9^R^ concentration in the cytosol (Cyt) or nucleus (Nuc) of Lys9^R^-pretreated Saos-2 cells incubated with Lipofectamine RNAiMAX ([C] = 0, 4, 8, 12, 16 μL/mL), LLOMe ([C] = 0, 0.125, 0.25, 0.5, 1.0 mM) or CPMPs/CPPs **1**^UL^–**5**^UL^ ([C] = 0.3, 0.6, 1.2, 2.4 μM). Intracellular Lys9^R^ concentrations were calculated for n > 10 cells. The delivery efficiency (DE) is the ratio of the measured intracellular (Cyt or Nuc) Lys9^R^ concentration to the Lys9^R^ incubation concentration (600 nM) multiplied by 100%. Error bars represent the standard error of the mean. ****p < 0.0001, ***p < 0.001, and not significant (ns) for p > 0.05 from one-way ANOVA with Dunnett post-test.

**Fig. S4.**
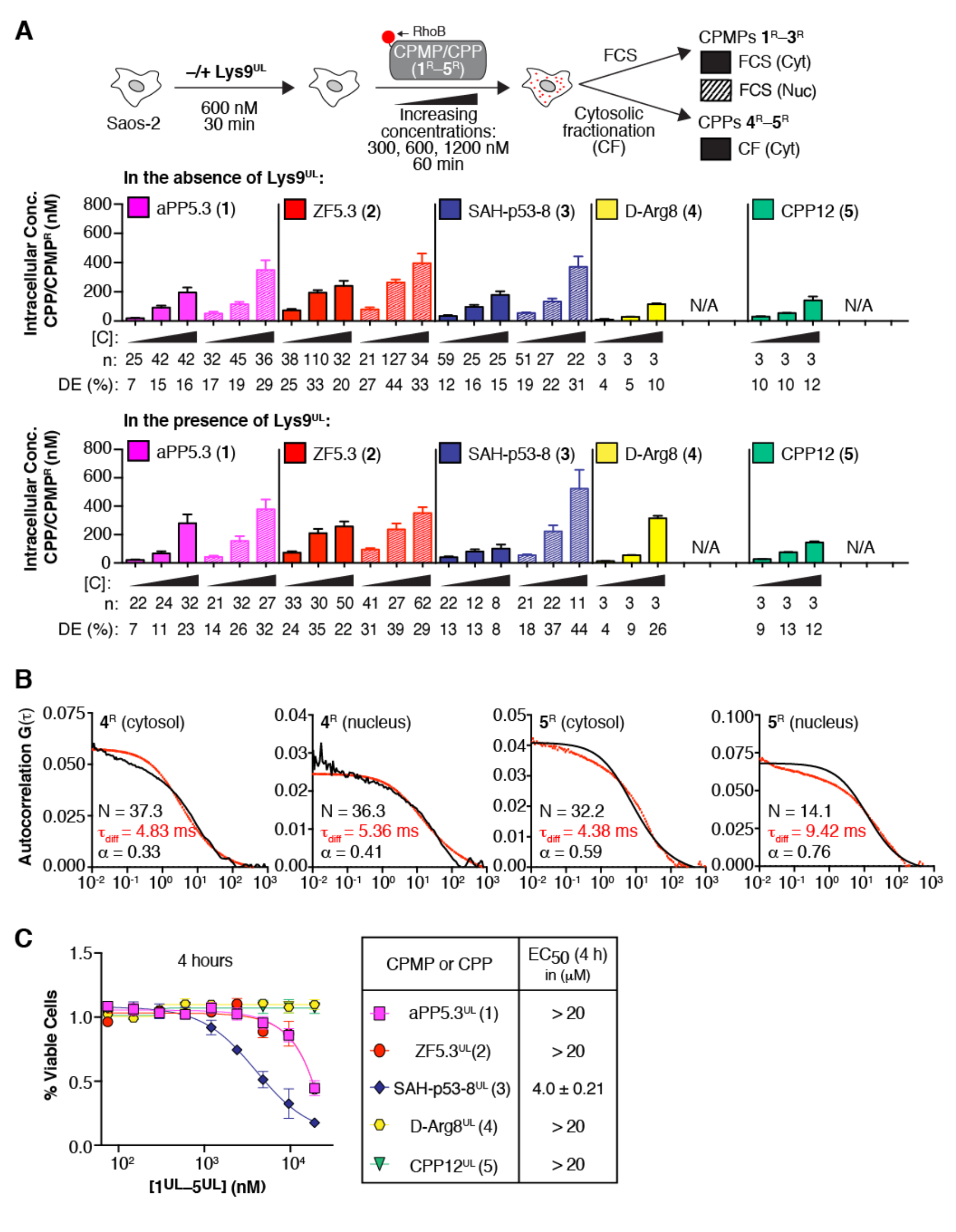
Control experiments to support and accompany Figure S3. (A) To investigate the effects of Lys9 on the intracellular delivery of peptides **1**–**5**, we incubated Saos-2 cells that had been pre-treated with or without unlabeled Lys9 (Lys9^UL^) (600 nM) with Lissamine Rhodamine B (RhoB)-tagged peptides **1**^R^–**5**^R^ (300, 600, and 1200 nM), and quantified intracellular delivery of **1**^R^–**3**^R^ *via* FCS and of **4**^R^ and **5**^R^ *via* fractionation. We were unable to perform FCS experiments at CPMP/CPP concentrations above 1200 nM because the concentration of fluorescent molecules in the confocal volume was too high—especially for CPMP **2**^R^ and hydrocarbon-stapled peptide **3**^R^**—**to obtain satisfactory fits to their corresponding autocorrelation curves. For FCS, concentrations were calculated for n cells in the cytosol (Cyt) and nucleus (Nuc) and error bars represent the standard error of the mean. For fractionation, concentrations were calculated for n biological replicates of 1.5 × 10^6^ homogenized cells and normalized to an equal population of cells treated with **2**^R^ at the same concentration. The cytosolic concentration of **2**^R^ is known because it was measured using FCS. For fractionation, nuclear concentrations are not available (N/A). The delivery efficiency (DE) is the ratio of the measured intracellular (Cyt or Nuc) Lys9^R^ concentration to the Lys9^R^ incubation concentration (600 nM) multiplied by 100%. (B) Representative traces of unusually long diffusion times observed for CPP12^R^ (**4**^R^) and D-Arg8^R^ (**5**^R^) in live cells. Saos-2 cells were treated with **4**^R^ or **5**^R^ (600 nM) for 30 min at 37 °C, 5% CO_2_. Then, cells were lifted with TripLE Express, washed, pelleted, and re-plated in a glass microscopy slide for FCS measurements. N: number of molecules detected in the confocal volume. *τ_diff_*: diffusion time. α: anomalous diffusion coefficient. (C) Effect of CPPs/CPMPs **1**^R^–**5**^R^ on cell proliferation. Plot of % viable cells remaining after 4 h treatment with [molecule] shown. Viability was assessed by monitoring oxyluciferin production by Ultra-Glow luciferase, a reaction that requires ATP. Error bars show standard error of the mean. EC_50_ values in μM.

**Fig. S5.**
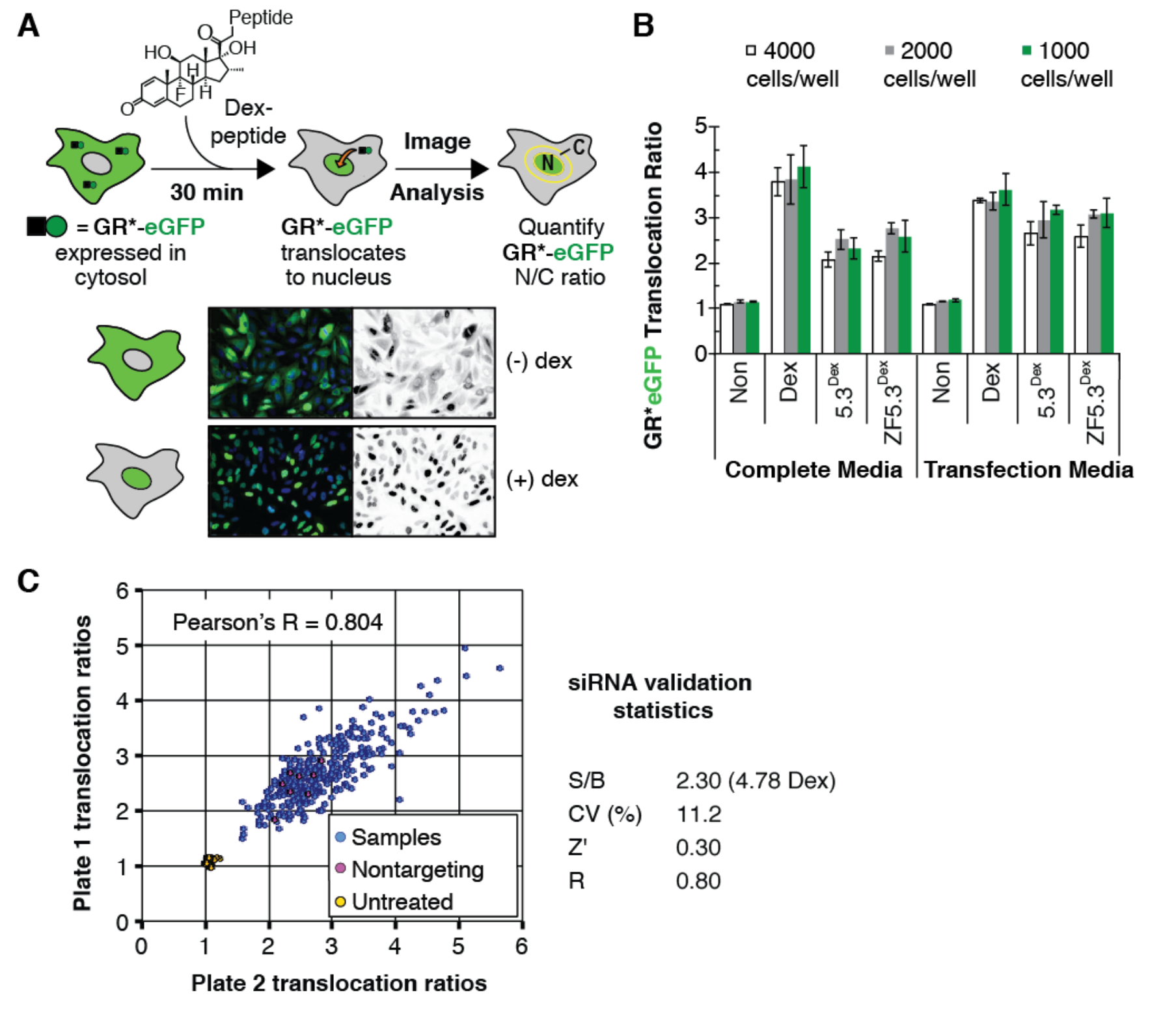
Supplementary figures to support and accompany Figure 2. (A) Overview of the glucocorticoid-induced eGFP translocation (GIGT) assay. Saos-2 cells stably expressing GR(C638G)-eGFP (= GR*-eGFP) were treated with dexamethasone (Dex) or a Dex-labeled peptide to induce the nuclear translocation of GR*-eGFP. The extent of nuclear translocation was quantified by a translocation ratio (TR), defined as the ratio of the nuclear to cytoplasmic (N/C) signal due to GR*-eGFP as determined using fluorescence microscopy and high content image analysis software (CellProfiler35 and Acapella). (B) Optimization of seeding density and RNAi growth medium in Saos-2(GIGT) cells. Saos-2(GIGT) cells were plated at various seeding densities in either complete growth medium (McCoy’s 5A, with pen/strep, 15% FBS) or transfection medium (complete growth medium/Opti-MEM/RNAi duplex buffer, 3:1:1). After 72 h, cells were treated with DMEM containing either 1 *μ*M Dex or a CPMP-Dex conjugate (**1**^Dex^ or **2**^Dex^) for 30 min, after which cells were fixed with paraformaldehyde, stained with Hoechst 33342, and imaged on an Opera High Content Screening System. Control cells (Non) were treated with media only. The RNAi screen was performed using 2,500 cells per well plated in transfection medium, as these conditions resulted in a high number of cells per well without overcrowding and resulted in the highest TRs for Dex alone. (C) Reproducibility of a RNAi pilot screen of 320 randomly chosen, duplicate siRNAs from the Dharmacon Human siGENOME siRNA Library (SMARTpool). Across this siRNA test panel, the GIGT system yielded a mean signal-to-background ratio of 2.3, coefficient of variation (CV) of 11.2%, Z-factor (Z′) of 0.3, and Pearson’s correlation coefficient (R) of 0.8.

**Fig. S6.**
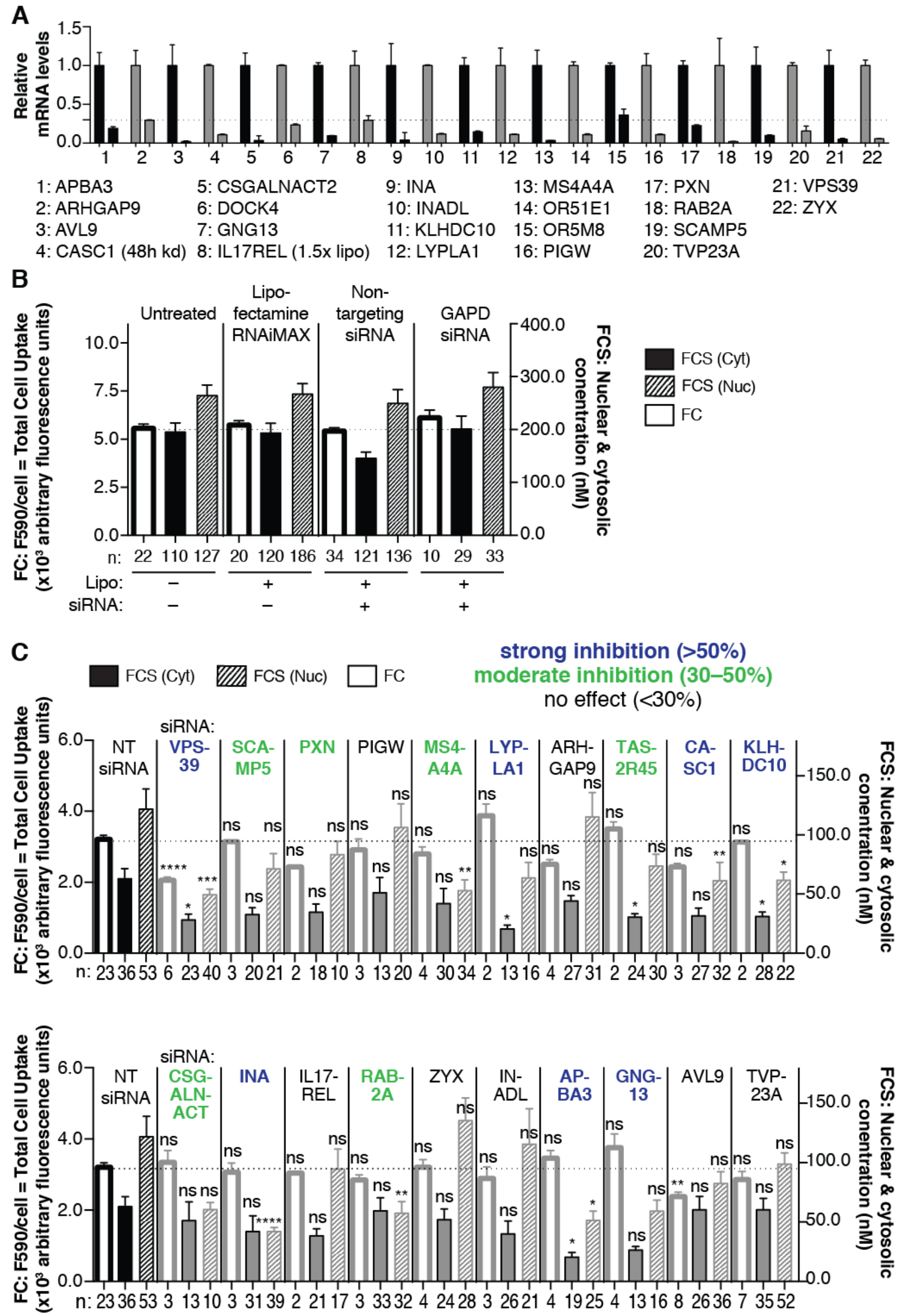
Supplementary figures related to to support and accompany Figure 3. (A) RT-qPCR to confirm gene knockdown: Saos-2 cells were transfected with Lipofectamine RNAiMAX and siGENOME SMARTpool siRNAs (Dharmacon). Total RNA was extracted 72 h (unless noted otherwise) after transfection. RNA was reverse-transcribed with gene-specific primers according to the manufacturer’s protocol. For RT-qPCR, cDNA was amplified using SsoFast EvaGreen Supermix assay. Each biological replicate was run in triplicate, and assayed for both the siRNA-targeted gene and GAPDH (endogenous reference gene). (B) Negative transfection controls for FC and FCS. Saos-2 cells were transfected with siGENOME SMARTpool siRNAs (Dharmacon) for 72 hours according to the manufacturer’s protocol. Cells were treated with CPMP **2**^R^ (600 nM) for 30 minutes and exogenously bound peptide was removed with TrypLE Express. Whole-cell fluorescence intensities were measured by FC and cytosolic and nuclear concentrations were measured by FCS in untreated (without Lipofectamine RNAiMAX, without siRNA) cells, in cells treated with Lipofectamine RNAiMAX, in cells treated with Lipofectamine RNAiMAX and a non-targeting (RISC-Free) siRNA, and in cells treated with Lipofectamine RNAiMAX and an siRNA against the *GAPD* housekeeping gene. No significant differences were detected among all controls. (C) FC and FCS data for cells transfected with the same series of 20 genes and treated with hydrocarbon-stapled peptide **3**^R^ (600 nM) and compared to non-targeting (NT, RISC-Free) siRNA-transfected cells. Left axis: FC = total cell uptake, fluorescence intensity per cell at 590 nm. Each data point (n) represents one biological replicate. For each FC replicate, the median fluorescence intensity at 590 nm was measured for at least 10,000 Saos-2 cells (gated for live cells). Right axis: FCS: cytosolic and nuclear concentration (nM). Each data point (n) represents a 50-second FCS measurement recorded in a single cell. Error bars represent the standard error of the mean. ****p < 0.0001, ***p < 0.001, **p < 0.01, *p < 0.05, and not significant (ns) for p > 0.05 from one-way ANOVA with Dunnett post-test.

**Fig. S7.**
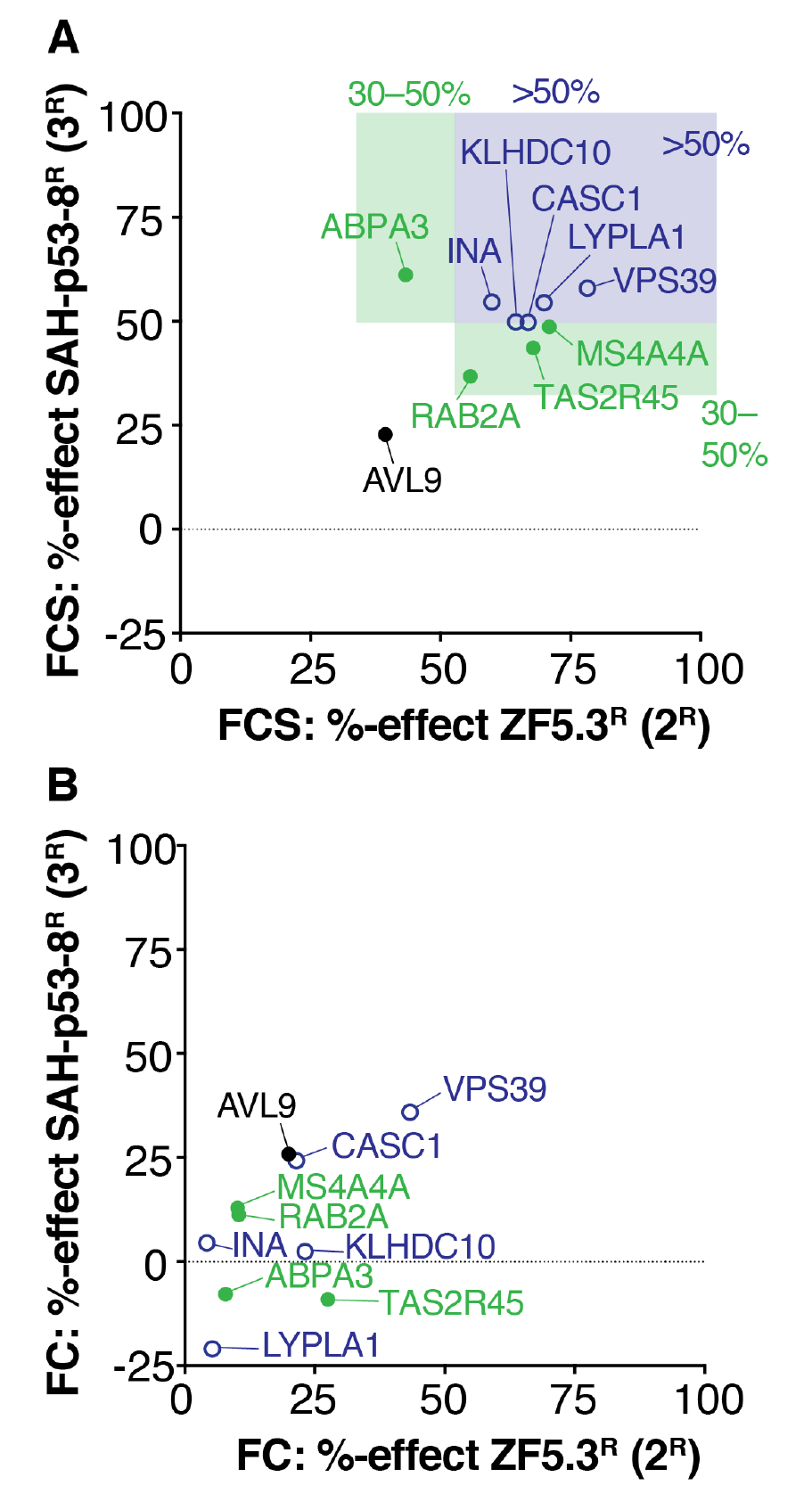
Comparison of the effects of gene-specific knockdown on uptake and cytosolic trafficking of CPMP2 2 and hydrocarbon-stapled peptide 3. (A) Comparison of the effects of gene-specific knockdowns on the cytosolic trafficking of CPMP **2**^R^ and hydrocarbon stapled peptide **3**^R^ as determined by FCS. Graph includes only those genes with significant (by one-way ANOVA with Dunnett post-test) FCS effects for both **2**^R^ and **3**^R^. The %-effect for each siRNA knockdown was calculated by dividing the average intracellular concentration (C_nucleus_ + C_Cytosol_ divided by two) measured by FCS in gene-specific siRNA-transfected cells by the average intracellular concentration measured in cells transfected with a non-targeting siRNA (RISC-Free), multiplied by 100%. (B) Comparison of the effects of gene-specific knockdowns (same genes as in (A)) on the cytosolic trafficking of CPMP **2**^R^ and hydrocarbon stapled peptide **3**^R^ as determined by flow cytometry (FC). The %-effect for each siRNA knockdown was calculated by dividing the average overall uptake fluorescence intensity measured by FC in gene-specific siRNA-transfected cells by the average overall uptake fluorescence intensity measured in cells transfected with a non-targeting siRNA (RISC-Free), multiplied by 100%.

**Fig. S8.**
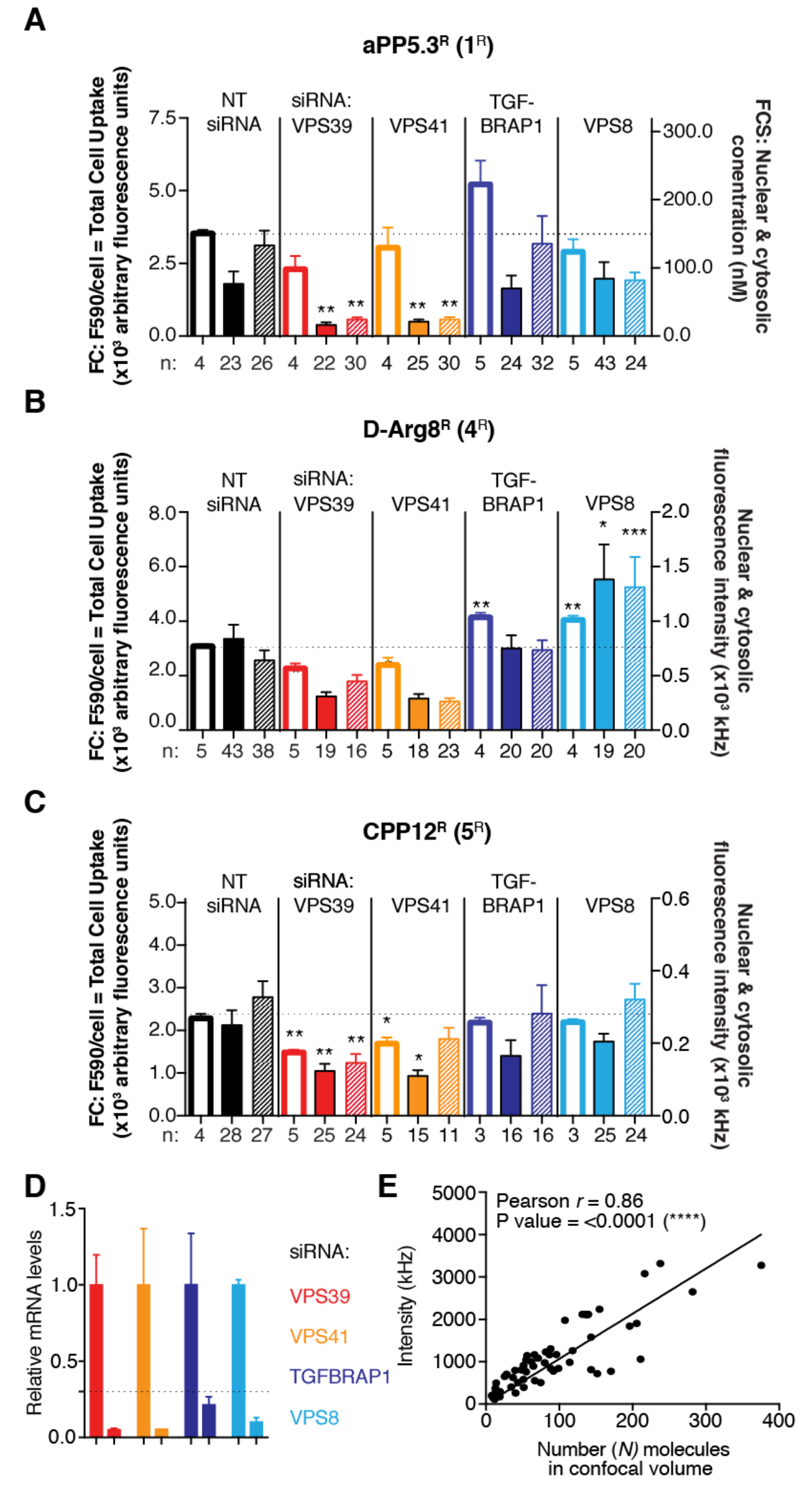
Trafficking of 1^R^, 4^R^, and 5^R^ also require the human HOPS complex. (A) The effect of VPS39 and VPS41 knockdown (HOPS-specific subunits) compared to TGFBRAP1 and VPS8 knockdown (CORVET-specific subunits) on overall uptake (FC) and cytosolic access (FCS) of aPP5.3^R^ (**1**^R^). Negative control: cells transected with non-targeting siRNA (RISC-Free siRNA). Left axis: FC = total cell uptake, fluorescence intensity per cell at 590 nm. Each data point (n) represents one biological replicate. For each FC replicate, the median fluorescence intensity at 590 nm was measured for at least 10,000 Saos-2 cells (gated for live cells). Right axis: FCS: cytosolic and nuclear concentration (nM). Each data point (n) denotes a 50-second FCS measurement recorded in a single cell. Error bars represent the standard error of the mean. (B and C) The effect of VPS39 and VPS41 knockdown (HOPS-specific subunits) compared to TGFBRAP1 and VPS8 knockdown (CORVET-specific subunits) on overall uptake (FC) and cytosolic access (FCS intensity without autocorrelation) of D-Arg8^R^ (**4**^R^) and CPP12^R^ (**5**^R^). Negative control: cells transected with non-targeting siRNA (RISC-Free siRNA). Left axis: FC = total cell uptake, fluorescence intensity per cell at 590 nm. Each data point (n) represents one biological replicate. For each FC replicate, the median fluorescence intensity at 590 nm was measured for at least 10,000 Saos-2 cells (gated for live cells). Right axis: FCS cytosolic and nuclear fluorescence intensity. Each data point (n) represents one 25-second FCS intensity measurement (without autocorrelation) recorded in the nucleus or cytosol of a single Saos-2 cell. Error bars represent the standard error of the mean. ****p < 0.0001, ***p < 0.001, **p < 0.01, and not significant (ns) for p > 0.05 from one-way ANOVA with Dunnett post-test. (D) Saos-2 cells were transfected with Lipofectamine RNAiMAX and siGENOME SMARTpool siRNAs (Dharmacon) against VPS39, VPS41, TGFBRAP1, and VPS8. Total RNA was extracted 72 h after transfection. RNA was reverse-transcribed with gene-specific primers according to the manufacturer’s protocol. cDNA was amplified using the SsoFast EvaGreen Supermix assay. Each biological replicate was run in triplicate, assayed for the siRNA-targeted gene, and GAPDH as an endogenous reference gene. (E) The FCS intensity (in kHz) is proportional to the number (*N*) of molecules detected in the confocal volume for a laser. Each data point represents one 50-second FCS measurement recorded in the nucleus or cytosol of a single Saos-2 cell treated with **2**^R^ (600 nM) for 30 min.

**Fig. S9.**
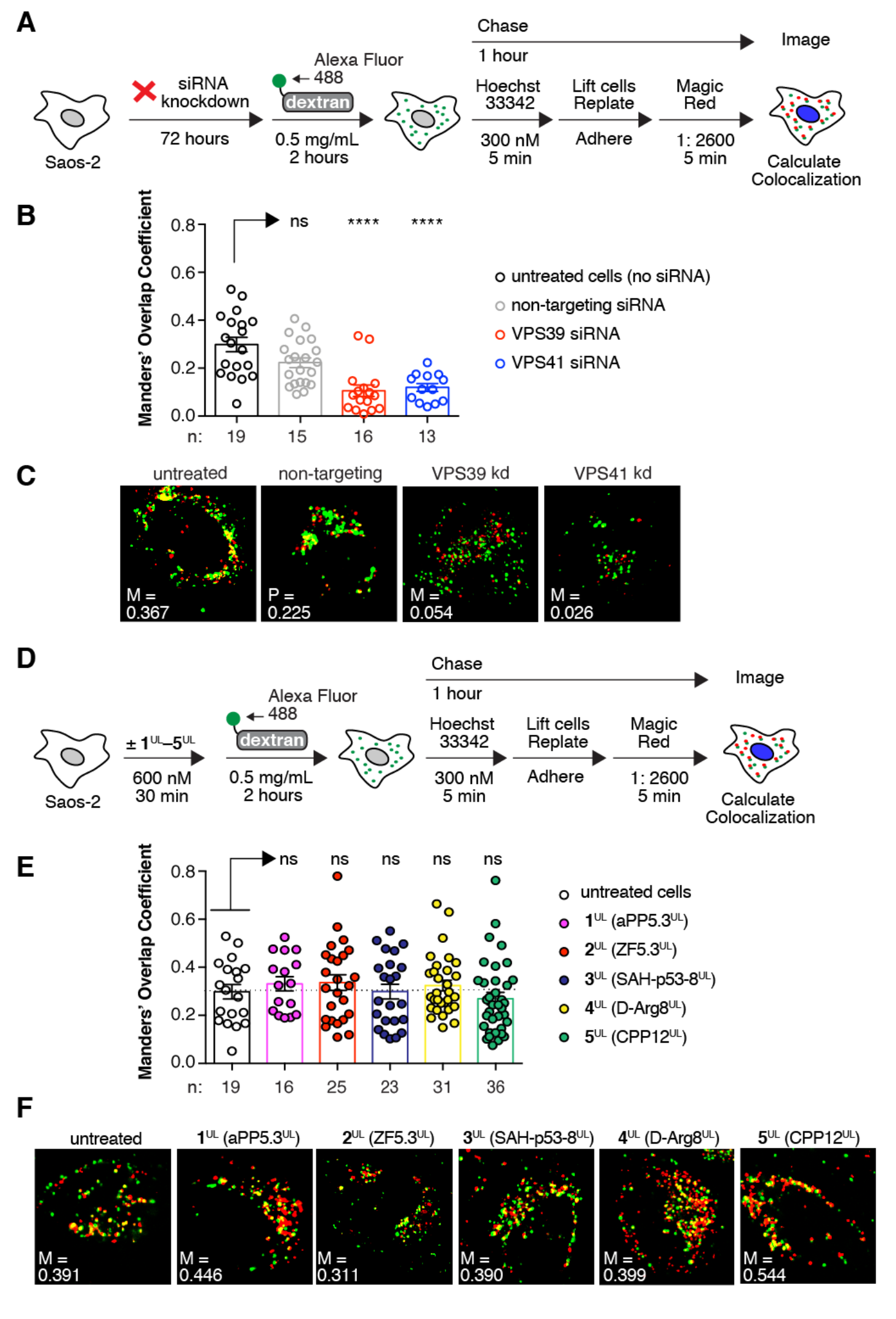
HOPS activity studies. (A) Saos-2 cells were transfected with Lipofectamine RNAiMAX and siGENOME SMARTpool siRNAs (Dharmacon) targeting VPS39 or VPS41. After 72 h, cells were incubated with Alexa Fluor 488-tagged dextran (10,000 MW, anionic, fixable) for 2 h. Cells were washed, nuclei were stained with Hoechst 33342, cells were replated, and lysosomes were stained with Magic Red for 5 min. (B) Colocalization of Alexa Fluor 488-tagged dextran with Magic Red (Manders M2) in live-cell confocal microscopy images after knockdown of HOPS proteins VPS39 and VPS41 with SMARTpool siRNA compared to a non-targeting (RISC-Free) RNA. Error bars represent the standard error of the mean. ****p < 0.0001 from one-way ANOVA with Dunnett post-test. (C) Representative live-cell confocal microscopy images of cells quantified in (B). (D) Saos-2 cells were treated with CPMPs/CPPs **1**^UL^–**5**^UL^. After 30 min, the media was replaced and cells were incubated with Alexa Fluor 488-tagged dextran for 2 h. Cells were washed, nuclei were stained with Hoechst 33342, cells were replated, and lysosomes were stained with Magic Red dye for 5 min. (E) Colocalization of Alexa Fluor 488-tagged dextran with Magic Red (Manders M2) in live-cell confocal microscopy images after treatment with CPMPs/CPPs **1**^UL^–**5**^UL^ for 30 min at 37 °C, 5% CO_2_. Error bars represent the standard error of the mean. Not significant (ns) for p > 0.05 from one-way ANOVA with Dunnett post-test. (F) Representative live-cell confocal microscopy images of cells quantified in (E).

**Fig. S10.**
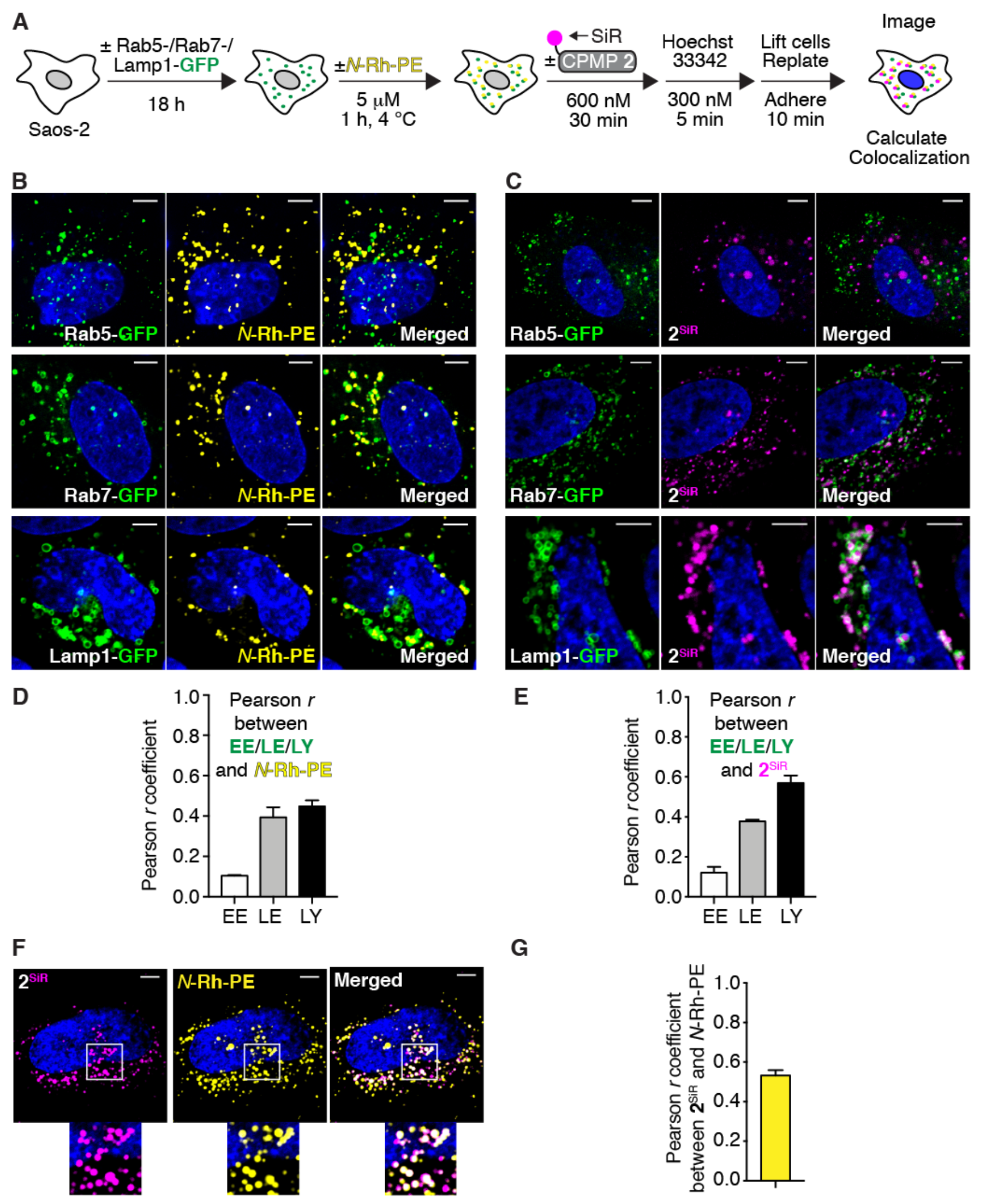
CPMP 2 localizes to the lumen of LEs and LYs. (A) Experimental scheme. Saos-2 cells were transduced with Rab5-, Rab7-, or Lamp1-GFP for 18 h using CellLight Reagents (BacMam 2.0). Cells were washed and incubated with (or without) *N*-Rh-PE for 1 h at 4 °C. Cells were washed and incubated with (or without) CPMP **2**^SiR^ for 30 min, nuclei were stained with Hoechst 33342, and cells were replated into microscopy slides. (B and C) Representative live-cell confocal microscopy images of Saos-2 cells prepared as described in (A) containing GFP markers and *N*-Rh-PE (B) or GFP markers and **2**^SiR^ (C). (D and E) Pearson correlation coefficients between GFP markers and *N*-Rh-PE (D), and GFP markers and **2**^SiR^ (E). (F) Representative live-cell confocal microscopy images of Saos-2 cells treated with CPMP **2**^SiR^ and ILV marker *N*-Rh-PE. (G) Pearson correlation coefficient between 2^SiR^ and ILV marker *N*-Rh-PE. Scale bars = 5 μm.

**Fig. S11.**
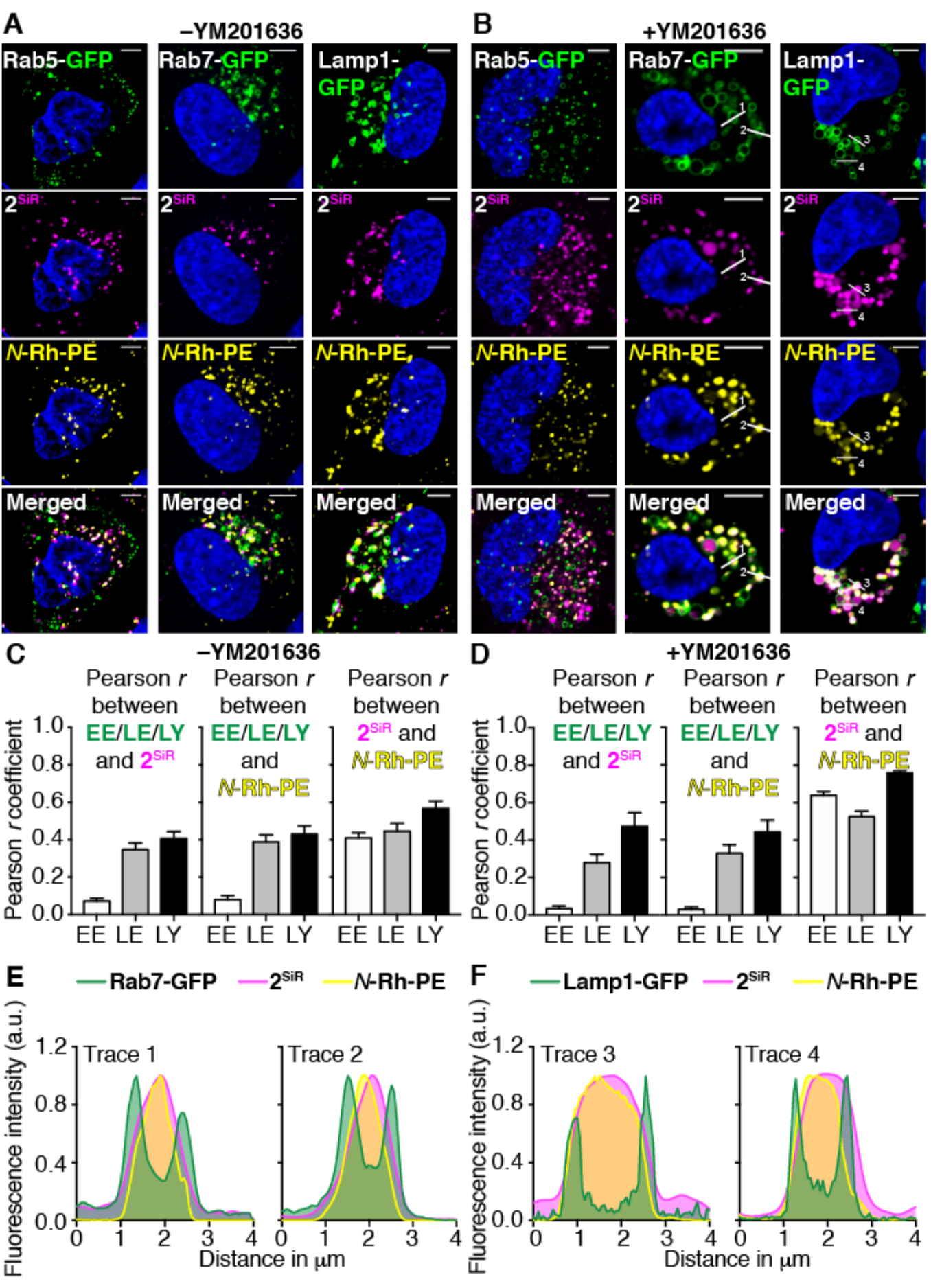
CPMP 2 colocalizes with ILV marker *N*-Rh-PE. CPMP **2** localizes to ILVs in LEs and LYs. Saos-2 cells were transduced with Rab5-, Rab7-, or Lamp1-GFP for 18 h using CellLight Reagents (BacMam 2.0). To label ILVs, cells were incubated with *N*-Rh-PE (5 μM) for 1 h at 4 °C. Cells were washed and incubated with CPMP **2**^SiR^ (600 nM) for 30 min and nuclei were stained with Hoechst 33342 (300 nM) for 5 min. Cells were trypsinized and re-plated into microscopy slides, after which they were incubated in media with or without YM201636 (800 nM) for 1 hour. (A and B) Representative live-cell confocal fluorescence microscopy images of Saos-2 cells incubated without YM201636 (A) or with YM201636 (B) present. (C and D) Pearson correlation coefficients between pairs of specified markers without YM201636 (C) or with YM201636 (D) present. (E and F) Fluorescence intensity line profiles of endosomes (#1–4, see panel C) generated in ImageJ. Scale bars = 5 μm.

**Table S1.**
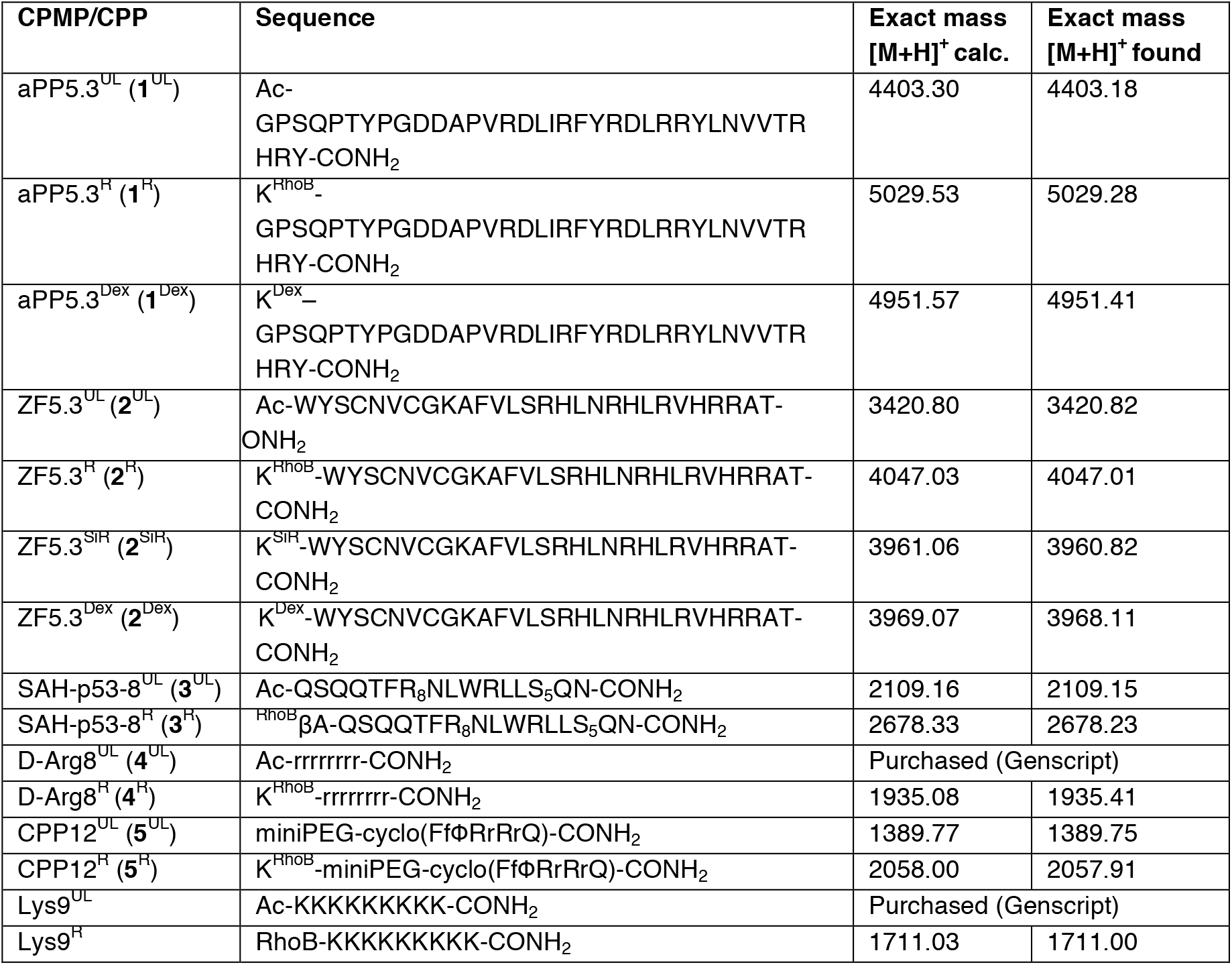
Sequences, calculated & observed exact masses of all CPMP and CPP variants.

**Table S2.** FCS diffusion parameters. Molecular weight, predicted and measured diffusion coefficients (*D*), average *in vitro* and *in cellulo* diffusion times (*τ*) in the cytosol and nucleus of RhoB-tagged CPMP and CPP variants.

| | **Molecular weight** | **Predicted $D$**<br>at 25 °C<br>($\mu\text{m}^2 \text{s}^{-1}$) | **Measured $D$**<br>at 25 °C<br>($\mu\text{m}^2 \text{s}^{-1}$) | **Average $\tau_{diff}$ *in vitro*** at<br>25 °C (ms) | **Average $\tau_{diff}$ *in cellulo*** at<br>37 °C (ms)<br>In cytosol | **Average $\tau_{diff}$ *in cellulo*** at<br>37 °C (ms)<br>In nucleus |
| --- | --- | --- | --- | --- | --- | --- |
| Alexa 594 | 758.79 | 388 | 383 ± 7 | 0.036 ±<br>0.0009 | N/A | N/A |
| aPP5.3<sup>R</sup><br>(1<sup>R</sup>) | 5031.77 | 207 | 171 ± 8 | 0.118 ±<br>0.012 | 0.262 ±<br>0.049 | 0.218 ±<br>0.017 |
| ZF5.3<sup>R</sup><br>(2<sup>R</sup>) | 4048.80 | 223 | 169 ± 4 | 0.074 ±<br>0.002 | 0.212 ±<br>0.025 | 0.227 ±<br>0.027 |
| SAH-p53-<br>8<sup>R</sup> (3<sup>R</sup>) | 2679.16 | 255 | 223 ± 6 | 0.068 ±<br>0.005 | 0.539 ±<br>0.155 | 0.326 ±<br>0.117 |
| D-Arg8<sup>R</sup><br>(4<sup>R</sup>) | 1935.37 | 285 | 202 ± 6 | 0.076 ±<br>0.006 | 5.67 ±<br>0.408 | 4.30 ±<br>0.368 |
| CPP12<sup>R</sup><br>(5<sup>R</sup>) | 2058.46 | 279 | 221 ± 2 | 0.057 ±<br>0.001 | 5.97 ±<br>0.348 | 4.83 ±<br>0.365 |
| Lys9<sup>R</sup> | 1711.26 | 296 | 206 ± 3 | 0.064 ±<br>0.001 | 0.274 ±<br>0.019 | 0.236 ±<br>0.015 |

**Table S3.**
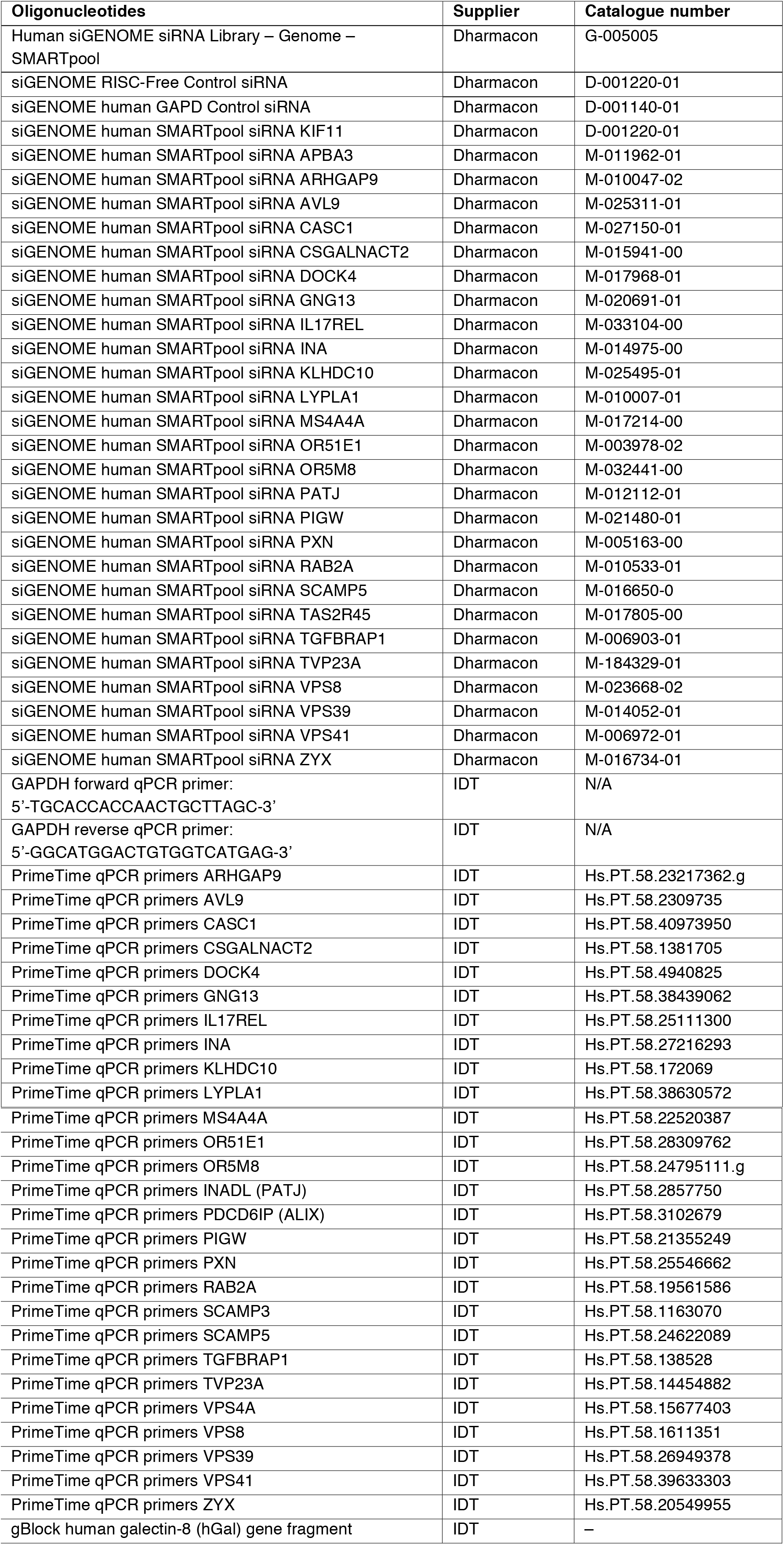
siRNAs and RT-qPCR primers.

**Movie S1. Single-color movie of ILV movement in Lamp1^+^ endosomes.** Saos-2 cells were transduced with Lamp1-GFP for 18 h using CellLight Reagents (BacMam 2.0). Cells were washed and incubated with CPMP 2^R^ (300 nM) for 30 min and nuclei were stained with Hoechst 33342 (300 nM) for 5 min. Cells were lifted with TrypLE Express and re-plated into microscopy slides. Cells were incubated in media containing YM201636 (800 nM) for 1 hour. The GFP, RhoB, and Hoechst channels corresponding to the first frame of this movie can be found in Fig. 5H.

**Movie S2. Dual-color movie of ILV movement in Lamp1^+^ endosomes.** Saos-2 cells were transduced with Lamp1-GFP for 18 h using CellLight Reagents (BacMam 2.0). Cells were washed and incubated with CPMP 2^R^ (300 nM) for 30 min and nuclei were stained with Hoechst 33342 (300 nM) for 5 min. Cells were lifted with TrypLE Express and re-plated into microscopy slides. Cells were incubated in media containing YM201636 (800 nM) for 1 hour.

**Additional Data Table S4. Primary hits (428) of the GIGT siRNA screen.** By setting an SSMD (see equation 12) threshold of ±2.0, we identified 263 genes whose knockdown enhance and 165 genes whose knockdown inhibit cytosolic access of CPMP **1**^Dex^.

**Additional Data Table S5. Prioritized hits (131) of the GIGT siRNA screen.** Of the 428 primary hits, we prioritized genes according to function. We retained genes with known roles in endocytic trafficking and genes with unknown or poorly characterized functions.

**Additional Data Table S6. Raw data of the GR*-eGFP translocation screen.** During this filtering step, we retained a gene if its knockdown did not induce a significant change in the translocation ratio (TR) for at least two of the four gene-specific siRNAs (A–D) compared to cells transfected with a non-targeting (RISC-Free) siRNA. This process eliminated 61 of 131 genes with 70 genes remaining for follow-up analysis. ****p < 0.0001, ***p < 0.001, **p < 0.01, *p < 0.05, and not significant (ns) for p > 0.05 from one-way ANOVA with Dunnett post-test.

**Additional Data Table S7. Raw data of the siRNA deconvolution screen with CPMPs 1^Dex^ and 2^Dex^.** In this final filtering step, we prioritized siRNAs that induced significant TR changes for at least two out of the four gene-specific siRNAs (A–D) in cells treated with either CPMP **1**^Dex^ or **2**^Dex^ (or both), when compared to cells transfected with a non-targeting (RISC-Free) siRNA. siRNAs inducing significant TR changes are highlighted in blue (decrease) and red (increase). Union (final hits): 28 genes passed this filter for *either* CPMP **1**^Dex^ or **2**^Dex^. Intersection: 9 genes passed this filter for *both* CPMPs **1**^Dex^ or **2**^Dex^. ****p < 0.0001, ***p < 0.001, **p < 0.01, *p < 0.05, and not significant (ns) for p > 0.05 from one-way ANOVA with Dunnett post-test.

## References

1. Lagasse HA, et al. (2017) Recent advances in (therapeutic protein) drug development. F1000Res 6:113.

2. Yao S, Zhu YW, & Chen LP (2013) Advances in targeting cell surface signalling molecules for immune modulation. Nat Rev Drug Discov 12:130–146.

3. Serna N, et al. (2018) Protein-Based Therapeutic Killing for Cancer Therapies. Trends Biotechnol 36:318–335.

4. Lamb YN & Syed YY (2018) LY2963016 Insulin Glargine: A Review in Type 1 and 2 Diabetes. Biodrugs 32:91–98.

5. Li P, Zheng Y, & Chen X (2017) Drugs for Autoimmune Inflammatory Diseases: From Small Molecule Compounds to Anti-TNF Biologics. Front Pharmacol 8.

6. Malviya G, et al. (2013) Biological Therapies for Rheumatoid Arthritis: Progress to Date. Biodrugs 27:329–345.

7. Strand V, Kimberly R, & Isaacs JD (2007) Biologic therapies in rheumatology: lessons learned, future directions. Nat Rev Drug Discov 6:75–92.

8. Agarraberes FA & Dice JF (2001) Protein translocation across membranes. Biochim Biophys Acta Bio 1513:1–24.

9. Schatz G & Dobberstein B (1996) Common principles of protein translocation across membranes. Science 271:1519–1526.

10. Green M & Loewenstein PM (1988) Autonomous functional domains of chemically synthesized human immunodeficiency virus tat trans-activator protein. Cell 55:1179–1188.

11. Frankel AD & Pabo CO (1988) Cellular uptake of the tat protein from human immunodeficiency virus. Cell 55:1189–1193.

12. Derossi D, Joliot AH, Chassaing G, & Prochiantz A (1994) The third helix of the Antennapedia homeodomain translocates through biological membranes. J Biol Chem 269:10444–10450.

13. Guidotti G, Brambilla L, & Rossi D (2017) Cell-Penetrating Peptides: From Basic Research to Clinics. Trends Pharm Sci 38:406–424.

14. Milletti F (2012) Cell-penetrating peptides: classes, origin, and current landscape. Drug Discov Today 17:850–860.

15. El-Sayed A, Futaki S, & Harashima H (2009) Delivery of Macromolecules Using Arginine-Rich Cell-Penetrating Peptides: Ways to Overcome Endosomal Entrapment. AAPS J 11:13–22.

16. Zondlo NJ & Schepartz A (1999) Highly specific DNA recognition by a designed miniature protein. J Am Chem Soc 121:6938–6939.

17. Chin JW & Schepartz A (2001) Concerted evolution of structure and function in a miniature protein. J Am Chem Soc 123:2929–2930.

18. Appelbaum JS, et al. (2012) Arginine topology controls escape of minimally cationic proteins from early endosomes to the cytoplasm. Chem Biol 19:819–830.

19. LaRochelle JR, Cobb GB, Steinauer A, Rhoades E, & Schepartz A (2015) Fluorescence correlation spectroscopy reveals highly efficient cytosolic delivery of certain penta-arg proteins and stapled peptides. J Am Chem Soc 137:2536–2541.

20. Holub JM, Larochelle JR, Appelbaum JS, & Schepartz A (2013) Improved assays for determining the cytosolic access of peptides, proteins, and their mimetics. Biochemistry 52:9036–9046.

21. Vives E, Brodin P, & Lebleu B (1997) A truncated HIV-1 Tat protein basic domain rapidly translocates through the plasma membrane and accumulates in the cell nucleus. Journal of Biological Chemistry 272:16010–16017.

22. Bernal F, et al. (2010) A stapled p53 helix overcomes HDMX-mediated suppression of p53. Cancer Cell 18:411–422.

23. Wissner RF, Steinauer A, Knox S, Thompson A, & Schepartz A (2018) Fluorescence correlation spectroscopy reveals efficient cytosolic delivery of protein cargo by cell-permeant miniature proteins. Submitted.

24. Balderhaar HJ & Ungermann C (2013) CORVET and HOPS tethering complexes - coordinators of endosome and lysosome fusion. J Cell Sci 126:1307–1316.

25. Maiolo JR, 3rd, Ottinger EA, & Ferrer M (2004) Specific redistribution of cell-penetrating peptides from endosomes to the cytoplasm and nucleus upon laser illumination. J Am Chem Soc 126:15376–15377.

26. Qian Z, et al. (2016) Discovery and Mechanism of Highly Efficient Cyclic Cell-Penetrating Peptides. Biochemistry 55:2601–2612.

27. Wender PA, et al. (2000) The design, synthesis, and evaluation of molecules that enable or enhance cellular uptake: Peptoid molecular transporters. Proc Natl Acad Sci USA 97:13003–13008.

28. Purkayastha N, et al. (2013) Enantiomeric and Diastereoisomeric (Mixed) L/D-Octaarginine Derivatives - A Simple Way of Modulating the Properties of Cell-Penetrating Peptides. Chem Biodiv 10:1165–1184.

29. Maier O, Marvin SA, Wodrich H, Campbell EM, & Wiethoff CM (2012) Spatiotemporal dynamics of adenovirus membrane rupture and endosomal escape. J Virol 86:10821–10828.

30. Montespan C, et al. (2017) Multi-layered control of Galectin-8 mediated autophagy during adenovirus cell entry through a conserved PPxY motif in the viral capsid. PLOS Path 13:e1006217.

31. Maejima I, et al. (2013) Autophagy sequesters damaged lysosomes to control lysosomal biogenesis and kidney injury. EMBO J 32:2336–2347.

32. Meunier E, et al. (2014) Caspase-11 activation requires lysis of pathogen-containing vacuoles by IFN-induced GTPases. Nature 509:366–+.

33. Thurston TL, Wandel MP, von Muhlinen N, Foeglein A, & Randow F (2012) Galectin 8 targets damaged vesicles for autophagy to defend cells against bacterial invasion. Nature 482:414–418.

34. Feeley EM, et al. (2017) Galectin-3 directs antimicrobial guanylate binding proteins to vacuoles furnished with bacterial secretion systems. Proc Natl Acad Sci USA 114:E1698–E1706.

35. Kilchrist KV, Evans BC, Brophy CM, & Duvall CL (2016) Mechanism of Enhanced Cellular Uptake and Cytosolic Retention of MK2 Inhibitory Peptide Nano-polyplexes. Cell Mol Bioeng 9:368–381.

36. Paz I, et al. (2010) Galectin-3, a marker for vacuole lysis by invasive pathogens. Cell Microbiol 12:530–544.

37. Houzelstein D, et al. (2004) Phylogenetic analysis of the vertebrate galectin family. Mol Biol Evol 21:1177–1187.

38. Rabinovich GA & Toscano MA (2009) Turning ‘sweet’ on immunity: galectin-glycan interactions in immune tolerance and inflammation. Nat Rev Immunol 9:338–352.

39. Aits S, et al. (2015) Sensitive detection of lysosomal membrane permeabilization by lysosomal galectin puncta assay. Autophagy 11:1408–1424.

40. Wittrup A, et al. (2015) Visualizing lipid-formulated siRNA release from endosomes and target gene knockdown. Nat Biotechnol 33:870–876.

41. Pagliero RJ, et al. (2016) Discovery of Small Molecules That Induce Lysosomal Cell Death in Cancer Cell Lines Using an Image-Based Screening Platform. Assay Drug Dev Tech 14:489–510.

42. D’Astolfo DS, et al. (2015) Efficient intracellular delivery of native proteins. Cell 161:674–690.

43. Zhao M, et al. (2008) Lipofectamine RNAiMAX: An efficient siRNA transfection reagent in human embryonic stem cells. Mol Biotechnol 40:19–26.

44. Schneider CA, Rasband WS, & Eliceiri KW (2012) NIH Image to ImageJ: 25 years of image analysis. Nat Meth 9:671–675.

45. Dumic J, Dabelic S, & Flogel M (2006) Galectin-3: an open-ended story. Biochim Biophys Acta 1760:616–635.

46. Srinivasan D, et al. (2011) Conjugation to the cell-penetrating peptide TAT potentiates the photodynamic effect of carboxytetramethylrhodamine. PLOS One 6:e17732.

47. Erazo-Oliveras A, et al. (2014) Protein delivery into live cells by incubation with an endosomolytic agent. Nat Methods 11:861–867.

48. Thiele DL & Lipsky PE (1990) Mechanism of L-Leucyl-L-Leucine Methyl Ester-Mediated Killing of Cytotoxic Lymphocytes - Dependence on a Lysosomal Thiol Protease, Dipeptidyl Peptidase-I, That Is Enriched in These Cells. Proc Natl Acad Sci USA 87:83–87.

49. Uchimoto T, et al. (1999) Mechanism of apoptosis induced by a lysosomotropic agent, L-Leucyl-L-leucine methyl ester. Apoptosis 4:357–362.

50. Erazo-Oliveras A, et al. (2016) The Late Endosome and Its Lipid BMP Act as Gateways for Efficient Cytosolic Access of the Delivery Agent dfTAT and Its Macromolecular Cargos. Cell Chem Biol 23:598–607.

51. Kuhn T, et al. (2011) Protein Diffusion in Mammalian Cell Cytoplasm. PLOS One 6.

52. Li M, et al. (2015) Discovery and characterization of a peptide that enhances endosomal escape of delivered proteins in vitro and in vivo. J Am Chem Soc 137:14084–14093.

53. Chakraborti PK, Garabedian MJ, Yamamoto KR, & Simons SS (1991) Creation of Super Glucocorticoid Receptors by Point Mutations in the Steroid Binding Domain. Journal of Biological Chemistry 266:22075–22078.

54. Chakraborti PK, Garabedian MJ, Yamamoto KR, & Simons SS (1992) Role of Cysteine-640, Cysteine-656, and Cysteine-661 in Steroid Binding to Rat Glucocorticoid Receptors. Journal of Biological Chemistry 267:11366–11373.

55. Birmingham A, et al. (2009) Statistical methods for analysis of high-throughput RNA interference screens. Nat Methods 6:569–575.

56. Zhang XD, et al. (2007) The use of strictly standardized mean difference for hit selection in primary RNA interference high-throughput screening experiments. J Biomol Screen 12:497–509.

57. Gad SC (2012) Development of therapeutic agents handbook (John Wiley & Sons, Hoboken, N.J.) pp xvii, 1258 p.

58. Collinet C, et al. (2010) Systems survey of endocytosis by multiparametric image analysis. Nature 464:243–249.

59. Szklarczyk D, et al. (2017) The STRING database in 2017: quality-controlled protein-protein association networks, made broadly accessible. Nucl Acids Res 45:D362–D368.

60. Laulederkind SJ, et al. (2013) The Rat Genome Database 2013–data, tools and users. Brief Bioinf 14:520–526.

61. Safran M, et al. (2010) GeneCards Version 3: the human gene integrator. Database (Oxford) 2010:baq020.

62. Jiao XL, et al. (2012) DAVID-WS: a stateful web service to facilitate gene/protein list analysis. Bioinformatics 28:1805–1806.

63. Huang DW, Sherman BT, & Lempicki RA (2009) Systematic and integrative analysis of large gene lists using DAVID bioinformatics resources. Nat Protoc 4:44–57.

64. Jackson AL, et al. (2003) Expression profiling reveals off-target gene regulation by RNAi. Nat Biotechnol 21:635–637.

65. Walensky LD, et al. (2004) Activation of apoptosis in vivo by a hydrocarbon-stapled BH3 helix. Science 305:1466–1470.

66. Cromm PM, Spiegel J, & Grossmann TN (2015) Hydrocarbon Stapled Peptides as Modulators of Biological Function. ACS Chem Biol 10:1362–1375.

67. Guerlavais V & Sawyer TK (2014) Advancements in Stapled Peptide Drug Discovery & Development. Ann Rev Med Chem 49:331–345.

68. Li ZF & Blissard G (2015) The vacuolar protein sorting genes in insects: A comparative genome view. Insect Biochem Mol Biol 62:211–225.

69. van der Kant R, et al. (2013) Late endosomal transport and tethering are coupled processes controlled by RILP and the cholesterol sensor ORP1L. Journal of Cell Science 126:3462–3474.

70. van der Kant R, et al. (2015) Characterization of the Mammalian CORVET and HOPS Complexes and Their Modular Restructuring for Endosome Specificity. Journal of Biological Chemistry 290:30280–30290.

71. Pols MS, Brink C, Gosavi P, Oorschot V, & Klumperman J (2013) The HOPS Proteins hVps41 and hVps39 Are Required for Homotypic and Heterotypic Late Endosome Fusion. Traffic 14:219–232.

72. Wartosch L, Gunesdogan U, Graham SC, & Luzio JP (2015) Recruitment of VPS33A to HOPS by VPS16 Is Required for Lysosome Fusion with Endosomes and Autophagosomes. Traffic 16:727–742.

73. Perini ED, Schaefer R, Stoter M, Kalaidzidis Y, & Zerial M (2014) Mammalian CORVET Is Required for Fusion and Conversion of Distinct Early Endosome Subpopulations. Traffic 15:1366–1389.

74. Bright NA, Wartosch L, & Luzio JP (2015) Lysosome fusion in cultured mammalian cells. Meth Cel Biol 126:101–118.

75. D’Agostino M, Risselada HJ, Lurick A, Ungermann C, & Mayer A (2017) A tethering complex drives the terminal stage of SNARE-dependent membrane fusion. Nature.

76. Mattie S, McNally EK, Karim MA, Vali H, & Brett CL (2017) How and why intralumenal membrane fragments form during vacuolar lysosome fusion. Mol Biol Cell 28:309–321.

77. Kweon DH, Kong B, & Shin YK (2017) Hemifusion in Synaptic Vesicle Cycle. Front Mol Neurosci 10.

78. Wang SY, Sun H, Tanowitz M, Liang XH, & Crooke ST (2017) Intra-endosomal trafficking mediated by lysobisphosphatidic acid contributes to intracellular release of phosphorothioate-modified antisense oligonucleotides. Nucl Acids Res 45:5309–5322.

79. Jefferies HBJ, et al. (2008) A selective PIKfyve inhibitor blocks PtdIns(3,5) P(2) production and disrupts endomembrane transport and retroviral budding. EMBO Rep 9:164–170.

80. Savina A, Furlan M, Vidal M, & Colombo MI (2003) Exosome release is regulated by a calcium-dependent mechanism in K562 cells. Journal of Biological Chemistry 278:20083–20090.

81. Willem J, ter Beest M, Scherphof G, & Hoekstra D (1990) A non-exchangeable fluorescent phospholipid analog as a membrane traffic marker of the endocytic pathway. Eur J Cell Biol 53:173–184.

82. Vidal M, Mangeat P, & Hoekstra D (1997) Aggregation reroutes molecules from a recycling to a vesicle-mediated secretion pathway during reticulocyte maturation. Journal of Cell Science 110:1867–1877.

83. Savina A, Vidal M, & Colombo MI (2002) The exosome pathway in K562 cells is regulated by Rab11. Journal of Cell Science 115:2505–2515.

84. Bhatnagar S & Schorey JS (2007) Exosomes released from infected macrophages contain mycobacterium avium glycopeptidolipids and are proinflammatory. Journal of Biological Chemistry 282:25779–25789.

85. Lukinavicius G, et al. (2013) A near-infrared fluorophore for live-cell super-resolution microscopy of cellular proteins. Nat Chem 5:132–139.

86. Futaki S & Nakase I (2017) Cell-Surface Interactions on Arginine-Rich Cell-Penetrating Peptides Allow for Multiplex Modes of Internalization. Accounts Chem Res 50:2449–2456.

87. Madani F, Lindberg S, Langel U, Futaki S, & Graslund A (2011) Mechanisms of cellular uptake of cell-penetrating peptides. J Biophys 2011:414729.

88. Erazo-Oliveras A, Muthukrishnan N, Baker R, Wang TY, & Pellois JP (2012) Improving the endosomal escape of cell-penetrating peptides and their cargos: strategies and challenges. Pharmaceuticals (Basel) 5:1177–1209.

89. Brock R (2014) The uptake of arginine-rich cell-penetrating peptides: putting the puzzle together. Bioconjug Chem 25:863–868.

90. van den Berg A & Dowdy SF (2011) Protein transduction domain delivery of therapeutic macromolecules. Curr Opin Biotechnol 22:888–893.

91. Bechara C & Sagan S (2013) Cell-penetrating peptides: 20 years later, where do we stand? Febs Letters 587:1693–1702.

92. Wender PA, Galliher WC, Goun EA, Jones LR, & Pillow TH (2008) The design of guanidinium-rich transporters and their internalization mechanisms. Adv Drug Deliver Rev 60:452–472.

93. Peraro L & Kritzer J (2018) Getting in: emerging methods and design principles for cell-penetrant peptides. Angew Chem Int Ed Engl.

94. Rabideau AE & Pentelute BL (2016) Delivery of Non-Native Cargo into Mammalian Cells Using Anthrax Lethal Toxin. Acs Chem Biol 11:1490–1501.

95. Thompson DB, Cronican JJ, & Liu DR (2012) Engineering and Identifying Supercharged Proteins for Macromolecule Delivery into Mammalian Cells. Method Enzymol 503:293–319.

96. Kalafatovic D & Giralt E (2017) Cell-Penetrating Peptides: Design Strategies beyond Primary Structure and Amphipathicity. Molecules 22.

97. Daniels DS & Schepartz A (2007) Intrinsically cell-permeable miniature proteins based on a minimal cationic PPII motif. J Am Chem Soc 129:14578–+.

98. Smith BA & Schepartz A (2009) Understanding cell uptake by minimally cationic cell-permeable miniature proteins. Abstr Pap Am Chem S 238:818–818.

99. Millet JK & Whittaker GR (2017) Physiological and molecular triggers for SARS-CoV membrane fusion and entry into host cells. Virology.

100. Burkard C, et al. (2014) Coronavirus Cell Entry Occurs through the Endo-/Lysosomal Pathway in a Proteolysis-Dependent Manner. Plos Pathogens 10.

101. Carette JE, et al. (2011) Ebola virus entry requires the cholesterol transporter Niemann-Pick C1. Nature 477:340–U115.

102. Li B, et al. (2017) The FgVps39-FgVam7-FgSso1 Complex Mediates Vesicle Trafficking and Is Important for the Development and Virulence of Fusarium graminearum. Mol Plant-Microbe Int 30:410–422.

103. Palmer GE, Cashmore A, & Sturtevant J (2003) Candida albicans VPS11 is required for vacuole biogenesis and germ tube formation. E Cell 2:411–421.

104. Palmer GE, Kelly MN, & Sturtevant JE (2005) The Candida albicans vacuole is required for differentiation and efficient macrophage killing. E Cell 4:1677–1686.

105. Liu X, Hu G, Panepinto J, & Williamson PR (2006) Role of a VPS41 homologue in starvation response, intracellular survival and virulence of Cryptococcus neoformans. Mol Microbiol 61:1132–1146.

106. Geissenhoner A, Sievers N, Brock M, & Fischer R (2001) Aspergillus nidulans DigA, a potential homolog of Saccharomyces cerevisiae Pep3 (Vps18), is required for nuclear migration, mitochondrial morphology and polarized growth. Mol Genet Gen 266:672–685.

107. Yang ST, Zaitseva E, Chernomordik LV, & Melikov K (2010) Cell-penetrating peptide induces leaky fusion of liposomes containing late endosome-specific anionic lipid. Biophys J 99:2525–2533.

108. Kobayashi T, et al. (2002) Separation and characterization of late endosomal membrane domains. Journal of Biological Chemistry 277:32157–32164.

109. Kobayashi T, et al. (1999) Late endosomal membranes rich in lysobisphosphatidic acid regulate cholesterol transport. Nat Cell Biol 1:113–118.

110. Bissig C & Gruenberg J (2014) ALIX and the multivesicular endosome: ALIX in Wonderland. Trends Cell Biol 24:19–25.

111. Raiborg C & Stenmark H (2009) The ESCRT machinery in endosomal sorting of ubiquitylated membrane proteins. Nature 458:445–452.

112. Falguieres T, et al. (2008) In Vitro Budding of Intralumenal Vesicles into Late Endosomes Is Regulated by Alix and Tsg101. Mol Biol Cell 19:4942–4955.

113. Matsuo H, et al. (2004) Role of LBPA and Alix in multivesicular liposome formation and endosome organization. Science 303:531–534.

## References

1. Lukinavicius G, et al. (2013) A near-infrared fluorophore for live-cell super-resolution microscopy of cellular proteins. Nat Chem 5:132–139.

2. Campeau E, et al. (2009) A Versatile Viral System for Expression and Depletion of Proteins in Mammalian Cells. Plos One 4.

3. Maejima I, et al. (2013) Autophagy sequesters damaged lysosomes to control lysosomal biogenesis and kidney injury. EMBO J 32:2336–2347.

4. Holub JM, Larochelle JR, Appelbaum JS, & Schepartz A (2013) Improved assays for determining the cytosolic access of peptides, proteins, and their mimetics. Biochemistry 52:9036–9046.

5. LaRochelle JR, Cobb GB, Steinauer A, Rhoades E, & Schepartz A (2015) Fluorescence correlation spectroscopy reveals highly efficient cytosolic delivery of certain penta-arg proteins and stapled peptides. J Am Chem Soc 137:2536–2541.

6. Sinclair JK, Denton EV, & Schepartz A (2014) Inhibiting epidermal growth factor receptor at a distance. J Am Chem Soc 136:11232–11235.

7. Qian Z, et al. (2016) Discovery and Mechanism of Highly Efficient Cyclic Cell-Penetrating Peptides. Biochemistry 55:2601–2612.

8. Yu P, Liu B, & Kodadek T (2005) A high-throughput assay for assessing the cell permeability of combinatorial libraries. Nat Biotechnol 23:746–751.

9. Mallik B, et al. (2005) Design and NMR characterization of active analogues of compstatin containing non-natural amino acids. J Med Chem 48:274–286.

10. Quach K, LaRochelle J, Li XH, Rhoades E, & Schepartz A (2017) Unique arginine array improves cytosolic localization of hydrocarbon-stapled peptides. Bioorg Med Chem.

11. Nitsche JM, Chang HC, Weber PA, & Nicholson BJ (2004) A transient diffusion model yields unitary gap junctional permeabilities from images of cell-to-cell fluorescent dye transfer between Xenopus oocytes. Biophys J 86:2058–2077.

12. Weiss M, Hashimoto H, & Nilsson T (2003) Anomalous protein diffusion in living cells as seen by fluorescence correlation spectroscopy. Biophys J 84:4043–4052.

13. Bronstein I, et al. (2009) Transient anomalous diffusion of telomeres in the nucleus of mammalian cells. Phys Rev Lett 103:018102.

14. Schneider CA, Rasband WS, & Eliceiri KW (2012) NIH Image to ImageJ: 25 years of image analysis. Nat Meth 9:671–675.

15. Blangy A, et al. (1995) Phosphorylation by p34(cdc2) regulates spindle association of human Eg5, a kinesin-related motor essential for bipolar spindle formation in vivo. Cell 83:1159–1169.

16. Malo N, Hanley JA, Cerquozzi S, Pelletier J, & Nadon R (2006) Statistical practice in high-throughput screening data analysis. Nature Biotechnology 24:167–175.

17. Zhang XD, et al. (2007) The use of strictly standardized mean difference for hit selection in primary RNA interference high-throughput screening experiments. J Biomol Screen 12:497–509.

18. VanNoorden CJF, et al. (1997) Ala-pro-cresyl violet, a synthetic fluorogenic substrate for the analysis of kinetic parameters of dipeptidyl peptidase IV (CD26) in individual living rat hepatocytes. Analytical Biochemistry 252:71–77.

19. Dunn KW, Kamocka MM, & McDonald JH (2011) A practical guide to evaluating colocalization in biological microscopy. Am J Physiol-Cell Ph 300:C723–C742.

